# Life at the extremes: Maximally divergent microbes with similar genomic signatures linked to extreme environments

**DOI:** 10.1101/2025.06.04.657665

**Authors:** Monireh Safari, Joseph Butler, Gurjit S. Randhawa, Kathleen A. Hill, Lila Kari

## Abstract

Extreme environments impose strong mutation and selection pressures that drive distinctive, yet understudied, genomic adaptations in extremophiles. In this study, we identify 15 bacterium–archaeon pairs that exhibit highly similar *k*-mer–based genomic signatures despite maximal taxonomic divergence, suggesting that shared environmental conditions can produce convergent, genome-wide patterns that transcend evolutionary distance. To uncover these patterns, we developed a computational pipeline to select a composite genome proxy assembled from non-contiguous subsequences of the genome. Using supervised machine learning on a curated dataset of 693 extremophile microbial genomes, we found that 6-mers and 100 kbp genome proxy lengths provide the best balance between classification accuracy and computational efficiency. Our results provide conclusive evidence of the pervasive nature of *k*-mer–based patterns across the genome, and uncover the presence of taxonomic and environmental components that persist across all regions of the genome. The 15 bacterium-archaeon pairs identified by our method as having similar genomic signatures were validated through multiple independent analyses, including 3-mer frequency profile comparisons, phenotypic trait similarity, and geographic co-occurrence data. These complementary validations confirmed that extreme environmental pressures can override traditionally recognized taxonomic components at the whole-genome level. Together, these findings reveal that adaptation to extreme conditions can carry robust, taxonomic domain-spanning imprints on microbial genomes, offering new insight into the relationship between environmental mutagenesis and selection and genome-wide evolutionary convergence.

## 1 Introduction

The study of extremophiles, organisms capable of thriving in Earth’s most extreme environments, provides crucial insights into genetic adaptations for survival under harsh conditions. These organisms have evolved to withstand extreme environmental conditions such as high temperature, pH, pressure, salinity, and radiation levels and, notably, they are often unable to survive outside of these physiologically extreme environmental conditions [60, 46]. Recent attention to microbial extremophiles has highlighted their value across diverse applications, including biotechnology, particularly in biorefineries and as sources of industrial extremozymes for high-temperature or high-pH processes [69, 75, 81, 18, 38], environmental bioremediation of metal-contaminated, saline, or radioactive environments [18], as well as agriculture and soil enhancement [52], and veterinary medicine [23]. Additionally, interest has grown in the ability of microbial extremophiles to survive the extreme conditions of outer space [49, 21].

Extremophiles have evolved specific genomic and proteomic adaptations to survive in environments with extreme conditions [17, 34]. At the genomic level, they frequently exhibit gene duplications [74, 57] and reduced genome sizes, particularly in thermophilic species [72]. The nucleotide composition of these organisms displays environment-specific patterns in G+C content and purine load [76, 31, 57]. Multiple molecular mechanisms, including gene duplication and horizontal gene transfer [61, 9, 56], also contribute to the genomic adaptation of extremophiles. Recent research has further emphasized the critical role of genomic regulatory elements in these environmental adaptations [63, 48, 18], as well as efficient DNA repair systems in extremophiles exposed to high radiation [38]. At the proteomic level, features such as codon usage bias and amino acid composition have been linked to thermal adaptation and have recently been used in machine learning models to predict optimal growth temperature [10].

A recent study applied supervised and unsupervised machine learning algorithms to explore the genomes of microbial extremophiles, uncovering both taxonomic and environmental components embedded within their genomic signatures [2]. In this approach, genomic signatures were derived from 500 kbp (contiguous) representative DNA fragments randomly selected from each genome, by computing the *k*-mer frequency vector of each fragment. Here, a *k*-mer is a DNA sequence of length *k*, and the *k*-mer frequency vector of a DNA fragment is a numerical vector comprising the counts of the occurrences of all possible *k*-mers in that fragment (in lexicographic order). These vectors enabled highly accurate classification and clustering tasks that revealed environmental components for extremophiles inhabiting environments with extreme temperature and/or pH conditions. While this approach provided valuable insights, it had some notable limitations. Firstly, the selection of the DNA fragment selected to act as a genome proxy was not entirely random (see Section 2.3), which could potentially introduce bias in the derived genomic signatures. Secondly, the study did not quantitatively test the hypothesis of the pervasiveness of the environmental components across the entire genome.

This paper addresses these limitations by first refining the process of selecting a genomic signature to enhance the accuracy of organism classification based on genomic data. We focus on three key areas: *(i)* improving the genome coverage and randomness of the representative DNA fragment by replacing it with a *composite genome proxy* constructed through the pseudo-concatenation of several randomly selected (non-contiguous) DNA fragments, *(ii)* comprehensively testing the hypothesis of the pervasiveness of taxonomic and environmental components across a genome, and *(iii)* analyzing the impact of varying *k*-mer sizes and composite genome proxy lengths, ranging from 10 kbp to entire genomes. Through a series of computational experiments involving several genome proxy selection methods, *k*-mer sizes, and fragment lengths, we aimed to identify the optimal parameters for extremophile genomic signature analysis. The results of these analyses were then used to design a multilayered pipeline that identified multiple bacterium–archaeon pairs with similar genomic signatures in spite of their maximal taxonomic divergence, potentially due to the shared characteristics of their extreme environments. The main contributions of this paper are:

1. Conclusive evidence of the pervasiveness of a *k*-mer-based genomic signature throughout an extremophile genome.
2. A broadly applicable method for the selection of a composite genome proxy (hereafter referred to simply as “genome proxy”) assembled from non-contiguous subsequences of the genome. Empirical determination of the optimal *k*-mer size (*k* = 6) and genome proxy length (100, 000 bp) for fast and accurate taxonomic and environment-type classifications.
3. Discovery of 15 maximally divergent bacterium-archaeon pairs with similar genomic signatures linked to the characteristics of their extreme environment, through a multi-layered filtering process used in conjunction with unsupervised machine learning.
4. Validation of the above computational findings through additional analyses, including 3-mer frequency profile analyses demonstrating agreement with known adaptative patterns in extremophiles, statistical confirmation of 3-mer frequency profiles similarity of the identified pairs using Spearman’s rank correlation analysis [66], and analysis of geographic co-occurrence data confirming that identified bacterium-archaeon pairs naturally co-occur in same extreme environments.

## 2 Materials and Methods

This section describes the methodology employed in this study’s computational experiments: Section 2.1 provides a detailed description of the genome sequence datasets utilized in this study; Section 2.2 provides an explanation of genomic signatures, Section 2.3 outlines the procedure employed for selecting a genome proxy to represent a genome for the purpose of taxonomic and environment-type based machine learning classifications, and the methods used to empirically optimize the *k*-mer value and genome proxy length; Section 2.4 describes the multilayer process utilized to discover microbes that share environmental genomic component in spite of belonging to different taxa of maximal evolutionary divergence.

### 2.1 Dataset

In the quest to evaluate genome-wide genomic signatures potentially shaped by similar extreme environments for maximally divergent microbes, it is essential to determine the optimal genome proxy to represent each genome. This selection is crucial for ensuring accurate *k*-mer-based classification and clustering.

For this reason, and in order to be able to perform apples-to-apples comparisons with existing results, we utilized the dataset from [2]. The dataset consists of 693 high-quality extremophile microbial genome assemblies curated via a comprehensive review of primary literature and cross-referenced with the Genome Taxonomy Database [51]. These microbial genomes were grouped into two different environment-type datasets, one based on the organisms’ optimal growth temperature (psychrophiles, mesophiles, thermophiles, hyperthermophiles, see Table 1) and the other based on their optimal growth pH levels (acidophiles and alkaliphiles, see Table 2).

**Table 1:** The *Temperature Dataset*: Taxonomic diversity of archaea and bacteria across temperature categories. The four temperature categories are defined, based on the optimal temperature for growth (OTG). These categories are as follows: Psychrophiles (OTG of *<* 20^◦^C), Mesophiles (OTG of 20−45^◦^C), Thermophiles (OTG of 45−80^◦^C), and Hyperthermophiles (OTG of *>* 80^◦^C) [2].

| Domain | Temperature Category | # Phyla | # Classes | # Orders | # Families | # Genera | # Species |
| --- | --- | --- | --- | --- | --- | --- | --- |
| Archaea | Psychrophiles | 2 | 4 | 4 | 5 | 7 | 8 |
|  | Mesophiles | 4 | 6 | 7 | 20 | 45 | 84 |
|  | Thermophiles | 6 | 11 | 14 | 21 | 41 | 67 |
|  | Hyperthermophiles | 5 | 6 | 8 | 15 | 31 | 70 |
| Bacteria | Psychrophiles | 4 | 4 | 6 | 13 | 19 | 140 |
|  | Mesophiles | 3 | 3 | 6 | 10 | 14 | 106 |
|  | Thermophiles | 15 | 19 | 24 | 27 | 47 | 116 |
|  | Hyperthermophiles | 5 | 5 | 5 | 5 | 5 | 7 |

**Table 2:**
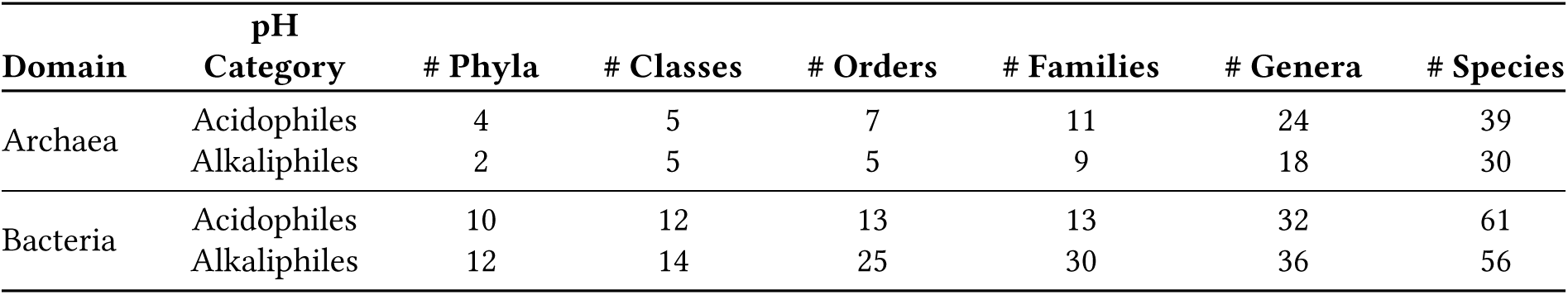
The *pH Dataset*: Taxonomic diversity of archaea and bacteria across pH categories. . The two pH categories are defined based on the optimal growth pH (OGpH). These are: Acidophiles (OGpH *<* pH 5) and Alkaliphiles (OGpH *>* pH 9) [2].

| Domain | pH Category | # Phyla | # Classes | # Orders | # Families | # Genera | # Species |
| --- | --- | --- | --- | --- | --- | --- | --- |
| Archaea | Acidophiles | 4 | 5 | 7 | 11 | 24 | 39 |
|  | Alkaliphiles | 2 | 5 | 5 | 9 | 18 | 30 |
| Bacteria | Acidophiles | 10 | 12 | 13 | 13 | 32 | 61 |
|  | Alkaliphiles | 12 | 14 | 25 | 30 | 36 | 56 |

The first dataset, called the *Temperature Dataset*, is composed of 598 genomes including 148 psychrophile genomes, 190 mesophile genomes, 183 thermophile genomes, and 77 hyperthermophile genomes. The second dataset, called the *pH Dataset*, is composed of 186 genomes, including 100 acidophile genomes and 86 alkaliphile genomes. There are 91 genomes that are present in both datasets, falling into one of two categories: mesophiles that live in acidic or alkaline environments (8 genomes), and polyextremophiles, that thrive in environments that are both acidic/alkaline and at extreme temperatures (83 genomes). The details of the samples that are in both datasets can be found in the Supplementary Materials, Section A.

### 2.2 Genomic signature

Due to the massive lengths of genomic sequences and the high computational demands of alignment-based methods [82], researchers are now using alignment-free methods leveraging “genomic signatures” for efficient genome classification or clustering. The genomic signature of an organism is typically represented by a *k*-mer frequency vector [13], derived from the entire genome or a “sufficiently long” DNA fragment that captures the pervasiveness of the signature [29]. These signatures have proven effective in differentiating species and have been applied in various contexts, including microbial diversity analysis [44, 4, 54, 77, 35, 5, 45], classification or subtyping of viral genomes [65, 15, 54, 55, 1, 50], and metagenomic classification and profiling [70, 35].

Chaos Game Representation (CGR) of DNA sequences, first introduced by Jeffrey in 1990 [25] has emerged as a particularly effective method for calculating and visualizing genomic signatures. Figure 1, left panel, provides a brief illustration of the process of generating the CGR of the sample DNA sequence “ACG.” A quantified version of CGR, called Frequency Chaos Game Representation (FCGR) [14], produces a 2^*k*^ × 2^*k*^ grayscale image, where the pixel intensities correspond to *k*-mer frequencies. The patterns in an FCGR of a genomic sequence reflect its composition, and several studies have demonstrated the effectiveness of FCGR images in taxonomic classification at various taxonomic levels [62, 80, 59]. As expected, the FCGRs of an archaeon and a bacterial species are visually different, which is consistent with the genetic difference anticipated for species of two different domains of life (Figure 1, right panel).

**Figure 1:**
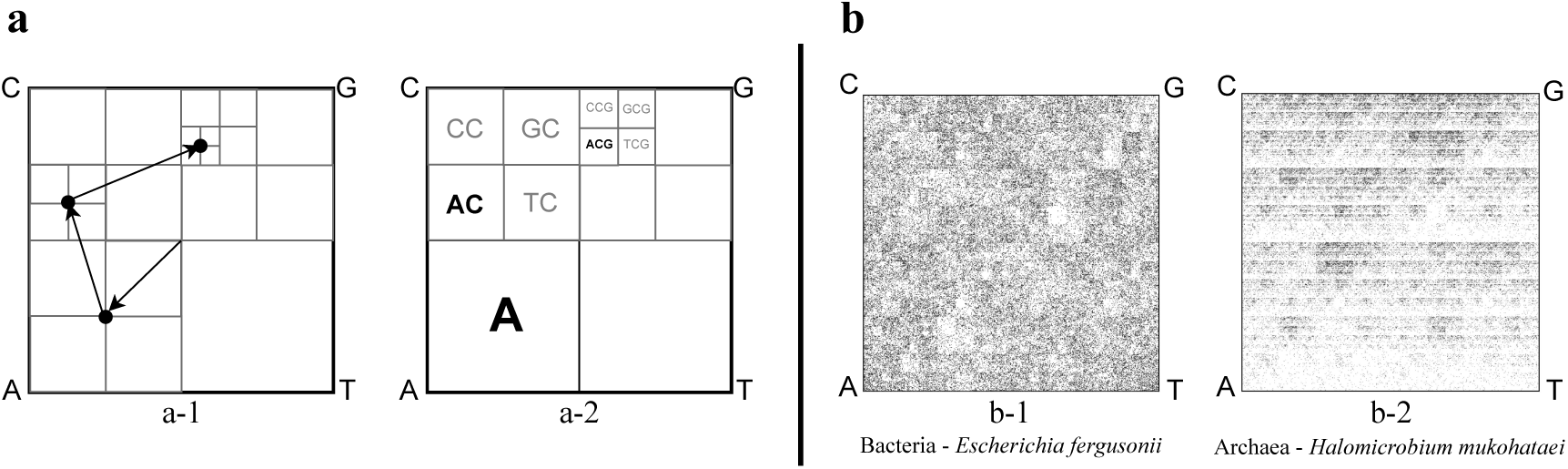
*Left:* Chaos Game Representation (CGR) of the DNA sequence “ACG.” *a-1:* The CGR is generated within a square with corners labelled A, C, G, T. The plot is generated by reading the sequence from left to right, and iteratively plotting the midpoint between the current point and the corner labelled by the nucleotide being read (the start point is the square’s center). For example, the sequence ACG consists of three points generated in the order illustrated by the arrows. *a-2:* The resulting CGR, where the square regions labelled by **A**, **AC**, and **ACG** are the regions where the *k*-mers A, AC, ACG would be plotted, regardless of their position in the sequence. *Right:* FCGRs of randomly selected 100 kbp genomic fragments belonging to organisms from two different domains of life, *b-1: E.fergusonii* (bacterium) and *b-2: H.mukohataei* (archaeon). Both species live in similar habitats with moderate temperatures (optimal growth temperature of 20 − 45^◦^C). The value of *k* is 8, and thus each image is a 256 × 256 grayscale grid (2^8^ = 256), where each pixel represents the frequency of a specific 8-mer in the DNA fragment. Darker (lighter) pixels indicate higher (lower) numbers of occurrences of the corresponding 8-mers in the respective sequences. The unique patterns in each image reflect the genome sequence composition for that species.

The capability of genomic signatures to differentiate between organisms across taxonomic levels, combined with the visualization power of FCGRs, makes genomic signatures particularly suitable for the analysis of extremophiles, where we seek to understand how environmental adaptations might influence genomic signatures across different taxa.

### 2.3 Selecting the genome proxy, and empirically optimizing the ***k***-mer value and genome proxy length

This section begins by proposing a new procedure for the selection of a genome proxy (Section 2.3.1). Using this selection method, along with supervised learning methods applied to the *Temperature Dataset* and *pH Dataset*, we then assessed the effect of genome proxy selection on classification accuracy, and empirically determined the optimal values for *k*-mer size and representative genome proxy length (Section 2.3.2).

#### 2.3.1 Selecting a genome proxy

While genomic signatures have been shown to be effective for classification and clustering of genomic sequences, the validity and accuracy of such analysis highly depend on the selection of representative DNA fragments capable of serving as a genome proxy. The approach utilized in [2] had notable limitations that our current methodology aims to address. Specifically, the selection process was not completely random, since the selected representative was a contiguous long fragment of the genome and the selection process prioritized longer contigs over shorter ones. Recall that the selection process in [2] starts from the list of contigs sorted in decreasing order of their length. If the longest contig exceeded 500 kbp in length, a 500 kbp subfragment was randomly chosen from that contig as the representative DNA fragment of that genome. Otherwise, the contigs were pseudo-concatenated one after another, until the pseudo-concatenated sequence reached a length of 500 kbp, and this sequence was taken to be the representative DNA fragment of that genome. Here, the *pseudo-concatenation* of DNA sequences is defined as listing them one after another, with a separator letter ‘N’ between every two consecutive sequences. Pseudo-concatenation prevents the formation of spurious *k*-mers during the process, and *k*-mers containing the letter ‘N’ are not counted when computing the *k*-mer frequency vector of the pseudo-concatenated sequence, with ‘N’ not being counted towards the pseudo-concatenated sequence length. One potential limitation of this selection process is that it biases the choice towards representative DNA fragments extracted from longer contigs. Another potential limitation is that, if the first contig is sufficiently large, the representative fragment will be selected from a single region of the genome. These limitations introduce a bias in the selection of the representative DNA fragment, which presupposed a genome-wide pervasive nature of a genomic signature and could potentially affect the classification accuracy.

To address these limitations we propose a procedure for selecting a composite genome proxy that ensures that all fragments in the genome have an equal probability of being included in the final genome proxy. In addition, this procedure ensures that the final genome proxy includes multiple sequences from various locations in the genome. The method of selecting a genome proxy has three steps:

1. The optimal values for *n* (the number of non-overlapping genomic subfragments that comprise a genome proxy *s*), and for *len*(*s*) (the total length of the genome proxy *s*) are empirically determined.
2. If the genome sequence is composed of multiple contigs, all contigs are pseudo-concatenated into a single large sequence;
3. *n* different, randomly selected, non-overlapping subfragments of length *len*(*s*)/*n* are pseudo-concatenated from either the genome (if it consists of a single contig), or from the pseudo-concatenated sequence obtained in the preceding step (if the genome consists of multiple contigs).

Figure 2 illustrates the selection process of a genome proxy when the genome consists of only one contig, *n* = 3, and *len*(*s*) = 15.

**Figure 2:**
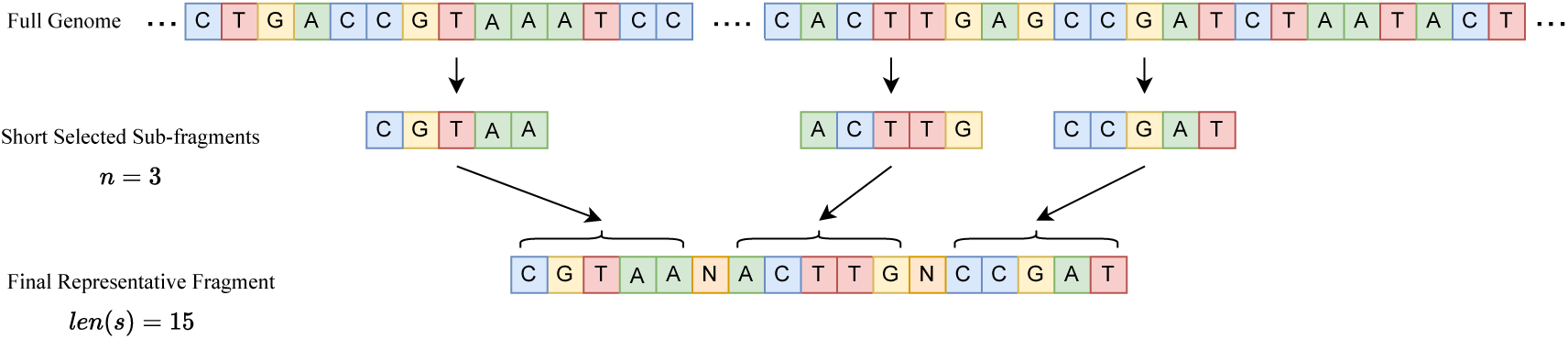
The selection process of a genome proxy *s*, comprising *n* = 3 non-overlapping sub-fragments, and with total length *len*(*s*) = 15. Top: Full genome, consisting of only one contig. Middle: *n* non-overlapping subfragments (here *n* = 3) randomly selected from the genome. Bottom: The genome proxy of length *len*(*s*) = 15 obtained by pseudo-concatenating the sub-fragments.

#### 2.3.2 Determining the optimal ***k***-mer size and genome proxy length

After selecting the genome proxy in a way that ensured randomness, we evaluated the impact of the following three factors on the supervised classifier accuracies *(i)* the choice of genome proxy, *(ii) k*-mer size, and *(iii)* genome proxy length. The feature vectors used as an input for these classifiers were *canonical k-mer* frequency vectors. Here, a “canonical *k*-mer” is defined as the first, in alphabetical order, of a *k*-mer and its Watson-Crick complement. For any DNA sequence, the final frequency vector was computed by averaging the *k*-mer counts of the sequence with *k*-mer counts of its Watson-Crick reverse complement [2]. In the remainder of this paper, only the canonical *k*-mer will be listed.

The classifier used for step *(i)* was the Support Vector Machine (SVM) with a Radial Basis Function (RBF) kernel, which has been shown to achieve high accuracies in genome sequence classification across various fields [2, 22, 79]. For steps *(ii)* and *(iii)*, we expanded our analysis to include six classifiers: SVM with RBF kernel, Random Forest with 100 estimators, and an Artificial Neural Network (ANN) with two hidden layers (sizes 256 and 64, and a learning rate of 0.001). Additionally, three variations of the Machine Learning with Digital Signal Processing (MLDSP) algorithm (MLDSP-1, MLDSP-2, and MLDSP-3)[54] were also included.

In these computational experiments, we explored nine *k*-mer sizes ranging from 1 to 9, to find the optimal value of *k*. Larger *k*-mer sizes were avoided to prevent sparsity in the feature vector, which can undermine classification accuracy. Six genome proxy lengths (*len*(*s*)) were evaluated: 10 kbp, 50 kbp, 100 kbp, 250 kbp, 500 kbp, and 1,000 kbp. The fragment lengths were chosen to include both short sequences (e.g., 10 kbp, approximately 3% of the average sequence length in our dataset) and longer sequences, allowing for a comprehensive comparison while also considering computational costs. For all experiments, the number of sub-fragments (*n*) comprising the sequence *s* was set to 10, a value empirically determined as optimal for the datasets used. This choice is supported by existing studies indicating that the minimum sequence length necessary to capture genomic patterns is of the order of 10^3^ [25], making *n* = 10 an effective choice for ensuring that each sub-fragment independently captures the relevant genomic patterns, particularly in the case of shorter fragments.

To implement step *(i)*, we repeated the following process 10 times: First, a random genome proxy was selected for each sequence. Then, for each combination of *k*-mer size and fragment length (9×6 combinations), we performed classification using an SVM with 10-fold cross-validation. To assess the performance of the classification, the accuracy was defined as the ratio of the number of sequences with correctly predicted labels to the total number of sequences classified. The variance of the classification accuracy over these ten runs of the classification was then calculated to determine if the accuracy was dependent on the choice of random genome proxy.

Following the observation that classification accuracy is not dependent on the choice of genome proxy, steps *(ii)*, finding the optimal *k*-mer size, and *(iii)*, finding the optimal fragment length, were carried out by running classification experiments using the six classifiers with all combinations of *k*-mer sizes and fragment lengths. Note that, for each fragment length, a fixed randomly selected genome proxy was utilized.

In addition, a separate experiment was performed using the full DNA genome (maximal sequence length) and the optimal *k*-mer size determined in step *(ii)*, to evaluate the impact of considering the information from the whole genome on classification accuracy, as opposed to a random shorter fragment.

The computational experiments were performed for the two different datasets, the *Temperature Dataset* and the *pH Dataset*. All classifications were conducted under two distinct supervised training scenarios: genome proxies labelled with taxonomic labels (bacteria or archaea), and genome proxies labelled with environment-type labels (for the *Temperature Dataset*, psychrophiles, mesophiles, thermophiles, and hyperthermophiles; for the *pH Dataset*, acidophiles and alkaliphiles).

The analysis used standard stratified 10-fold cross-validation. For the environment-type classification, the tests were performed under two different labelling scenarios, called *restriction-free* scenario, and *restricted* (also called *non-overlapping genera*) scenario, respectively. In the restriction-free scenario, we utilized standard stratified 10-fold cross-validation and reported the average classification accuracy for each of the ten test folds. The restricted scenario aimed to untangle the taxonomic (genus level) component from the environmental component in the genomic signatures [2]. To this end, the restricted scenario test used an adjusted 10-fold cross-validation approach, where all sequences belonging to the same genus appeared in the same fold, while the distribution of labels in each fold remained the same as in the entire dataset.

### 2.4 Finding bacterium–archaeon pairs with similar genomic signatures, linked to their extreme environments

Once the performance of the optimal parameters was validated, the main objective of this study was to identify, if any microbe pairs from two different taxonomic domains (archaea and bacteria) shared similarities in their genomic signatures that were linked to their shared extreme environment types.

A multi-layer approach was used, where the first layer was the generation of “candidate bacterium–archaeon pairs,” i.e., maximally different microbe pairs clustered together by non-parametric clustering machine learning algorithms (Section 2.4.1). To eliminate potential clustering algorithm errors, this candidate pair list was subjected to a second selection layer, comprising two additional criteria: FCGR comparison (Section 2.4.2) and metadata analysis of the environment of the organism isolation (Section 2.4.3). After finalizing the list of bacterium–archaeon pairs, the 3-mer frequency profiles of these pairs were analyzed, and corroboration with biological findings regarding over– and under-representations of 3-mers was assessed in the genes and genomes of extremophiles living in certain extreme environments (Section 2.4.4). In the last step, we also studied the geographic habitat co-occurrence of these final bacterium–archaeon pairs (Section 2.4.5).

#### 2.4.1 Non-parametric clustering

In this study, only non-parametric clustering algorithms were used, since they have the advantage of not needing the expected number of clusters as an input parameter. Specifically, the five algorithms used were the non-parametric version of the *i*DeLUCS algorithm [44], and four other non-parametric algorithms (HDBSCAN [41], Affinity Propagation [16], MeanShift [11] and iterative medoids [47]). These algorithms were applied in conjunction with two different dimensionality reduction techniques, Variational Autoencoders (VAE) [47], and Uniform Manifold Approximation and Projection (UMAP) [42]. The primary objective was to identify the optimal combinations of dimensionality reduction techniques and clustering algorithms that effectively reproduce clusters corresponding to true genera in our datasets. To achieve this, we evaluated the performance of each combination by calculating the *completeness* and *contamination* of the resulting clusters. Completeness refers to the proportion of true members within a cluster (cluster members belonging to the same genus) relative to the total cluster size, and contamination indicates the proportion of incorrect members (cluster members that belong to a different genus) relative to the total cluster size. Only those clusters were accepted as “genus-accurate” that had completeness greater than 50% and contamination less than 50%. The next step was to rank the aforementioned combinations by the ratio of the number of genus-accurate clusters to the total number of generated clusters. The top five combinations were selected, namely: VAE+iterative medoids (IM), VAE+ Affinity Propagation, VAE+HDBSCAN, UMAP+HDBSCAN, and *i*DeLUCS. In the final step, we used all output clusters from the selected top five combinations to identify pairs of archaea and bacteria that clustered together. Specifically, for each of the top five combinations, we ran the clustering process 10 times, each time with a different random seed, each time producing the pairs of maximally divergent microbes that were clustered together. From the resulting set of pairs, the pairs that appeared in more than five runs, and were clustered together by the majority of the five combinations, were retained, as being “candidate bacterium–archaeon pairs,” subjected to the next layer of analysis.

#### 2.4.2 FCGR comparison of candidate pairs

To address the errors inherent in any unsupervised clustering method, we then analyzed the “candidate bacterium–archaeon pairs” identified in Section 2.4.1. The first step involves investigating the similarities of the FCGR patterns of the members of each candidate pair. For this analysis, FCGR images of candidate pairs were generated from the selected genome proxy using the optimal *k*-mer size determined previously. Subsequently, three distinct distance metrics were used, Descriptor [28], structural dissimilarity index measure (DSSIM) [73], and learned perceptual image patch similarity (LPIPS) [78] to calculate the distances between each pair of candidate bacterium–archaeon pairs that were clustered together. The refined set of candidate pairs was selected based on the similarity of their FCGR images. Specifically, pairs were selected if the distance between their FCGRs was below certain distance-dependent thresholds for at least two out of the three distance metrics. The distance-dependent thresholds were 0.2 for the Descriptor distance, 0.5 for DSSIM, and 0.5 for LPIPS, and were empirically determined as detailed below.

The thresholds for the distance metrics were determined based on the idea that two members of a bacterium–archaeon pair can be considered similar if their FCGR distance is less than the distance among FCGRs of species of the same genus. To this end, the maximum intra-genus distance in the dataset was computed as follows. First, we selected all unique genera from both the *Temperature dataset* and *pH dataset*, excluding those with only a single sample, which resulted in 92 unique genera. Then, for each genus, the pairwise distances between the FCGRs of all sequences were calculated. The average of these distances within each genus was deemed to be the intra-genus distance for that genus. Of the obtained intra-genus distances, 5% of the distances were excluded as outliers (the top and bottom 2.5%). In the final step, the maximum of these average intra-genus distances was considered as the empirical threshold for FCGR comparison for that distance. More details of intra-genus distance computations can be found in Supplementary Materials, Section B. This approach ensured that the identified microbial pairs clustered together based on genomic signature similarity in the previous layer, also exhibited significant similarities in their FCGR patterns. The output of this layer was a list of “FCGR-accepted candidate pairs”.

#### 2.4.3 Comparisons of environmental labelling and phenotypic features for candidate pairs

In parallel with FCGR analysis, “candidate bacterium–archaeon pairs” identified through non-parametric clustering were further investigated in the context of the environmental conditions of their original habitats. The first step involved comparing the environmental labels assigned to each species within a pair (e.g., temperature and pH). Microbial pairs with matching (implying the same temperature and/or pH labels, i.e. both species are acidophiles) or nearly matching (similar temperature and/or pH labels, i.e. both species inhabit high-temperature environments, though one is thermophilic and the other is hyperthermophilic) environmental labels were considered as “environmentally-accepted candidate pairs” and considered for further analysis. For these pairs, we conducted a more detailed analysis. We retrieved the original studies that first characterized these microbes from PubMed (https://pubmed.ncbi.nlm.nih.gov/). The growth parameters and environmental metadata, such as optimal pH and temperature ranges, were compared across species. Additionally, we examined phenotypic traits and habitat-specific characteristics to gain a deeper understanding of shared environmental adaptations and similar phenotype features of the pairs. Finally, the common pairs found in both “FCGR-accepted candidate pairs” and the “environmentally-accepted candidate pairs” were considered “confirmed bacterium–archaeon pairs”, which were further categorized into five groups based on their environmental labels.

#### 2.4.4 Analysis of **3**-mer frequency profiles of confirmed bacterium–archaeon pairs

Following the refinement steps, we conducted a comprehensive 3-mer usage bias analysis by comparing the 3-mer frequency profiles of the “confirmed bacterium–archaeon pairs”. We selected *k* = 3 for this analysis because this *k*-mer length effectively captures codon usage bias, amino acid bias, and protein-associated phenotypic adaptations [2, 8, 3]. Our analysis consisted of four main components.

First, for each 3-mer, we calculated its average frequency across all samples in the *Temperature Dataset* and *pH Dataset*, then calculated the deviation of the 3-mer frequency of each member of the “confirmed bacterium–archaeon pairs” from its dataset average. This approach revealed patterns of 3-mer over– and under-representation in candidates compared to the entire dataset, allowing us to investigate whether similar environmental conditions induced comparable patterns of 3-mer usage across microbial pairs.

Second, we tested the correlation between the 3-mer counts of members of the confirmed pair in each group using Spearman’s rank correlation coefficient, a nonparametric measure of the strength and direction of association between two variables measured on an ordinal scale [66, 55, 7, 30, 20]. This step investigated the pairwise correlation of 3-mer representation among confirmed pairs, providing a *p*-value to assess the significance of similarity or dissimilarity in the 3-mer over– and under-representation.

Third, we identified the specific 3-mers that influenced environmental label prediction in supervised classification for each microbial species in the “confirmed bacterium–archaeon pairs” set. We used the SHapley Additive exPlanations (SHAP) [36] feature importance method to quantify each 3-mer’s contribution to the model’s environmental classification decisions. We referred to these 3-mers as “environment-relevant 3-mers” due to their impact on the model’s ability to distinguish between sequences belonging to organisms living in different environmental conditions.

Finally, we treated the “environment-relevant 3-mers” as quasi-codons and translated them to corresponding amino acids. This translation step enabled direct comparisons between the environment-relevant 3-mers discovered by our method and both codon and amino acid usage biases previously reported in the literature for the respective extremophilic groups. This comparative analysis serves to validate our methodology by demonstrating that the 3-mers we identified as important for environmental-based classification align with known adaptive patterns in extremophiles reported in the literature [2].

#### 2.4.5 Geographic habitat co-occurrence analysis of confirmed bacterium–archaeon pairs

In this analysis, the Microbe Atlas Project (MAP) database [39], cataloging 16S rRNA reads of microbes isolated from a wide range of environments, was used to analyze the geographic habitat co-occurrence of species in “confirmed bacterium–archaeon pairs”. 16S rRNA is a gene encoding a ribosomal subunit highly conserved between different prokaryotes (including bacteria and archaea) [26]. The sequencing of this gene permits highly sensitive taxonomic classification/identification of prokaryotic samples, proving extremely helpful in identifying species found in diverse microbiomes. The MAP tool compiles millions of samples isolated across the world, along with their taxonomic classifications down to the species level, and geographic metadata (including coordinate information) associated with the sample collection site. The MAP was thus employed to identify the location data of 16S rRNA read occurrences of each species in the “confirmed bacterium–archaeon pairs” list. After identifying the 16S rRNA sample reads catalogued for a particular species, the read locations along with project and sample IDs (linking to project descriptions on the MAP database, which further characterize the geographic metadata) were exported to a spreadsheet. In the next step, the project IDs associated with the reads of each species within each respective group were cross-referenced to identify samples isolated from the same project ID (i.e., the same geographic location or microbiome). The projects found to contain 16S rRNA reads for each of the species within the final groups were identified via their respective ID in the MAP tool. Finally, environmental metadata including environmental descriptors and longitude and latitude coordinates for each particular read were identified. Through this process, we investigated the geographic habitat co-occurrence (referred to simply as “co-occurrence” throughout the remainder of the paper) of the pairs of “confirmed bacterium–archaeon pairs”, as well as descriptions of the unique environments that organisms in these groups inhabit.

## 3 Results

In the following section, Section 3.1 provides the results of assessing the effect of the random selection of a genome proxy on classification accuracy. Section 3.2 details the findings from the second experiment, focusing on the optimal values for *k*-mer size and genome proxy length, as well as the supervised classification accuracy using these optimal parameters. Finally, Section 3.3 presents the candidate bacterium–archaeon pairs identified through non-parametric methods, the results of filtering layers, the confirmed set of bacterium–archaeon pairs, the analysis of 3-mer frequency profile in these pairs, and the results of co-occurrence of “confirmed bacterium–archaeon pairs”.

### 3.1 Genome proxy

As described in Section 2.3.2, we conducted an experiment to assess the impact of a randomly selected genome proxy on taxonomic and environment-type classification under two different scenarios: the “restricted scenario” and the “restriction-free scenario”. For each scenario, we used 10-fold cross-validation classification with SVM classifier and repeated the classification process 10 times for each genome proxy length. To evaluate the results, we calculated the average accuracy and variance over the 10 runs for each genome proxy length. The results for the “restricted scenario,” are summarized in Table 3 (*Temperature Dataset*) and Table 4 (*pH Dataset*). For each tested genome proxy length, we reported the maximum average accuracy across the *k*-mer values and the value of *k* for which it was obtained. The results for the restriction-free scenario are similar and can be found in the Supplementary Materials, Section C.

**Table 3:** Maximum average accuracy across six genome proxy lengths in ten repeated SVM classification trials on the *Temperature Dataset* under the restricted scenario, for *k*-mer sizes 1 to 9. The table lists the highest average accuracy for each genome proxy length, alongside the *k*-mer size that achieved this accuracy and the variance in percentage. The *Temperature Dataset* has 598 samples, consisting of 369 bacteria and 229 archaea. There are 148 psychrophiles, 190 mesophiles, 183 thermophiles, and 77 hyperthermophiles in this dataset.

| Genome proxy length | Class labelling type | Max avg accuracy (%) | Variance (%) | $k$-value |
| --- | --- | --- | --- | --- |
| 10 kbp | Taxonomy | 98.35 | 0.0010 | 5 |
|  | Temperature | 67.39 | 0.0267 | 6 |
| 50 kbp | Taxonomy | 99.03 | 0.0001 | 6 |
|  | Temperature | 72.36 | 0.0224 | 7 |
| 100 kbp | Taxonomy | 99.13 | 0.0000 | 6 |
|  | Temperature | 73.18 | 0.0209 | 7 |
| 250 kbp | Taxonomy | 99.15 | 0.0000 | 6 |
|  | Temperature | 75.31 | 0.0035 | 9 |
| 500 kbp | Taxonomy | 99.15 | 0.0000 | 6 |
|  | Temperature | 76.97 | 0.0035 | 9 |
| 1,000 kbp | Taxonomy | 99.15 | 0.0000 | 6 |
|  | Temperature | 76.91 | 0.0013 | 9 |

**Table 4:** Maximum average accuracy across six genome proxy lengths in ten repeated SVM classification trials on the *pH Dataset* under the restricted scenario, for *k*-mer sizes 1 to 9. The table lists the highest average accuracy for each genome proxy length alongside the *k*-mer size that achieved this accuracy and the variance in percentage. The *pH Dataset* has 186 samples, consisting of 117 bacteria and 69 archaea. There are 100 acidophiles and 86 alkaliphiles in this dataset.

| Genome proxy length | Class labelling type | Max avg accuracy (%) | Variance (%) | $k$-value |
| --- | --- | --- | --- | --- |
| 10 kbp | Taxonomy | 97.18 | 0.0041 | 5 |
|  | pH | 83.17 | 0.0469 | 6 |
| 50 kbp | Taxonomy | 98.25 | 0.0022 | 7 |
|  | pH | 84.89 | 0.0040 | 7 |
| 100 kbp | Taxonomy | 98.63 | 0.0006 | 7 |
|  | pH | 85.61 | 0.0092 | 8 |
| 250 kbp | Taxonomy | 98.62 | 0.0012 | 8 |
|  | pH | 85.41 | 0.0053 | 9 |
| 500 kbp | Taxonomy | 98.94 | 0.0000 | 9 |
|  | pH | 86.20 | 0.0030 | 9 |
| 1,000 kbp | Taxonomy | 98.94 | 0.0000 | 9 |
|  | pH | 85.74 | 0.0035 | 9 |

In spite of the fact that each experiment was repeated 10 times, each time using a different randomly selected genome proxy, the maximum average accuracies are consistently high for taxonomy classifications and medium-high for environment-type classifications, with low variance across 10 different runs. These results support the hypothesis that the genomic signature, herein defined as the *k*-mer frequency vector of a short genomic fragment, is pervasive across the genome. Overall, these results indicate that selecting and pseudo-concatenating random regions of the genome into a contiguous genome proxy does not affect the taxonomic and environment-type classification accuracy, and is thus a valid selection method for these purposes.

The notable difference in environment-type classification accuracy between the two datasets can be partially attributed to the complexity of the classification task. Indeed, the *Temperature Dataset* has four unique labels while the *pH Dataset* has only two, making the latter an inherently simpler classification task.

### 3.2 Optimal ***k***-mer size and genome proxy length

The aim of this experiment is to identify the optimal *k*-mer size and the optimal genome proxy length for the purpose of taxonomy and environment-type classifications. To achieve this, we began by first determining the optimal *k*-mer size and then proceeded to determine the optimal genome proxy length. Our approach, especially when analyzing the various *k*-mer sizes, was to find a balance between computational time complexity/memory usage and classification accuracy.

Figure 3 presents the classification accuracy results of SVM classifiers applied to both the *Temperature Dataset* and the *pH Dataset* under the “restricted scenario,” with taxonomy and environment-type labelling, respectively. This figure illustrates how the classification accuracy changes as the value of *k* increases, for the six different genome proxy lengths analyzed. The classification accuracies for the other five classifiers, and for all six classifiers under the restriction-free scenario, for both the *Temperature Dataset* and the *pH Dataset* are similar, and can be found in the Supplementary Materials, Section D.

**Figure 3:**
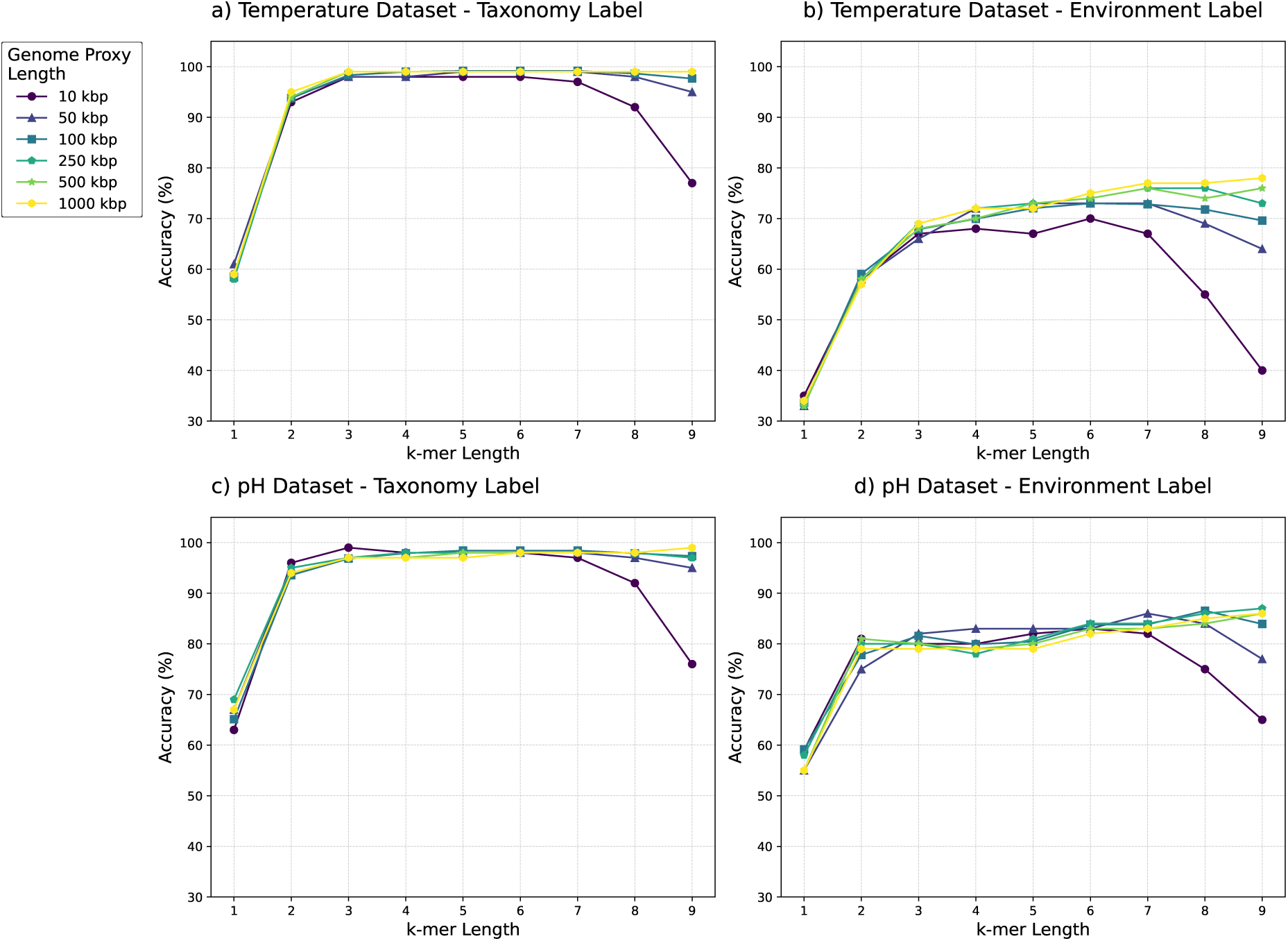
Classification accuracy of SVM classifier under restricted scenarios. *a: Temperature Dataset* with taxonomy labels. *b: Temperature Dataset* with environment-type labels. *c: pH Dataset* with taxonomy labels. *d: pH Dataset* with environment-type labels. Each subfigure shows accuracy across nine *k*-mer sizes and six genome proxy lengths.

As seen in Figure 3, increasing the length of *k*-mer from 1 to 6 leads to a significant improvement in classification accuracy. For values of *k* higher than 6, the changes in accuracy depend on the genome proxy length. For longer genome proxies (100 kbp, 250 kbp, 500 kbp, and 1,000 kbp), the taxonomic classification accuracy remains stable for increasing values of *k* from *k* = 6 to *k* = 9, and the environment-type classification accuracy increases with the increase in *k*-mer size. However, for shorter genome proxies (10 kbp and 50 kbp), both the taxonomic and environment-type accuracies decrease with the increase of *k*-mer sizes from 6 to 9.

The classification accuracy decline with the *k*-mer size increase, when *k* is higher than a certain threshold, is due to the fact that the increase in the length of the *k*-mer frequency feature vector is exponential in *k*. For small values of *k*, this increase results in more information available to the classifier. However, the number of *k*-mers that actually occur in the sequence is bound by the length of the sequence. Thus, after *k* passes a certain threshold, the feature vector becomes so sparse that it increasingly fails to capture the genomic patterns necessary for an accurate classification. This threshold is reached earlier for shorter sequences (10 kbp or 50 kbp) than for longer sequences.

Since, for all genome proxy lengths considered, the classification accuracies increase until *k* = 6, we concluded that the value of *k* should be 6 at the minimum, and performed a detailed analysis for values *k* = 6, 7, 8, 9.

The detailed analysis for *k*-mer sizes of 6 to 9 for the *Temperature Dataset* shows that the highest taxonomic classification accuracy for the six fragment lengths considered in the “restricted scenario”, ranges from 98.15% to 99.50%, and the highest environment-type classification accuracy ranges from 70.29% to 78.14%. Also, the results for the *pH Dataset*, indicate that the highest taxonomic classification accuracy for different fragment lengths ranges from 97.89% to 98.95%, and for environment-type classification ranges from 83.30% to 87.10%. The classifier’s performance in the “restriction-free” scenario is similar.

The detailed results of these experiments for both restricted and restriction-free scenarios can be found in Supplementary Materials, Section E. Overall, one observes that increasing the value of *k* from 6 to 9 does not result in significant increases in classification accuracy. This, combined with the fact that increasing *k* leads to an exponential increase in memory usage (the feature vector size increase from 2^12^ to 2^18^) and time complexity, leads to the conclusion that *k* = 6 is the optimal choice for the *k*-mer size in this context.

In the next step, we maintained a fixed *k*-mer size of *k* = 6 and assessed the effectiveness of six classifiers for the six genome proxy lengths considered in this study. This allowed us to identify the optimal genome proxy length for both the *Temperature Datasets* and *pH Dataset*. Table 5 displays the highest classification accuracy achieved for each genome proxy length, for both the “restriction-free scenario” and the “restricted scenarios”. As observed in Table 5, a fragment length of 100 kbp achieves the highest accuracy in three of the classification tasks: the restriction-free taxonomic classification for both datasets and the restricted taxonomic classification for the *Temperature Dataset*. In the remaining cases, the difference between the best performance and the 100 kbp performance was less than 0.5% in the restriction-free scenario and less than 2% in the restricted scenario. Thus, a genome proxy length of 100 kbp (at *k* = 6) is the optimal overall selection.

**Table 5:**
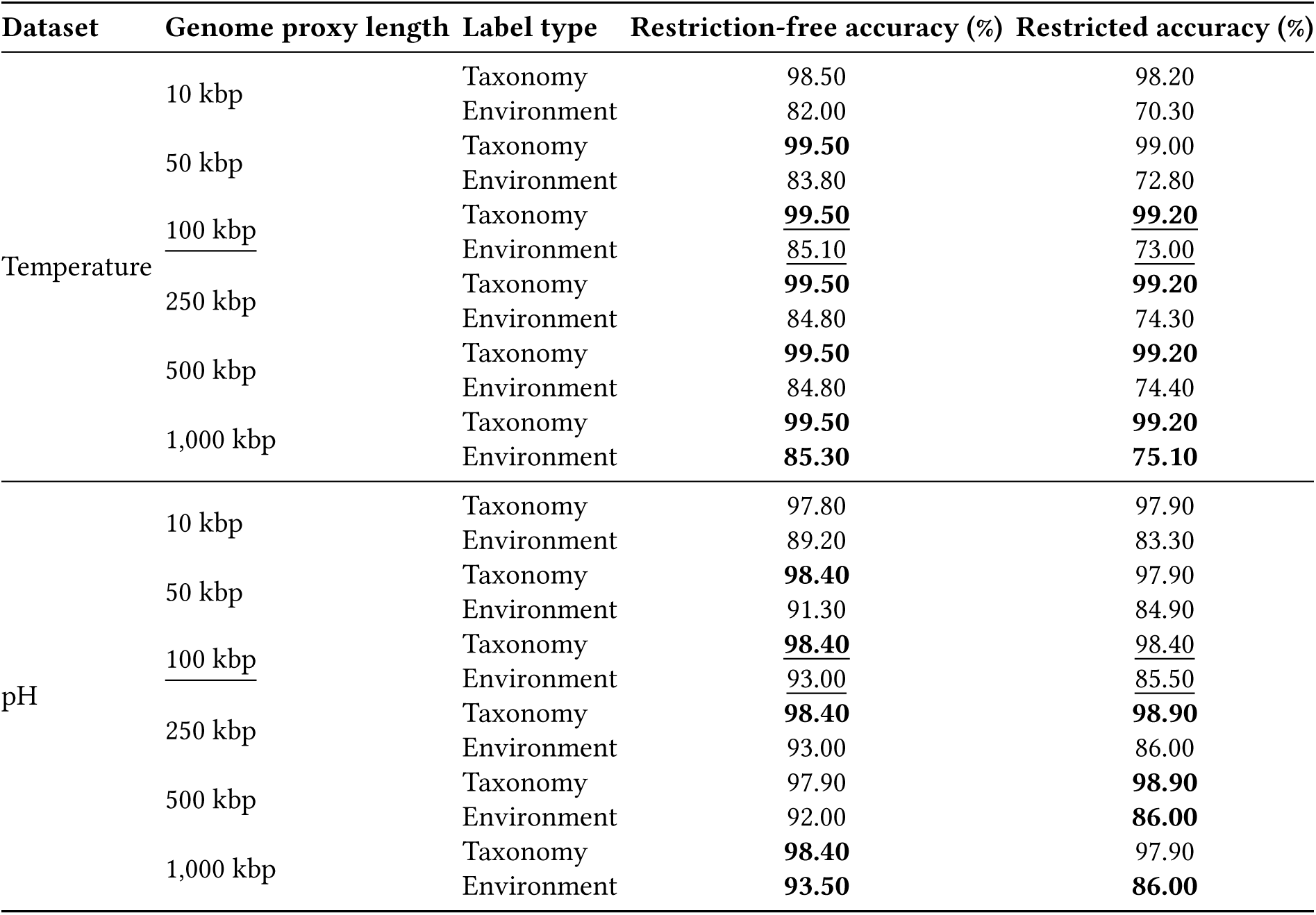
Comparison of the best classification accuracy across all classifiers, using *k* = 6, the optimal chosen value for *k*, for all six genome proxy lengths. All occurrences of maximum accuracy are shown in bold, and the performance for a fragment length of 100 kbp is shown as underlined.

In our last experiment, our objective was to determine whether using a short genome proxy might lead to any loss of information compared to using the whole genome. To evaluate this, we performed taxonomic and environment-type classification using entire genomes, while maintaining the same setup as our previous supervised experiment, under restricted scenarios with *k* equal to 6. Our findings show that for taxonomic classification of whole genomes with *k* = 6, the accuracy was 99.15% (compared to 99.20% using random 100 kbp genome proxies) for the *Temperature Dataset*, and 98.42% (compared to 98.40%) for the *pH Dataset*. For environment-type classification, the best accuracy for whole genomes was 75.51% (compared to 73.00%) for the *Temperature Dataset*, and 84.35% (compared to 85.50%) for the *pH Dataset*. These results indicate that classification accuracy using a genome proxy of length 100 kbp is comparable to using the entire genome, which in our datasets has an average length of 3,500 kbp (the genome proxy is 35 times shorter on average).

### 3.3 A multi-layered pipeline to find bacterium-archaeon pairs with similar genomic signatures

The identification of bacterium–archaeon pairs is a multi-layered filtering process that progressively narrows down the candidate pairs generated through unsupervised clustering, to reach the confirmed bacterium–archaeon pairs. We started with non-parametric clustering, computed the FCGR of genome proxies of all DNA sequences in the datasets, and filtered the candidate bacterium–archaeon pairs generated by the non-parametric clustering based on their FCGR distances. Additionally, we conducted biological analysis, including isolating the environment metadata analysis. This approach allowed us to identify 15 bacterium–archaeon pairs that passed the biological analysis and the FCGR analysis. Figure 4 illustrates the details of this multi-layered filtering approach. We further investigated the 3-mer usage bias in these 15 bacterium–archaeon pairs (which passed all filtering layers) and found that they demonstrate a similar genomic signature linked to their extreme environment despite their maximal taxonomic differences. As the last analysis, we also studied the co-occurrence of “confirmed bacterium–archaeon pairs”.

**Figure 4:**
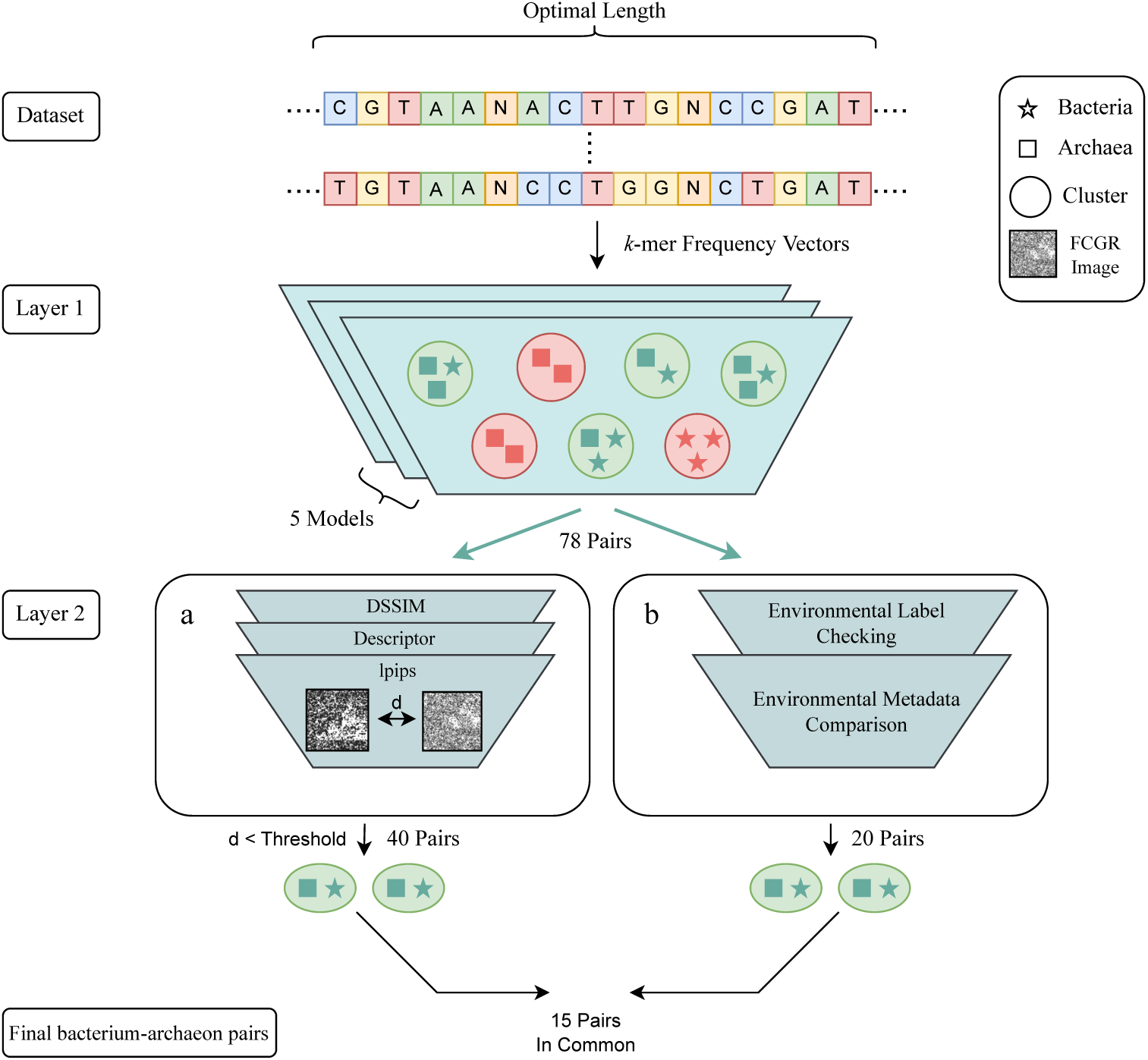
Multi-layered pipeline for identifying bacterium/archaeon pairs with similar genomic signatures. *Layer 1:* Five selected non-parametric clustering methods identify clusters of organisms with similar genomic signatures. The clusters containing both bacteria and archaea (green) generate a list of 78 candidate bacterium–archaeon pairs, grouped by these algorithms based on their similar genomic signatures. *Layer 2-a:*: The candidate pairs from Layer 1 undergo pairwise distance calculations between their FCGRs using four different distance metrics. Only 40 pairs, with the majority of distances below empirically determined thresholds, are retained. *Layer 2-b:* In parallel to FCGR comparison, a biological analysis is conducted on the output pairs from Layer 1. This includes checking environment labels and examining metadata about their living environments to select pairs isolated from similar types of extreme environments, resulting in 20 pairs. The final output is a list of 15 bacterium/archaeon pairs (comprising 16 unique genera and 20 unique species) that passed all filtering layers. These pairs can confidently be proposed as maximally divergent microbes that share similar genomic signatures associated with their living environments.

#### 3.3.1 Layer 1: non-parametric clustering

We initiated the process using non-parametric clustering algorithms in combination with dimensionality reduction methods. As described in Section 2.4.1, we evaluated the contamination and completeness scores of the clusters and identified the top five performing clustering methods, selecting those that performed best at generating clusters that correspond to true genera.

From the clusters obtained using the chosen algorithms, a set of candidate pairs, consisting of bacterium–archaeon pairs whose genomic signatures were consistently clustered together by the majority of the algorithms, was identified for each dataset. To ensure robustness, we repeated the above analysis (clustering, and selecting bacterium–archaeon pairs) 10 times. We then selected the bacterium–archaeon pairs that appeared in at least 5 of the 10 runs. This initial step generated 78 bacterium–archaeon candidate pairs (38 unique genera, 85 unique species) which were named “initial bacterium–archaeon candidate pairs”.

#### 3.3.2 Layer 2-a: FCGR comparison of candidate pairs

In the second step, we filtered the “initial bacterium–archaeon candidate pairs” based on their FCGR distances. As described in Section 2.4.2, we calculated the FCGR images for each pair of sequences, using a genome proxy length of 100 kbp and *k* value of 6, and measured the distances between these FCGRs using three distance metrics. We selected bacterium–archaeon candidate pairs with distances below empirically determined thresholds for the majority of distance metrics.

After this filtering layer, we identified 40 bacterium–archaeon candidate pairs (32 unique genera, 48 unique species), with similar FCGR images, determined by the three distance metrics. These candidate pairs are considered “FCGR-accepted bacterium–archaeon candidate pairs” and pass this filtering layer. Figure 5 shows the FCGR images of two pairs, one extremophile (*Thermotoga petrophila* and *Geoglobus acetivorans*) and one polyextremophile pair (*Thermoanaerobacterium thermosaccharolyticum* and *Caldisphaera lagunensis*). For better visualization, the value *k* = 8 was used, and the images confirmed that the FCGRs show visual pattern similarities, in addition to the distance between FCGRs being below the empirically determined threshold.

**Figure 5:**
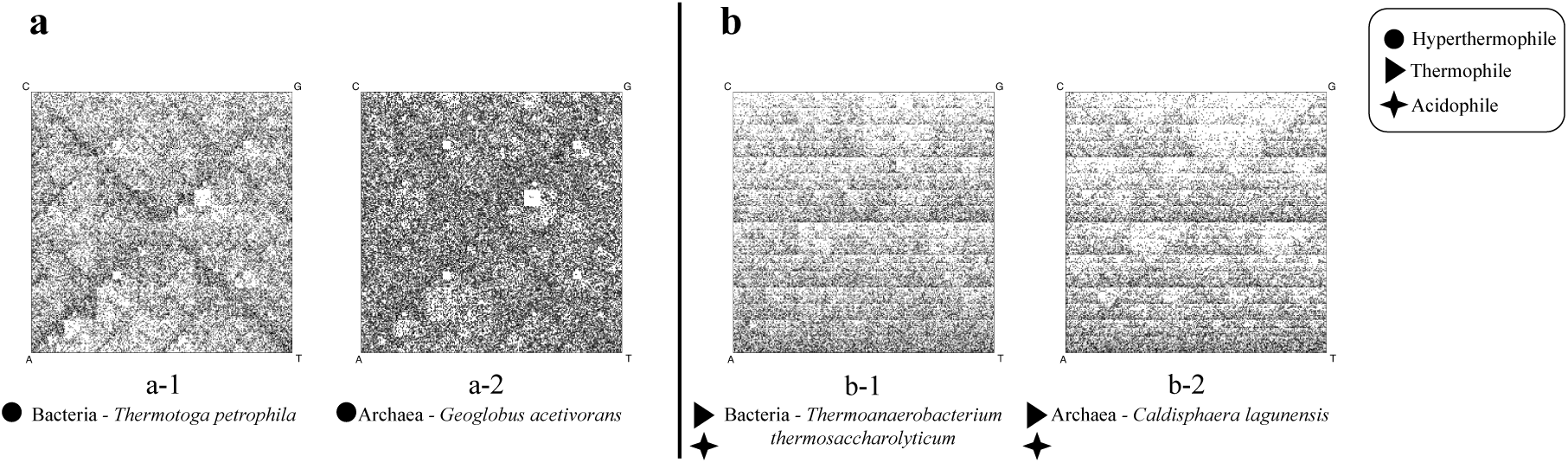
FCGR images of two candidate pairs (four unique species) were selected from the “FCGR-accepted bacterium–archaeon candidate pairs” list, with a resolution of *k* = 8. The first candidate pair includes *a-1:* a hyperthermophilic bacterium and *a-2:* a hyperthermophilic archaeon, while the second pair consists of *b-1:* an acidophilic thermophilic bacterium and *b-2:* an acidophilic thermophilic archaeon. The first pair was drawn from the *Temperature Dataset*, and the second pair appears in both *Temperature Dataset* and *pH Dataset*. In both candidate pairs, the FCGRs display strikingly similar patterns between the two species, despite belonging to different taxonomic domains (Bacteria and Archaea).

#### 3.3.3 Layer 2-b: analysis of candidate bacterium-archaeon pairs considering phenotypic and isolating environment metadata

As outlined in Section 2.4.3, an environmental metadata analysis complemented the FCGR comparison, serving as an additional refinement step. This analysis involved comparing the environmental labels of the species comprising the microbial pairs identified by non-parametric clustering. As previously mentioned, this included identifying the original study that reported and characterized a given species, which enabled us to both annotate each species according to its isolating environment label and conduct a detailed comparison assessing the isolating environments and growth parameters.

Based on our analysis, out of 78 candidate pairs obtained from the non-parametric clustering layer, a total of 20 candidate pairs (17 unique genera and 25 unique species) passed this filtering stage based on their environment metadata and labels. The details of this environmental data of the selected pairs can be found in the Supplementary Materials, Section F.

The bacterium–archaeon candidate pairs that passed both the FCGR filtering layer and the biological analysis layer resulted in 15 pairs (16 unique genera and 20 unique species), which we propose as the “confirmed bacterium–archaeon pairs” set. These pairs are organized into five groups based on their environmental metadata, with each group sharing nearly matching or matching environment labels. Notably, Groups 1, 2, and 3 include sequences of organisms isolated from extreme environments, while the majority of organisms in Groups 4 and 5 are associated with normal temperature (mesophiles) and normal pH (absent from the *pH Dataset*) conditions. We further examined the 3-mer usage bias of species in these confirmed 15 pairs, as well as their co-occurrences. For Groups 4 and 5, we also investigated any potential extreme conditions in their environments other than extreme temperature or pH.

The details of these five groups are shown in Figure 6, and their FCGR images can be found in Supplementary Materials, Section G.

**Figure 6:**
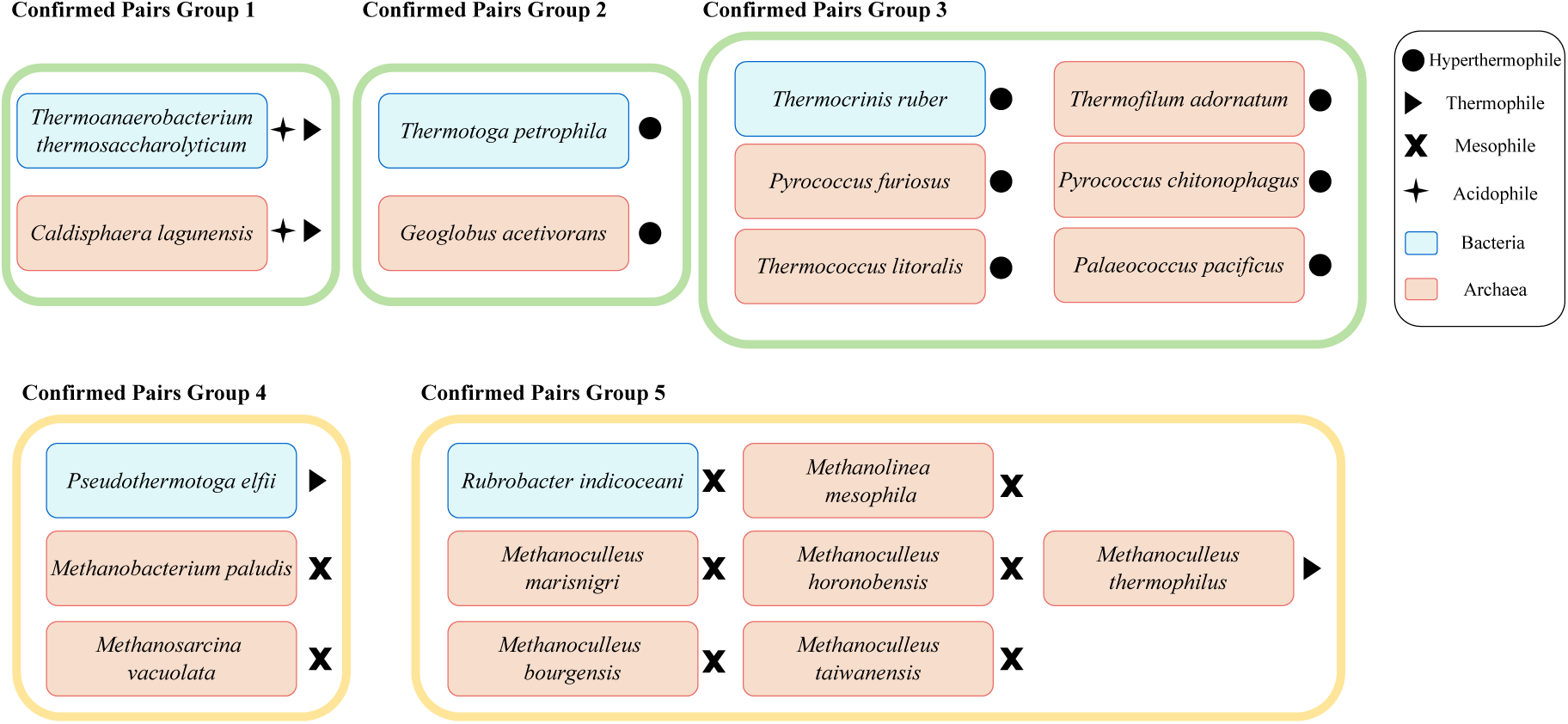
Confirmed bacterium–archaeon pairs after two filtering layers. These pairs passed both FCGR comparison and environment similarity analysis, resulting in 3 groups of pairs with extreme environment conditions (*green*) and 2 groups with normal environment conditions (*yellow*). The confirmed pairs set comprises 20 species, including five bacteria and 15 archaea from 16 unique genera. Among these, two species are poly-extremophiles (acidophilic thermophiles), 10 are extremophiles (eight hyperthermophiles and two thermophiles), and the remaining eight are mesophiles.

#### 3.3.4 Analysis of 3-mer frequency profiles of confirmed bacterium-archaeon pairs

To investigate potential biases in 3-mer usage associated with environmental adaptation, we conducted a detailed analysis of the 3-mer frequency profiles for the genome proxies of the organisms in the “confirmed bacterium–archaeon pairs” groups. We focused on *k* = 3 due to its biological relevance, since the set of codons is a subset of the set of 3-mers. Following the four-step analysis outlined in Section 2.4.4, this section examines how 3-mer frequencies reflect environmental adaptations across taxonomically divergent microbes. The results of this analysis are summarized in Table 6 for each of the five “confirmed bacterium–archaeon pair” groups.

**Table 6:** Combined summary of 3-mer profile analysis for Groups 1 to 5. For each pair, we calculated the number of shared environment-relevant 3-mers exhibiting similar over– or under-representation patterns between the two species of the pair, and reported the ratio out of 15. The Spearman rank correlation coefficient (*r*ℎ*o*) was computed to quantify the correlation between the 3-mer representation patterns of each pair; all correlations were statistically significant (*p <* 10^−5^). A combined score was calculated as the average of the shared environmentrelevant 3-mer ratio and *r*ℎ*o* to assess the overall similarity of each pair. For further validation, the last column reports the number of shared environment-relevant 3-mers that have over-or under-representation patterns consistent with findings in the biological literature.

| Group | Bacterium | Archaeon | Shared environment relevant 3-mers (ratio) | $\rho$ | Score | Overall 3-mer similarity | Biology literature observed shared 3-mers |
| --- | --- | --- | --- | --- | --- | --- | --- |
| Group 1 | *Thermoanaerobacterium thermosaccharolyticum* | *Caldisphaera lagunensis* | 0.83 | 0.96 | 0.89 | Compelling | 9 |
| Group 2 | *Thermotoga petrophila* | *Geoglobus acetivorans* | 1.00 | 0.81 | 0.90 | Compelling | 4 |
| Group 3 | *Thermocrinis ruber* | *Thermofilum adornatum* | 1.00 | 0.77 | 0.89 | Compelling | 5 |
|  |  | *Pyrococcus chitonophagus* | 1.00 | 0.81 | 0.90 | Compelling | 4 |
|  |  | *Palaeococcus pacificus* | 0.90 | 0.76 | 0.83 | Very strong | 4 |
|  |  | *Pyrococcus furiosus* | 0.89 | 0.81 | 0.85 | Compelling | 5 |
|  |  | *Thermococcus litoralis* | 0.91 | 0.80 | 0.85 | Compelling | 4 |
| Group 4 | *Pseudothermotoga elfii* | *Methanobacterium paludis* | 1.00 | 0.94 | 0.97 | Compelling | 1 |
|  |  | *Methanosarcina vacuolata* | 0.75 | 0.95 | 0.85 | Compelling | 1 |
| Group 5 | *Rubrobacter indicooceani* | *Methanolinea mesophila* | 0.60 | 0.83 | 0.71 | Moderate | 1 |
|  |  | *Methanoculleus marisnigri* | 0.75 | 0.94 | 0.84 | Very strong | 2 |
|  |  | *Methanoculleus bourgensis* | 0.73 | 0.91 | 0.82 | Very strong | 2 |
|  |  | *Methanoculleus horonobensis* | 0.69 | 0.95 | 0.82 | Very strong | 3 |
|  |  | *Methanoculleus taiwanensis* | 0.64 | 0.92 | 0.78 | Strong | 3 |
|  |  | *Methanoculleus thermophilus* | 0.78 | 0.89 | 0.83 | Very strong | 0 |

For each confirmed bacterium-archaeon pair, the set of “shared environment-relevant 3-mers” is defined as the intersection of the set of environment-relevant 3-mers of the bacterium genome proxy with that of the archaeon genome proxy. Among these shared 3-mers, we calculate the proportion of 3-mers that show the same pattern of over-or under-representation in both species and reported it in Table 6. Additionally, we calculated the Spearman rank correlation coefficient (*r*ℎ*o*) between the 3-mer representation patterns of the two organisms in each pair. Notably, all correlations were statistically significant with *p <* 10^−5^ for all pairs. Since the shared 3-mer ratio and *r*ℎ*o* collectively represent the results of steps 1 to 3 of the 3-mer frequency profile analysis pipeline (see Section 2.4.4), we calculated a combined score as the average of these two values to provide an overall measure of 3-mer frequency profile similarity between the species of each pair. Based on the combined score, we also assigned a descriptive term to each pair for a clear comparison. Specifically, we labeled pairs as “Compelling” for scores greater than or equal to 0.85, “Very strong” for scores between 0.85 and 0.80, “Strong” for scores between 0.80 and 0.75 and “Moderate” for scores between 0.75 and 0.70. These thresholds were determined empirically based on the distribution of our results.

Finally, as outlined in the last step of the 3-mer frequency profile analysis (see Section 2.4.4), we examined the biological literature on codon usage to determine whether the observed over-or under-representation of each shared environment-relevant 3-mer had been previously reported in biological literature. The final column of Table 6 reports the number of shared 3-mers for which our findings in over-or under-representation align with evidence from prior studies, providing further validation of the observed similarities. Importantly, we did not include this literature-based validation in the combined score calculation, as low values in this step may only reflect a lack of prior research in the literature rather than true biological absence.

The results revealed eight pairs with compelling 3-mer similarity, five pairs with very strong similarity, one pair with strong similarity and one pair with moderate similarity. No pairs exhibited very low similarity (the minimum similarity score is 0.71), indicating a moderate to high level of 3-mer frequency profile similarity across all confirmed pairs. Notably, the first three groups, which include poly-extremophile or extremophiles, showed a higher average number of shared environment-relevant 3-mers observed in the biological literature (average: five) compared to Groups 4 and 5, which predominantly consist of mesophiles (average: two).

Interestingly, despite being composed mainly of mesophiles, Groups 4 and 5 included two pairs with compelling similarity and four pairs with very strong similarity. This unexpected finding suggests that factors beyond temperature or pH, such as other environmental pressures, may contribute to genomic convergence in these pairs, which is further discussed in Section 4. Detailed results of the 3-mer frequency profile analysis are presented in Supplementary Materials, Section H.

#### 3.3.5 Co-occurrence of organisms from the confirmed bacterium-archaeon pairs

In the final analysis, using the MAP tool [39], we analyzed the habitats of all “confirmed bacterium–archaeon pairs” within their respective groups, to identify any shared environments, as outlined in Section 2.4.5.

This analysis revealed distinct patterns of co-occurrence across different groups. Both species in **Group 1** were found together in Washburn Hot Springs, a geothermal hot spring in Yellowstone National Park, Wyoming, USA [43]. Notably, this co-occurrence habitat differs from the environments where the species were originally isolated [24, 37, 40]. Despite the large geographic distances between the original isolation and co-occurrence sites, these habitats have similar environmental pressures and geochemical properties.

Similar observations were made for the species in the pair of **Group 2**, which were found to co-occur in two distinct habitats: Brothers Volcano, a submarine volcano in the Pacific Ocean near New Zealand [58], and Juan de Fuca Ridge, a mid-ocean ridge flank near Vancouver Island [27]. Note that these species were initially isolated from a deep Japanese oil reservoir [67] and a deep-sea hydrothermal vent [64], respectively.

In **Group 3**, a subset of species co-occurred in environments overlapping with those of Group 1 and Group 2, including Brothers Volcano and Washburn Hot Springs. Additional co-occurrence sites were found across Yellowstone National Park. Similar to Group 1 and Group 2, the environmental conditions of the co-occurrence habitats resemble the conditions of isolating environments of the respective species. It is worth mentioning that even though species from Groups 1, 2, and 3 were found to co-occur in the same habitat, our clustering methods provide the sensitivity to detect specific 3-mer biases within their genomic signatures. This enables classification based on their evolved niche adaptations rather than their current habitat, which explains why these groups were clustered separately despite sometimes sharing the same environment. Detailed geographic maps and cooccurrence data for these three groups can be found in the Supplementary Materials Section I.

Although the majority of species in **Group 4** are mesophiles, they co-occurred in multiple independent environments characterized by other common extreme environment conditions such as anaerobic, and methanogenic conditions. These habitats include the Shengli Oil Field in China, hypothesized to involve anaerobic, mesophilic microbiomes in the “methanogenic degradation of hydrocarbons” [53], and a Japanese bioreactor [32]. No cooccurrence was identified for **Group 5** species. Detailed geographic maps and co-occurrence for Group 4 and Group 5 can be found in Supplementary Materials Section I.

Importantly, this co-occurrence analysis supports the bacterium–archaeon pairs clusters identified by our multilayered approach. Indeed, it demonstrates that many of the species pairs that were computationally grouped together by our method, despite being originally isolated from different environments, were later found to cooccur naturally in shared environments distinct from their isolation sites.

## 4 Discussion

Our computational analysis revealed that both taxonomic and environmental components can be pervasive throughout extremophile prokaryotic genomes, suggesting that environmental adaptations influence the entire genome rather than specific genic or regulatory regions exclusively. Indeed, our novel computational pipeline resulted in high classification and clustering accuracies, despite using as “genome proxy” a relatively short DNA fragment constructed by the pseudo-concatenation of 10 randomly selected 10,000 bp fragments (total length 100,000 bp, that is ≈ 35 times shorter than a complete genome). This indicates that taxonomic and environmental components are detectable even with limited genomic samples, which has important implications for studying environmental adaptations when complete genome sequences are not available.

Our multi-layered approach identified 15 pairs of maximally distant organisms that have similar genomic signatures, grouped into five distinct categories. The statistically significant 3-mer over-representation and underrepresentation analysis further confirmed the genomic composition similarity of these pairs. Notably, the identified environmentally-relevant 3-mer representation patterns align with known extremophile adaptation mechanisms as detailed below.

In **Group 1**, the over-representation of the 3-mer “CAA” (corresponding to a glutamine codon) in the genome of thermophilic acidophiles aligns with previous findings of codon usage bias in acidophilic prokaryotes which prefer the “CAA” codon when calling for glutamine [31]. Moreover, the under-representation of the 3-mer “ACG” (corresponding to a threonine codon) in this group is consistent with amino acid abundance patterns found in thermophilic prokaryotic proteins which demonstrate a relative lack of threonine [19].

In **Group 2** and **Group 3**, consisting of hyperthermophiles, observations of the elevated representation of 3-mers corresponding to arginine codons, and decreased representation of 3-mers corresponding to asparagine and glutamine codons align with previous observations related to amino acid abundances in hyperthermophilic and thermophilic prokaryotic proteins [76, 12, 68]. Specifically, (hyper)thermophilic proteins demonstrate an increased abundance of arginine, and decreased abundance of asparagine and glutamine amino acids, which is reflected by the relative representations of 3-mers respectively. Note that several species in Group 3, specifically, *Thermocrinis ruber* (bacteria) and three archaeal species (*Pyrococcus furiosus*, *Thermococcus litoralis*, and *Pyrococcus chitonophagus*), were previously identified as having similar genomic signatures by using slightly different methods [2], which further validates our multi-layered approach.

The 3-mer frequency profile analysis of **Group 4** also showed some agreement with known codon usage patterns. In this group, all species, including the thermophilic bacterium, exhibited an under-representation of 3-mers corresponding to a serine codon. This pattern aligns for mesophilic species, which demonstrate a codon usage bias against the 3-mer “AGC” when calling for serine [6] and with the observed lower serine amino acid in thermophilic proteins relative to mesophilic proteins [33]. The grouping of mesophilic species from maximally divergent taxa in Group 4, along with their similarity in genomic compositions and 3-mer representations, suggests the influence of extreme environmental pressures beyond temperature and pH. Indeed, we observed that Group 4 species co-occur in anaerobic, methanogenic environments and share the phenotypic trait of oxygen intolerance (Supplementary Materials Section F). This indicates that additional extreme factors, such as high concentrations of endogenously-produced methane, or exogenous hydrocarbons encountered in oil fields or wells, could potentially influence extremophilic genomic signature composition.

In **Group 5**, in contrast with Groups 1, 2, 3 and 4, our findings revealed unique genomic signature patterns that differ from previous biological findings of extremophile codon usage bias. In this group, our findings showed an under-representation of the 3-mer “CTA,” which codes for leucine. This was expected in mesophilic species of this group, as mesophilic prokaryotes commonly exhibit a bias against using this codon [6]. However, surprisingly, we found the same under-representation in the thermophilic species of this group, in contrast with previous studies which showed “CTA” to be typically abundant in other thermophiles [76]. Our finding contradicts previous assumptions of codon usage bias in thermophilic prokaryotes, suggesting that the impact of environmental adaptation on prokaryotic genomes may be more nuanced than previously thought and needs further investigation. Although no co-occurrence environments were found for Group 5, their initial isolation from predominantly methanogenic habitats, as described in their discovering papers, suggests a potential role of methanogenic processes in shaping the genomic signatures of these species. Further investigation is needed to clarify these relationships.

Our computational approach also has some limitations. For example, this *k*-mer based method cannot capture long-range genomic interactions, although this could potentially be addressed through the use of transformer models [71]. Additionally, the exponential growth in the size of feature vectors with increasing *k*-mer size limited our analysis to *k* ≤ 9, potentially obscuring larger sequence patterns. Also, while the parameters that were empirically determined to be optimal proved effective for the classification/clustering of this extremophile dataset, they may not generalize across all genomic analyses, as they likely depend on dataset characteristics and the complexity of the classification task. Lastly, the choice of *n* = 10 DNA sub-fragments to pseudo-concatenate into a “representative DNA fragment” to act as genome proxy for these computational analyses is not absolute, and represented a necessary trade-off between computational efficiency and genomic coverage.

Overall, our findings demonstrate that extreme environmental adaptation significantly impacts prokaryotic genomic signature compositions, with environmental pressures capable of overriding traditionally recognized taxonomic influences. The biological significance of our approach is highlighted by the discovery of 15 microbial species pairs that share genomic signatures despite maximal taxonomic divergence, suggesting that shared environmental pressures can drive convergent genomic adaptations across vastly different species. These results provide compelling evidence that environment-driven genomic components persist across diverse taxa, offering new perspectives on how environment-associated mutagenesis and selection shape microbial genomes. Our work broadens the field’s perspective beyond the traditional focus on phenotype, proteome, and gene-specific analyses to genome-wide considerations. By bridging computational methods with biological context, this work advances machine learning applications in genomics and our understanding of extremophile adaptation mechanisms. Future research will explore the biological mechanisms underlying these shared genomic signatures and their implications for evolutionary biology, biotechnology, and environmental genomics.

## 5 Data Availability

All sequence data used in this paper is publicly available for download at NCBI. The unique assembly accession IDs of all the sequences and their respective labels used in this study and the representative genome proxies used in this paper are available at https://github.com/Kari-Genomics-Lab/Extreme_Env_2.

## 6 Code availability

All the codes used in this study are available at: https://github.com/Kari-Genomics-Lab/Extreme_Env_ 2

## 7 Author Contributions Statement

L.K., K.A.H., M.S. designed the computational experiment, M.S. and G.S.R. conducted the computational experiments. J.B. and K.A.H. provided the dataset curation and biological interpretation of the findings. M.S., L.K., K.A.H., and J.B. contributed to the writing of the manuscript, and M.S. and J.B. analyzed the results, and all authors reviewed the manuscript.

