## Supplementary Materials for "Life at the extremes: Maximally divergent microbes with similar genomic signatures linked to extreme environments"

### Supplementary Materials for the paper "Life at the extremes: Maximally divergent microbes with similar genomic signatures linked to extreme environments"

Monireh Safari[1,*], Joseph Butler[2], Gurjit S. Randhawa[3], Kathleen A. Hill[2], and Lila Kari[1,*]

[1]School of Computer Science, University of Waterloo
[2]Department of Biology, University of Western Ontario
[3]School of Computer Science, University of Guelph

June 3, 2025

### A Dataset Details

In this study, we used two datasets of microbial genomes that live in extreme temperature (*Temperature Dataset*) or extreme pH (*pH Dataset*). In total, there are 91 species that are shared in both the *Temperature dataset* and *pH dataset*. Eight genomes are mesophiles and live in acidic or alkaline environments, as detailed in Table 1. From the remaining 83 genomes that are poly-extremophiles, there are 40 Archaea (Table 2), and 43 Bacteria (Table 3).

*Table 1:* Genomes that are listed in both the *Temperature* and *pH datasets*. The domain of all the sequences in this category is Archaea. These genomes are not poly-extremophiles, as their temperature label is mesophile.

| **Assembly ID** | **Species** | **Temperature Label** | **pH Label** |
|---|---|---|---|
| GCA_000336935.1 | *Halococcus salifodinae* | Mesophiles | Alkaliphiles |
| GCA_000337135.1 | *Natrialba chahannaoensis* | Mesophiles | Alkaliphiles |
| GCA_000337575.1 | *Natrialba hulunbeirensis* | Mesophiles | Alkaliphiles |
| GCA_001971705.1 | *Natronorubrum daqingense* | Mesophiles | Alkaliphiles |
| GCA_004745425.1 | *Methanolobus sp004745425* | Mesophiles | Alkaliphiles |
| GCA_014647115.1 | *Halarchaeum rubridurum* | Mesophiles | Acidophiles |
| GCA_900104065.1 | *Natronobacterium texcoconense* | Mesophiles | Alkaliphiles |
| GCA_900188065.1 | *Halorubrum vacuolatum* | Mesophiles | Alkaliphiles |

*Table 2:* The poly-extremophiles from Archaea domain, shared between both the *Temperature Dataset* and *pH Dataset.*

| **Assembly ID** | **Species** | **Temperature Label** | **pH Label** |
|---|---|---|---|
| GCA_000008665.1 | *Archaeoglobus fulgidus* | Hyperthermophiles | Alkaliphiles |
| GCA_000018305.1 | *Caldivirga maquilingensis* | Hyperthermophiles | Acidophiles |
| GCA_000018365.1 | *Thermococcus onnurineus* | Hyperthermophiles | Alkaliphiles |
| GCA_000022485.1 | *Saccharolobus islandicus* | Hyperthermophiles | Acidophiles |
| GCA_000148385.1 | *Vulcanisaeta distributa* | Hyperthermophiles | Acidophiles |
| GCA_000215995.1 | *Pyrococcus yayanosii* | Hyperthermophiles | Acidophiles |
| GCA_000223395.1 | *Pyrolobus fumarii* | Hyperthermophiles | Acidophiles |
| GCA_000253055.1 | *Thermoproteus tenax* | Hyperthermophiles | Acidophiles |
| GCA_001748385.1 | *Vulcanisaeta_B thermophila* | Hyperthermophiles | Acidophiles |
| GCA_002116695.1 | *Acidianus manzaensis* | Hyperthermophiles | Acidophiles |
| GCA_009729015.1 | *Acidianus ambivalens* | Hyperthermophiles | Acidophiles |
| GCA_014646555.1 | *Vulcanisaeta souniana* | Hyperthermophiles | Acidophiles |
| GCA_900079115.1 | *Saccharolobus solfataricus* | Hyperthermophiles | Acidophiles |
| GCA_000011185.1 | *Thermoplasma volcanium* | Thermophiles | Acidophiles |
| GCA_000012285.1 | *Sulfolobus acidocaldarius* | Thermophiles | Acidophiles |
| GCA_000025665.1 | *Aciduliprofundum boonei* | Thermophiles | Acidophiles |
| GCA_000144915.1 | *Acidilobus saccharovorans* | Thermophiles | Acidophiles |
| GCA_000193375.1 | *Thermoproteus uzoniensis* | Thermophiles | Acidophiles |
| GCA_000195915.1 | *Thermoplasma acidophilum* | Thermophiles | Acidophiles |
| GCA_000196895.1 | *Halalkalicoccus jeotgali* | Thermophiles | Alkaliphiles |
| GCA_000204925.1 | *Metallosphaera cuprina* | Thermophiles | Acidophiles |
| GCA_000243315.1 | *Metallosphaera yellowstonensis* | Thermophiles | Acidophiles |
| GCA_000317795.1 | *Caldisphaera lagunensis* | Thermophiles | Acidophiles |
| GCA_000336615.1 | *Haloarcula amylolytica* | Thermophiles | Alkaliphiles |
| GCA_000336755.1 | *Haloferax elongans* | Thermophiles | Acidophiles |
| GCA_000337735.1 | *Natronorubrum sulfidifaciens* | Thermophiles | Alkaliphiles |
| GCA_000455345.1 | *Halopiger_A goleimassiliensis* | Thermophiles | Alkaliphiles |
| GCA_000508305.1 | *Sulfolobus acidocaldarius_A* | Thermophiles | Acidophiles |
| GCA_000632495.1 | *Acidianus copahuensis* | Thermophiles | Acidophiles |
| GCA_001719125.1 | *Saccharolobus sp001719125* | Thermophiles | Acidophiles |
| GCA_002153915.1 | *Methanonatronarchaeum thermophilum* | Thermophiles | Alkaliphiles |
| GCA_003201675.2 | *Metallosphaera hakonensis* | Thermophiles | Acidophiles |
| GCA_003201765.2 | *Acidianus sulfidivorans* | Thermophiles | Acidophiles |
| GCA_003201835.2 | *Acidianus_B brierleyi* | Thermophiles | Acidophiles |
| GCA_003967175.1 | *Sulfodiicoccus acidiphilus* | Thermophiles | Acidophiles |
| GCA_009729035.1 | *Stygiolobus azoricus* | Thermophiles | Acidophiles |
| GCA_009729055.1 | *Sulfurisphaera ohwakuensis* | Thermophiles | Acidophiles |
| GCA_013340765.1 | *Conexivisphaera calida* | Thermophiles | Acidophiles |
| GCA_013343295.1 | *Metallosphaera tengchongensis* | Thermophiles | Acidophiles |
| GCA_900176435.1 | *Picrophilus oshimae* | Thermophiles | Acidophiles |

*Table 3:* The poly-extremophiles from Bacteria domain, shared between both the *Temperature Dataset* and *pH Dataset.*

| **Assembly ID** | **Species** | **Temperature Label** | **pH Label** |
|---|---|---|---|
| GCA_000014185.1 | *Rubrobacter_B xylanophilus* | Thermophiles | Alkaliphiles |
| GCA_000015025.1 | *Acidothermus cellulolyticus* | Thermophiles | Acidophiles |
| GCA_000024285.1 | *Alicyclobacillus acidocaldarius* | Thermophiles | Acidophiles |
| GCA_000092425.1 | *Truepera radiovictrix* | Thermophiles | Alkaliphiles |
| GCA_000145615.1 | *Thermoanaerobacterium thermosaccharolyticum* | Thermophiles | Acidophiles |
| GCA_000175575.2 | *Acidithiobacillus_A caldus* | Thermophiles | Acidophiles |
| GCA_000219875.1 | *Alicyclobacillus acidocaldarius_A* | Thermophiles | Acidophiles |
| GCA_000226295.1 | *Chloracidobacterium thermophilum* | Thermophiles | Alkaliphiles |
| GCA_000421625.1 | *Thermus islandicus* | Thermophiles | Acidophiles |
| GCA_000429525.1 | *Alicyclobacillus_A contaminans* | Thermophiles | Acidophiles |
| GCA_000430585.1 | *Alicyclobacillus_G herbarius* | Thermophiles | Acidophiles |
| GCA_000444055.1 | *Alicyclobacillus acidoterrestris* | Thermophiles | Acidophiles |
| GCA_000472905.1 | *Alicyclobacillus_A pomorum* | Thermophiles | Acidophiles |
| GCA_000833605.1 | *Anoxybacillus ayderensis* | Thermophiles | Alkaliphiles |
| GCA_000953475.1 | *Methylacidiphilum fumariolicum* | Thermophiles | Acidophiles |
| GCA_001280565.1 | *Sulfobacillus thermosulfidooxidans_A* | Thermophiles | Acidophiles |
| GCA_001447355.1 | *Alicyclobacillus tengchongensis* | Thermophiles | Acidophiles |
| GCA_001552255.1 | *Alicyclobacillus_G shizuokensis* | Thermophiles | Acidophiles |
| GCA_001552655.1 | *Alicyclobacillus_G kakegawensis* | Thermophiles | Acidophiles |
| GCA_001552675.1 | *Alicyclobacillus sendaiensis* | Thermophiles | Acidophiles |
| GCA_001570745.1 | *Alicyclobacillus mali* | Thermophiles | Acidophiles |
| GCA_003057965.1 | *Thermodesulfobium acidiphilum* | Thermophiles | Acidophiles |
| GCA_004366795.1 | *Alicyclobacillus sacchari* | Thermophiles | Acidophiles |
| GCA_007475525.1 | *Methylacidiphilum kamchatkense* | Thermophiles | Acidophiles |
| GCA_007991715.1 | *Alicyclobacillus acidoterrestris_A* | Thermophiles | Acidophiles |
| GCA_013760845.1 | *Anoxybacillus_B calidus* | Thermophiles | Alkaliphiles |
| GCA_014196195.1 | *Anoxybacillus_A rupiensis* | Thermophiles | Alkaliphiles |
| GCA_014201585.1 | *Anoxybacillus tengchongensis* | Thermophiles | Alkaliphiles |
| GCA_017298635.1 | *Alicyclobacillus_B ferrooxydans_B* | Thermophiles | Acidophiles |
| GCA_017310505.1 | *Methylacidiphilum sp004421255* | Thermophiles | Acidophiles |
| GCA_900107035.1 | *Alicyclobacillus hesperidum* | Thermophiles | Acidophiles |
| GCA_900111795.1 | *Anoxybacillus pushchinoensis* | Thermophiles | Alkaliphiles |
| GCA_900116805.1 | *Alicyclobacillus_H macrosporangiidus* | Thermophiles | Acidophiles |
| GCA_900129115.1 | *Thermoanaerobacter uzonensis* | Thermophiles | Acidophiles |
| GCA_900142255.1 | *Alicyclobacillus_I montanus* | Thermophiles | Acidophiles |
| GCA_900156755.1 | *Alicyclobacillus vulcanalis* | Thermophiles | Acidophiles |
| GCA_900176145.1 | *Sulfobacillus thermosulfidooxidans* | Thermophiles | Acidophiles |
| GCA_000195575.1 | *Carnobacterium_A sp000195575* | Psychrophiles | Alkaliphiles |
| GCA_900110375.1 | *Flavobacterium sinopsychrotolerans* | Psychrophiles | Alkaliphiles |
| GCA_018861005.1 | *Polaribacter vadi_A* | Psychrophiles | Alkaliphiles |
| GCA_003259835.1 | *Flavobacterium aquaticum* | Psychrophiles | Alkaliphiles |
| GCA_001761365.1 | *Polaribacter vadi* | Psychrophiles | Alkaliphiles |
| GCA_001975665.1 | *Polaribacter reichenbachii* | Psychrophiles | Alkaliphiles |

### B Intra-genus distance details

The intra-genus distances in the *Temperature dataset* and *pH dataset* were calculated, and the maximum of them for each metric was considered as a threshold for filtering the FCGRs of bacterium-archaeon pairs. Table 4, and Table 5 show the maximum, average, and minimum of intra-genus distances after removing the 5% outliers, for each distance metric in the *Temperature dataset* and *pH dataset*, respectively. Since the values are so close for both datasets, the minimum of the maximum intra-genus values in the two datasets is considered as the threshold.

*Table 4:* Intra-genus distances for the *Temperature dataset*. The table shows the maximum, average, and minimum intra-genus distances for the three distance metrics, DSSIM, Descriptor, and LPIPS.

| Metric | DSSIM | Descriptor | LPIPS |
|---|---|---|---|
| **Max** | 0.517 | 0.274 | 0.503 |
| **Min** | 0.470 | 0.102 | 0.077 |
| **Avg** | 0.493 | 0.151 | 0.129 |

*Table 5:* Intra-genus distances for the *pH dataset*. The table shows the maximum, average, and minimum intra-genus distances for the three distance metrics, DSSIM, Descriptor, and LPIPS.

| Metric | DSSIM | Descriptor | LPIPS |
|---|---|---|---|
| **Max** | 0.51 | 0.31 | 0.66 |
| **Min** | 0.46 | 0.09 | 0.07 |
| **Avg** | 0.49 | 0.15 | 0.14 |

### C Pervasiveness results for the restriction-free scenario

The random genome proxy was tested for classification using SVM models. This classification was repeated 10 times, each time a 10-fold cross-validation with a new random genome proxy was performed to check the pervasiveness of the genomic signature across the genome. The results of this analysis under the restriction-free scenario are provided in Table 6 and Table 7, for the *Temperature Dataset* and *pH Dataset* respectively.

*Table 6:* Maximum average accuracy across six genome proxy lengths in ten repeated SVM classification trials on the *Temperature Dataset* under the restriction-free scenario, for $k$-mer sizes 1 to 9. The table lists the highest average accuracy for each genome proxy length, alongside the $k$-mer size that achieved this accuracy and the variance in percentage.

| Genome proxy length | Class labeling type | Max average accuracy (%) | Variance (%) | $k$-value |
|---|---|---|---|---|
| 10 kbp | Taxonomy | 99.03 | 0.0007 | 5 |
| | Temperature | 81.66 | 0.007 | 5 |
| 50 kbp | Taxonomy | 99.41 | 0.0001 | 6 |
| | Temperature | 84.42 | 0.005 | 7 |
| 100 kbp | Taxonomy | 99.49 | 0.0001 | 7 |
| | Temperature | 85.06 | 0.003 | 7 |
| 250 kbp | Taxonomy | 99.48 | 0.0002 | 7 |
| | Temperature | 85.90 | 0.002 | 9 |
| 500 kbp | Taxonomy | 99.51 | 0.0002 | 7 |
| | Temperature | 86.15 | 0.004 | 9 |
| 1,000 kbp | Taxonomy | 99.51 | 0.0002 | 9 |
| | Temperature | 86.20 | 0.0007 | 9 |

*Table 7:* Maximum average accuracy across six genome proxy lengths in ten repeated SVM classification trials on the *pH Dataset* under the restriction-free scenario, for $k$-mer sizes 1 to 9. The table lists the highest average accuracy for each genome proxy length, alongside the $k$-mer size that achieved this accuracy and the variance in percentage.

| Genome proxy length | Class labeling type | Max average accuracy (%) | Variance (%) | $k$-value |
|---|---|---|---|---|
| 10 kbp | Taxonomy | 97.44 | 0.002 | 4 |
| | pH | 89.80 | 0.014 | 6 |
| 50 kbp | Taxonomy | 97.80 | 0.003 | 7 |
| | pH | 91.40 | 0.006 | 7 |
| 100 kbp | Taxonomy | 98.33 | 0.0008 | 6 |
| | pH | 91.63 | 0.007 | 8 |
| 250 kbp | Taxonomy | 98.32 | 0.002 | 8 |
| | pH | 92.52 | 0.007 | 8 |
| 500 kbp | Taxonomy | 98.54 | 0.001 | 8 |
| | pH | 92.59 | 0.006 | 9 |
| 1,000 kbp | Taxonomy | 92.91 | 0.002 | 9 |
| | pH | 98.43 | 0.002 | 9 |

### D Performance of all classification models across all nine $k$ values

The classification of extremophiles using taxonomic and environment-type labels was done using 6 classifiers, 9 different $k$-mer sizes, and 6 different genome proxy lengths, for both the *Temperature Dataset* and *pH Dataset*, under restriction-free and restricted scenarios.

Figure 1 and Figure 2 present the taxonomy classification accuracy for the *Temperature Dataset* under both restriction-free and restricted scenarios. Similarly, Figure 3 and Figure 4 display the taxonomy classification accuracy for the *pH Dataset* under the same scenarios.

Figure 5 and Figure 6 present the environment-type classification accuracy for the *Temperature Dataset* under both restriction-free and restricted scenarios. Similarly, Figure 7 and Figure 8 display the environment-type classification accuracy for the *pH Dataset* under the same scenarios.

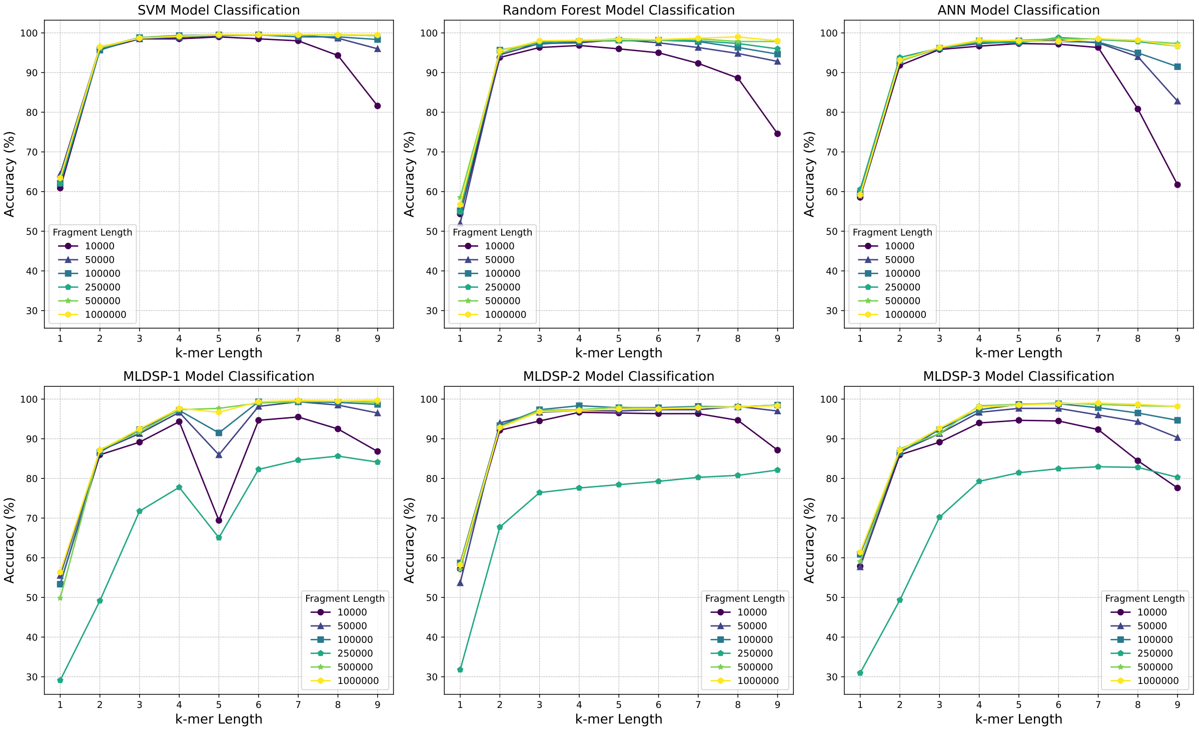


*Figure 1:* Taxonomic classification accuracy of the *Temperature Dataset* under restriction-free scenario.

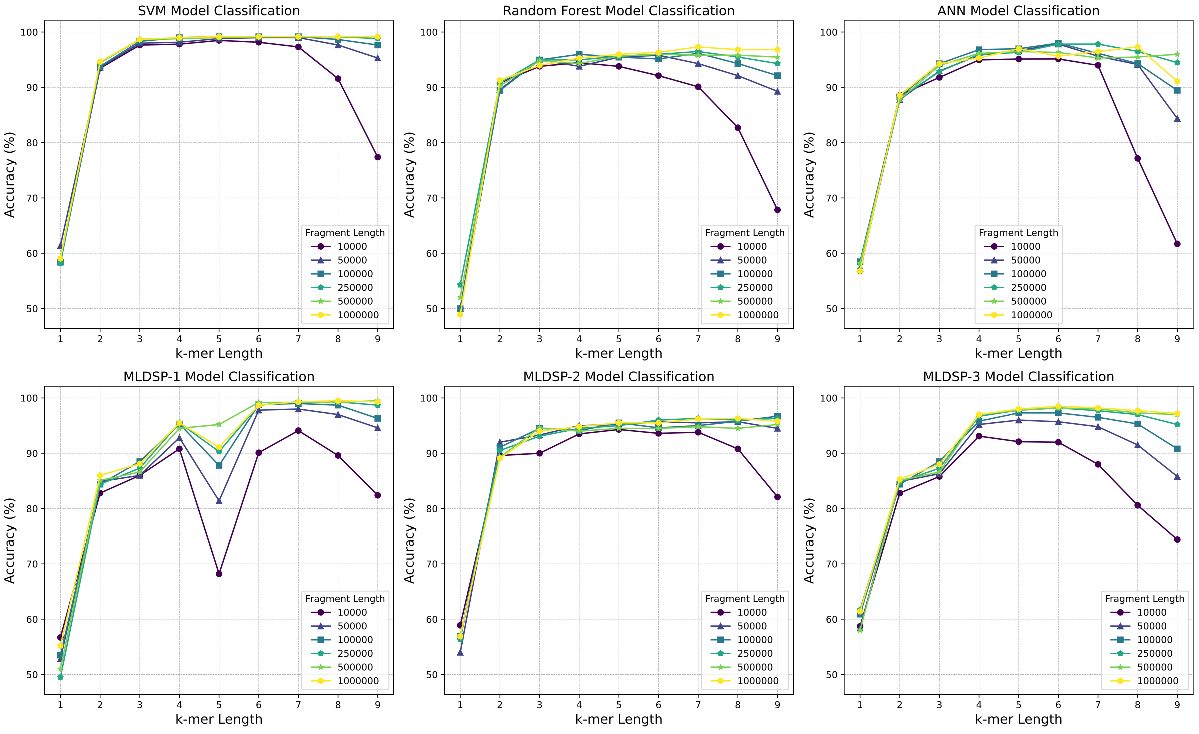


*Figure 2:* Taxonomic classification accuracy of the *Temperature Dataset* under restricted scenario.

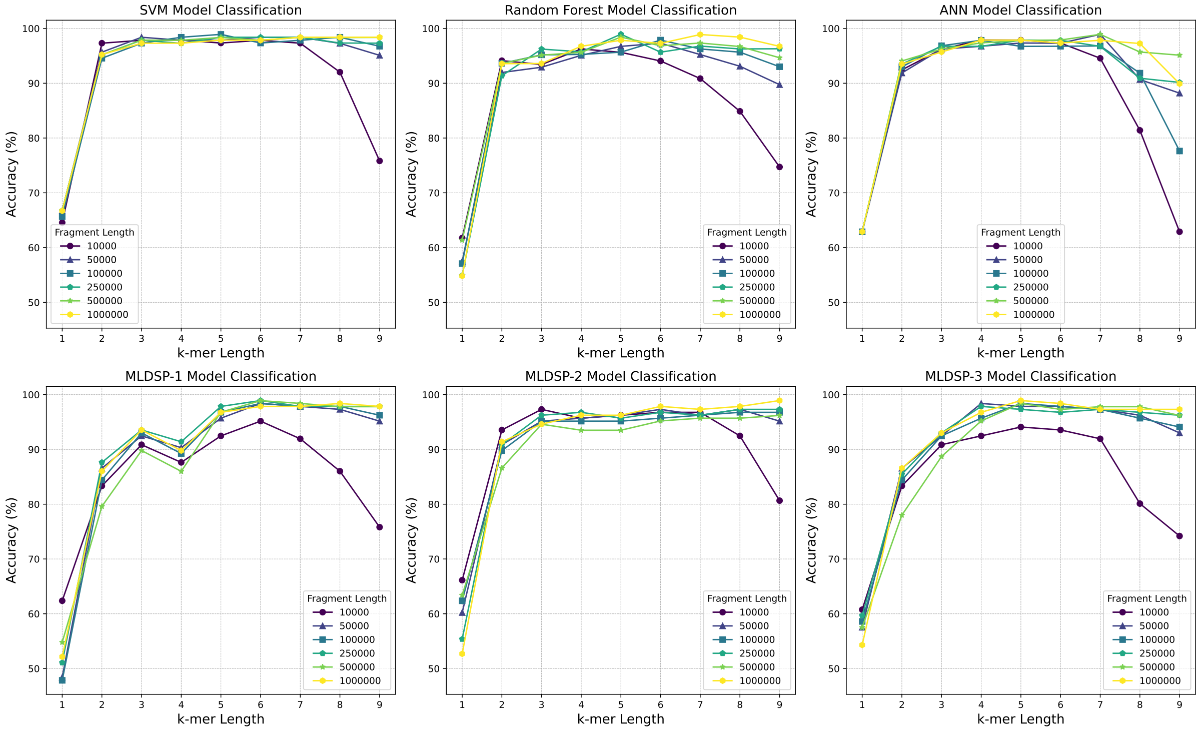


*Figure 3:* Taxonomic classification accuracy of the *pH Dataset* under restriction-free scenario.

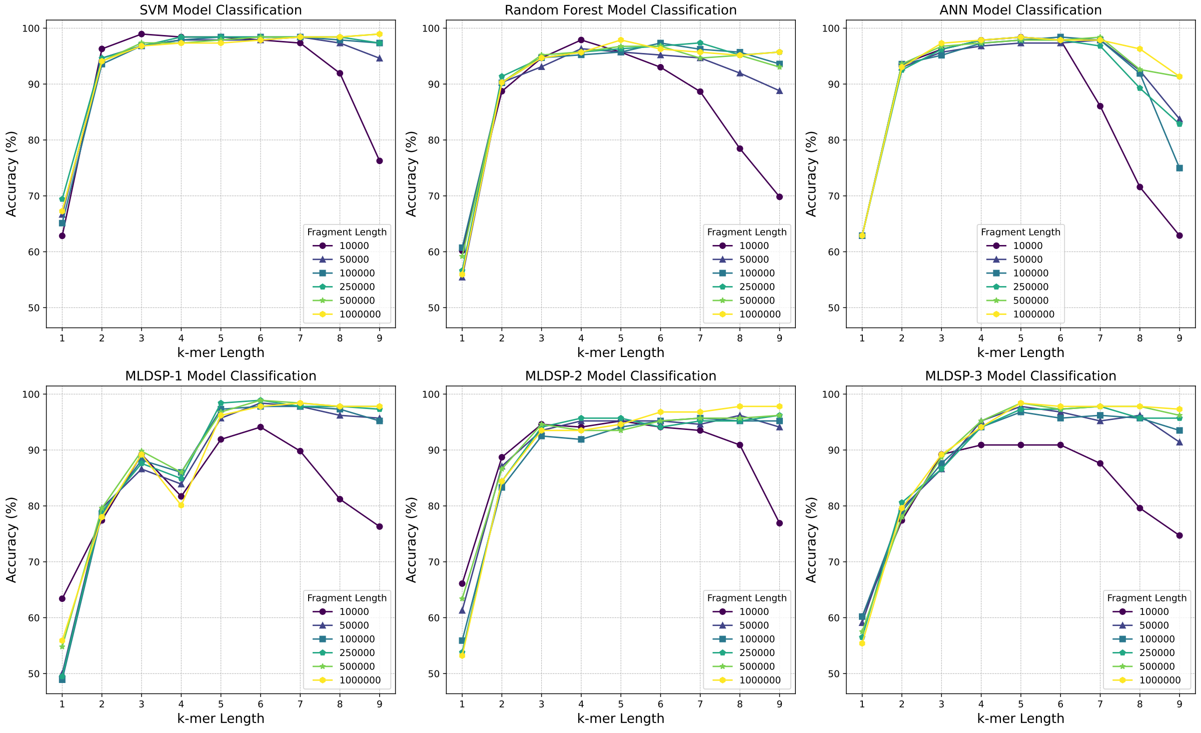


*Figure 4:* Taxonomic classification accuracy of the *pH Dataset* under restricted scenario.

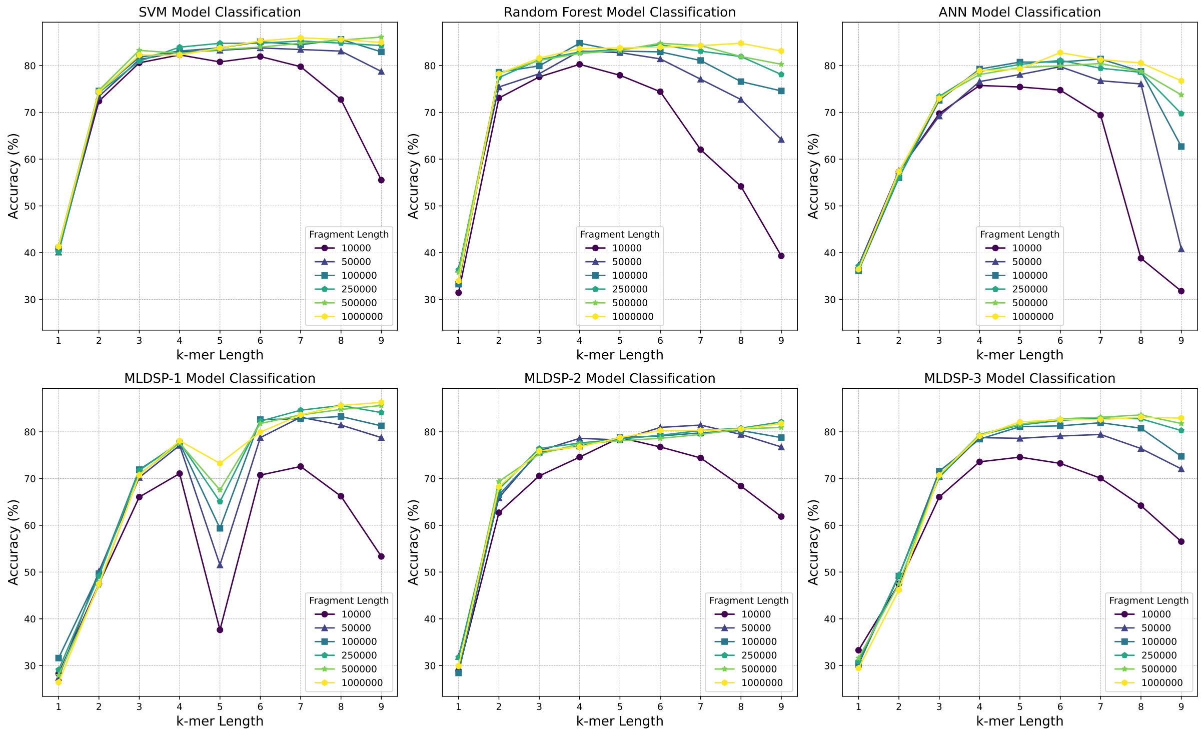


*Figure 5:* Environment-type classification accuracy of the *Temperature Dataset* under restriction-free scenario.

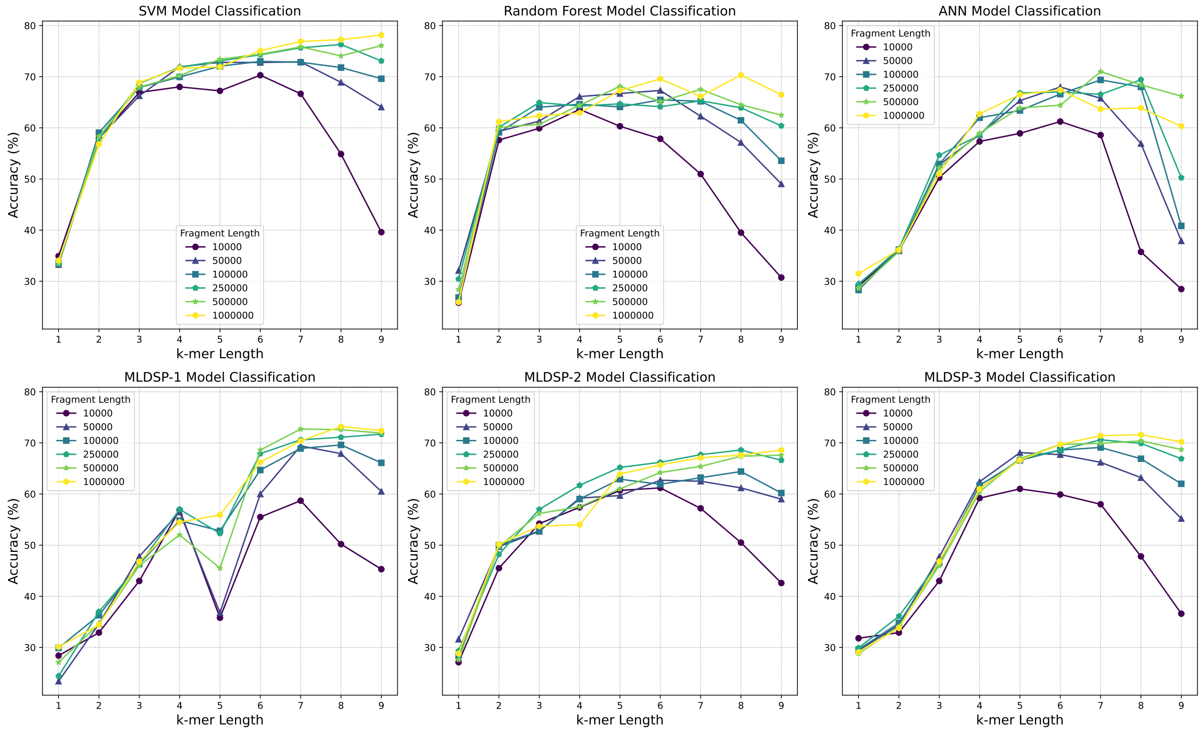


*Figure 6:* Environment-type classification accuracy of the *Temperature Dataset* under restricted scenario.

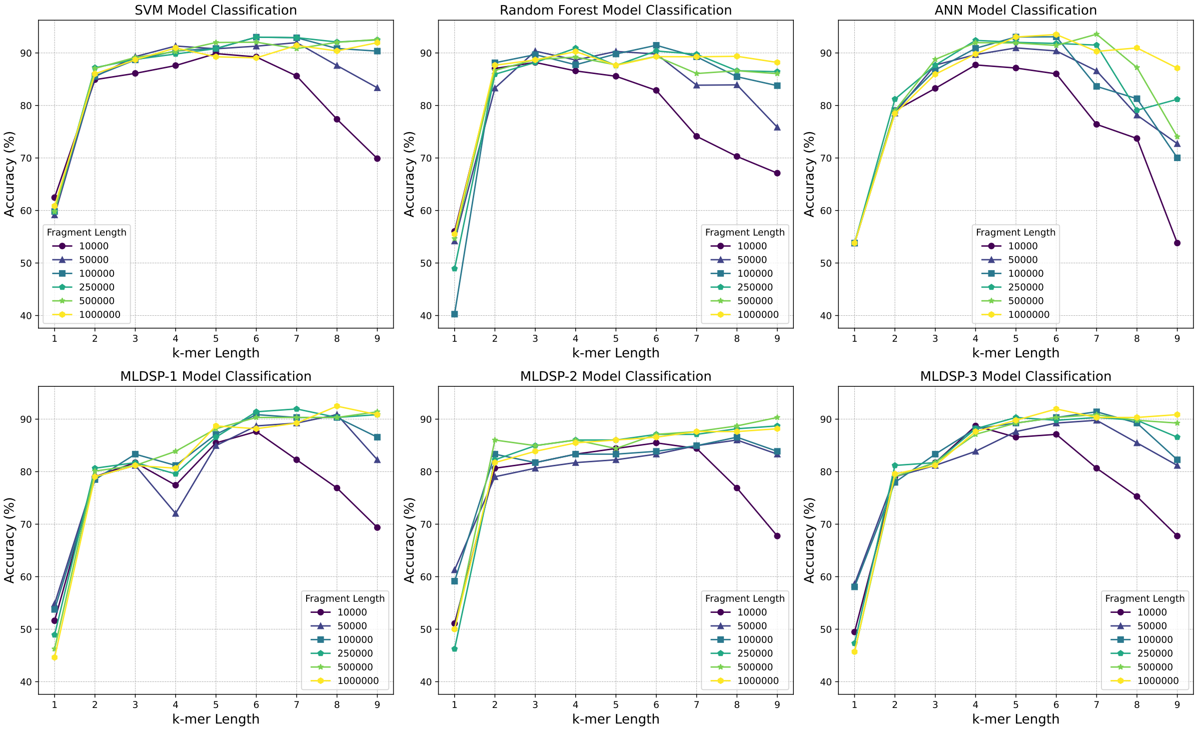


*Figure 7:* Environment-type classification accuracy of the *pH Dataset* under restriction-free scenario.

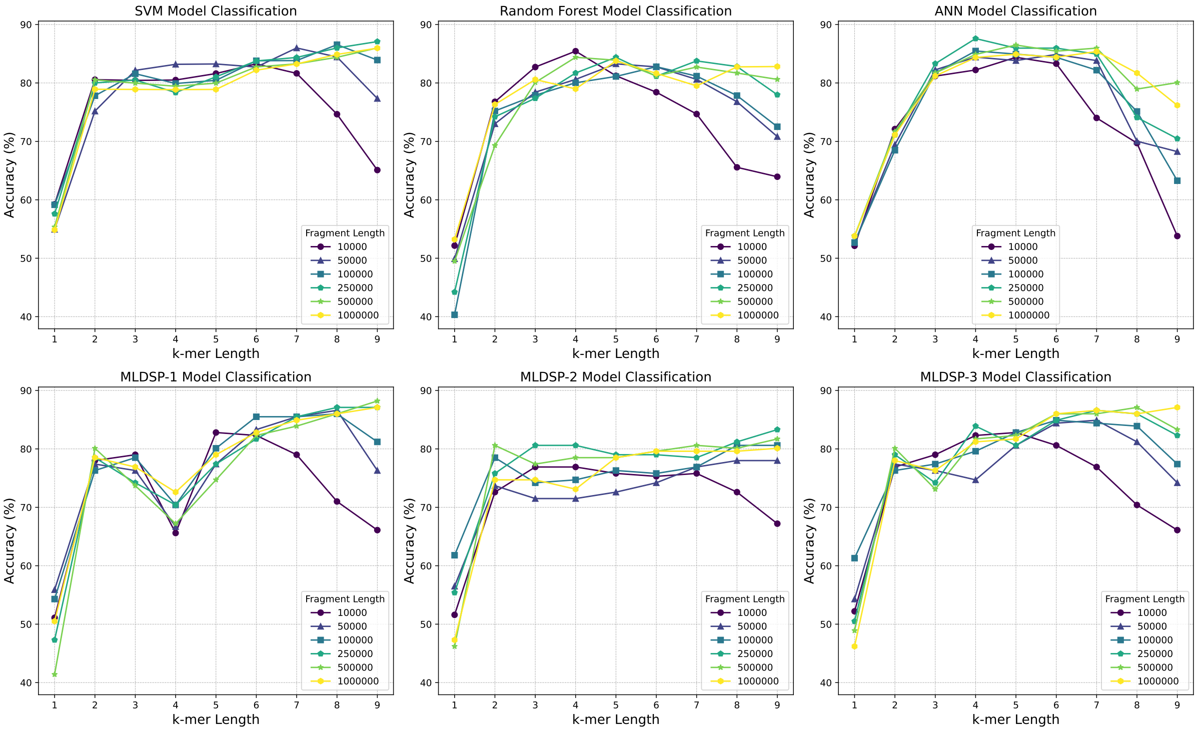


*Figure 8:* Environment-type classification accuracy of the *pH Dataset* under restricted scenario.

### E The detailed results of all classifiers for six genome proxy length and $k$-mer sizes ranging from 6 to 9

The detailed classification accuracy of all six classifiers for all the genome proxy lengths and $k$-mer sizes ranging from 6 to 9 for the restriction-free scenario is provided in Table 8, and Table 10, and for the restricted scenario is provided in Table 9, and Table 11.

*Table 8:* The accuracy of six supervised learning classifiers trained on the *Temperature Dataset*, under the restriction-free scenario, for two different label assignments, taxonomy and environment category, and $k$ values ranging from 6 to 9 and fragment lengths of 10kbp, 50kbp, 100 kbp, 250 kbp, 500 kbp, and 1000 kbp. The classification accuracy was determined through stratified 10-fold cross-validation with non-overlapping genera. The numbers in bold show the highest accuracy for the respective genome proxy length.

| Genome proxy length | k-mer | Labeling type | ANN | RF | SVM | MLDSP-1 | MLDSP-2 | MLDSP-3 |
|---|---|---|---|---|---|---|---|---|
| **10 kbp** | 6 | Temperature | 74.75 | 74.42 | **81.94** | 70.74 | 76.76 | 73.24 |
| | | Taxonomy | 97.15 | 94.99 | **98.50** | 94.65 | 96.32 | 94.48 |
| | 7 | Temperature | 69.40 | 62.03 | 79.77 | 72.58 | 74.41 | 70.07 |
| | | Taxonomy | 96.31 | 92.31 | 98.00 | 95.48 | 96.32 | 92.31 |
| | 8 | Temperature | 38.80 | 54.17 | 72.74 | 66.22 | 68.39 | 64.21 |
| | | Taxonomy | 80.80 | 88.62 | 94.31 | 92.47 | 94.65 | 84.45 |
| | 9 | Temperature | 31.77 | 39.30 | 55.51 | 53.34 | 61.87 | 56.52 |
| | | Taxonomy | 61.71 | 74.57 | 81.59 | 86.79 | 87.12 | 77.59 |
| **50 kbp** | 6 | Temperature | 79.77 | 81.44 | **83.79** | 78.76 | 80.94 | 79.10 |
| | | Taxonomy | 98.16 | 97.50 | **99.50** | 98.16 | 97.32 | 97.66 |
| | 7 | Temperature | 76.76 | 77.09 | 83.44 | 83.11 | 81.44 | 79.43 |
| | | Taxonomy | 97.49 | 96.32 | **99.50** | 99.33 | 97.32 | 95.99 |
| | 8 | Temperature | 76.08 | 72.74 | 83.10 | 81.44 | 79.43 | 76.42 |
| | | Taxonomy | 93.99 | 94.81 | 98.66 | 98.49 | 98.16 | 94.31 |
| | 9 | Temperature | 40.79 | 64.19 | 78.75 | 78.76 | 76.76 | 72.07 |
| | | Taxonomy | 82.79 | 92.81 | 95.99 | 96.49 | 96.99 | 90.30 |
| **100 kbp** | 6 | Temperature | 80.77 | 82.94 | **85.11** | 82.61 | 79.10 | 81.27 |
| | | Taxonomy | 97.83 | 98.16 | **99.50** | 99.33 | 97.83 | 98.83 |
| | 7 | Temperature | 81.41 | 81.10 | 84.45 | 82.78 | 79.77 | 81.94 |
| | | Taxonomy | 97.66 | 97.82 | 98.99 | 99.33 | 98.16 | 97.83 |
| | 8 | Temperature | 78.78 | 76.58 | 85.62 | 83.28 | 80.27 | 80.77 |
| | | Taxonomy | 94.96 | 96.33 | 99.00 | 99.16 | 97.99 | 96.49 |
| | 9 | Temperature | 62.69 | 74.59 | 82.94 | 81.27 | 78.76 | 74.75 |
| | | Taxonomy | 91.48 | 94.65 | 98.33 | 98.66 | 98.49 | 94.65 |
| **250 kbp** | 6 | Temperature | 81.12 | 84.45 | 84.78 | 82.27 | 79.26 | 82.44 |
| | | Taxonomy | 98.83 | 98.16 | 99.49 | 99.00 | 97.66 | 99.00 |
| | 7 | Temperature | 79.44 | 83.10 | 85.28 | 84.62 | 80.27 | 82.94 |
| | | Taxonomy | 98.32 | 98.00 | **99.50** | 99.16 | 98.16 | 98.16 |
| | 8 | Temperature | 78.57 | 81.94 | 84.80 | **85.62** | 80.77 | 82.78 |
| | | Taxonomy | 97.82 | 97.33 | **99.50** | 99.33 | 98.66 | 97.99 |
| | 9 | Temperature | 69.73 | 78.10 | 84.29 | 84.11 | 82.11 | 80.27 |
| | | Taxonomy | 96.66 | 95.97 | 99.33 | 99.16 | 98.83 | 96.49 |
| **500 kbp** | 6 | Temperature | 79.94 | 84.78 | 83.95 | 81.77 | 78.60 | 82.78 |
| | | Taxonomy | 98.50 | 98.17 | **99.50** | 99.00 | 97.66 | 99.00 |
| | 7 | Temperature | 80.45 | 84.27 | 84.78 | 83.61 | 79.43 | 83.11 |
| | | Taxonomy | 98.50 | 98.50 | **99.50** | 99.33 | 97.99 | 98.66 |
| | 8 | Temperature | 78.75 | 81.94 | 85.45 | 84.78 | 80.60 | 83.61 |
| | | Taxonomy | 97.99 | 98.30 | **99.50** | **99.50** | 97.99 | 98.33 |
| | 9 | Temperature | 73.75 | 80.28 | **86.11** | 85.62 | 80.94 | 81.77 |
| | | Taxonomy | 97.32 | 97.83 | 99.33 | 99.16 | 98.49 | 98.16 |
| **1,000 kbp** | 6 | Temperature | 82.77 | 83.94 | 85.28 | 79.93 | 80.27 | 82.61 |
| | | Taxonomy | 97.83 | 98.33 | 99.50 | 99.33 | 97.49 | 98.83 |
| | 7 | Temperature | 81.29 | 84.28 | 85.94 | 83.61 | 80.27 | 82.61 |
| | | Taxonomy | 98.50 | 98.66 | 99.50 | **99.67** | 97.83 | 99.00 |
| | 8 | Temperature | 80.56 | 84.78 | 85.61 | 85.62 | 80.60 | 83.11 |
| | | Taxonomy | 98.16 | 99.00 | 99.49 | 99.50 | 97.99 | 98.66 |
| | 9 | Temperature | 76.75 | 83.13 | 84.95 | **86.29** | 81.77 | 82.94 |
| | | Taxonomy | 96.66 | 98.00 | 99.50 | **99.67** | 98.33 | 98.16 |

*Table 9:* The classification accuracy of six supervised classifiers trained on the *Temperature Dataset*, under the restricted scenario, for two different label assignments, taxonomy and environment category, and $k$ values ranging from 6 to 9 and fragment lengths of 10 kbp, 50kbp, 100 kbp, 250 kbp, 500 kbp, and 1000 kbp. The classification accuracy was determined through stratified 10-fold cross-validation with non-overlapping genera. The numbers in bold show the highest accuracy for the respective genome proxy length.

| Genome proxy length | k-mer | Labeling type | ANN | RF | SVM | MLDSP-1 | MLDSP-2 | MLDSP-3 |
|---|---|---|---|---|---|---|---|---|
| **10 kbp** | 6 | Temperature | 61.23 | 57.85 | **70.29** | 55.50 | 61.20 | 59.90 |
| | | Taxonomy | 95.13 | 92.11 | **98.15** | 90.10 | 93.60 | 92.00 |
| | 7 | Temperature | 58.59 | 50.97 | 66.66 | 58.70 | 57.20 | 58.00 |
| | | Taxonomy | 93.98 | 90.10 | 97.31 | 94.10 | 93.80 | 88.00 |
| | 8 | Temperature | 35.74 | 39.51 | 54.87 | 50.20 | 50.50 | 47.80 |
| | | Taxonomy | 77.15 | 82.71 | 91.58 | 89.60 | 90.80 | 80.60 |
| | 9 | Temperature | 28.47 | 30.73 | 39.60 | 45.30 | 42.60 | 36.60 |
| | | Taxonomy | 61.69 | 67.84 | 77.38 | 82.40 | 82.10 | 74.40 |
| **50 kbp** | 6 | Temperature | 67.98 | 67.31 | 72.78 | 60.00 | 62.70 | 67.70 |
| | | Taxonomy | 97.82 | 95.80 | **98.99** | 97.80 | 95.70 | 95.70 |
| | 7 | Temperature | 65.79 | 62.27 | **72.89** | 69.40 | 62.50 | 66.20 |
| | | Taxonomy | 95.63 | 94.29 | **98.99** | 98.00 | 95.50 | 94.80 |
| | 8 | Temperature | 56.93 | 57.13 | 68.90 | 67.90 | 61.20 | 63.20 |
| | | Taxonomy | 94.13 | 92.11 | 97.65 | 97.00 | 95.70 | 91.50 |
| | 9 | Temperature | 37.89 | 49.03 | 64.06 | 60.50 | 59.00 | 55.20 |
| | | Taxonomy | 84.37 | 89.27 | 95.30 | 94.60 | 94.50 | 85.80 |
| **100 kbp** | 6 | Temperature | 66.60 | 65.47 | **73.00** | 64.70 | 61.90 | 68.60 |
| | | Taxonomy | 97.99 | 95.13 | **99.16** | 98.80 | 94.60 | 97.30 |
| | 7 | Temperature | 69.37 | 65.19 | 72.83 | 68.90 | 63.20 | 69.10 |
| | | Taxonomy | 96.15 | 96.15 | **99.16** | 99.00 | 95.00 | 96.50 |
| | 8 | Temperature | 67.99 | 61.47 | 71.79 | 69.60 | 64.40 | 66.90 |
| | | Taxonomy | 94.30 | 94.30 | 98.65 | 98.70 | 95.80 | 95.30 |
| | 9 | Temperature | 40.83 | 53.56 | 69.61 | 66.10 | 60.20 | 62.00 |
| | | Taxonomy | 89.45 | 92.12 | 97.65 | 96.30 | 96.70 | 90.80 |
| **250 kbp** | 6 | Temperature | 67.01 | 64.13 | **74.29** | 67.90 | 66.20 | 68.60 |
| | | Taxonomy | 97.81 | 95.97 | 99.16 | 98.70 | 96.00 | 98.20 |
| | 7 | Temperature | 66.54 | 65.29 | 75.65 | 70.60 | 67.70 | 70.60 |
| | | Taxonomy | 97.82 | 96.48 | 99.16 | **99.30** | 96.30 | 97.70 |
| | 8 | Temperature | 69.41 | 63.92 | 76.29 | 71.10 | 68.60 | 69.90 |
| | | Taxonomy | 96.49 | 95.48 | 99.16 | **99.30** | 96.00 | 97.00 |
| | 9 | Temperature | 50.24 | 60.39 | 73.09 | 71.70 | 66.60 | 66.90 |
| | | Taxonomy | 94.47 | 94.30 | 98.82 | 98.70 | 96.30 | 95.20 |
| **500 kbp** | 6 | Temperature | 64.42 | 65.11 | 74.40 | 68.60 | 64.20 | 69.70 |
| | | Taxonomy | 96.30 | 95.97 | 99.16 | 99.20 | 94.50 | 98.20 |
| | 7 | Temperature | 70.99 | 67.53 | 75.82 | 72.70 | 65.40 | 69.90 |
| | | Taxonomy | 95.30 | 95.81 | 99.16 | 99.20 | 94.80 | 98.00 |
| | 8 | Temperature | 68.40 | 64.50 | 74.08 | 72.60 | 67.40 | 70.40 |
| | | Taxonomy | 95.48 | 95.81 | 99.16 | 99.20 | 94.50 | 97.30 |
| | 9 | Temperature | 66.20 | 62.48 | **76.08** | 71.90 | 67.60 | 68.70 |
| | | Taxonomy | 95.98 | 95.47 | 99.16 | **99.50** | 95.20 | 97.00 |
| **1,000 kbp** | 6 | Temperature | 67.41 | 69.58 | 75.08 | 66.20 | 65.70 | 69.70 |
| | | Taxonomy | 95.64 | 96.31 | 99.16 | 98.70 | 95.50 | 98.50 |
| | 7 | Temperature | 63.66 | 66.14 | 76.89 | 70.40 | 67.10 | 71.40 |
| | | Taxonomy | 96.48 | 97.31 | 99.16 | 99.30 | 96.20 | 98.20 |
| | 8 | Temperature | 63.88 | 70.31 | 77.25 | 73.20 | 67.60 | 71.60 |
| | | Taxonomy | 97.32 | 96.81 | 99.16 | **99.50** | 96.30 | 97.70 |
| | 9 | Temperature | 60.32 | 66.48 | **78.14** | 72.40 | 68.60 | 70.20 |
| | | Taxonomy | 91.08 | 96.81 | 99.16 | 99.30 | 95.80 | 97.20 |

*Table 10:* The accuracy of six supervised learning classifiers trained on the *pH Dataset*, under the restriction-free scenario, for two different label assignments, taxonomy and environment category, and *k* values ranging from 6 to 9 and fragment lengths of 10kbp, 50kbp, 100 kbp, 250 kbp, 500 kbp, and 1000 kbp. The classification accuracy was determined through stratified 10-fold cross-validation with non-overlapping genera. The numbers in bold show the highest accuracy for the respective fragment length.

| Genome proxy length | k-mer | Labeling type | ANN | RF | SVM | MLDSP-1 | MLDSP-2 | MLDSP-3 |
|---|---|---|---|---|---|---|---|---|
| **10 kbp** | 6 | pH | 86.02 | 82.87 | **89.24** | 87.63 | 85.48 | 87.10 |
| | | Taxonomy | 97.34 | 94.06 | **97.84** | 95.16 | 96.77 | 93.55 |
| | 7 | pH | 76.40 | 74.12 | 85.61 | 82.26 | 84.41 | 80.65 |
| | | Taxonomy | 94.56 | 90.85 | 97.31 | 91.94 | 96.77 | 91.94 |
| | 8 | pH | 73.71 | 70.29 | 77.37 | 76.88 | 76.88 | 75.27 |
| | | Taxonomy | 81.40 | 84.88 | 92.02 | 86.02 | 92.47 | 80.11 |
| | 9 | pH | 53.80 | 67.11 | 69.88 | 69.35 | 67.74 | 67.74 |
| | | Taxonomy | 62.89 | 74.71 | 75.82 | 75.81 | 80.65 | 74.19 |
| **50 kbp** | 6 | pH | 90.38 | 89.80 | 91.29 | 88.71 | 83.33 | 89.25 |
| | | Taxonomy | 97.31 | 97.31 | 98.39 | 98.39 | 97.31 | 97.85 |
| | 7 | pH | 86.58 | 83.86 | **91.96** | 89.25 | 84.95 | 89.78 |
| | | Taxonomy | **98.92** | 95.23 | 98.39 | 97.85 | 96.24 | 97.31 |
| | 8 | pH | 78.16 | 83.92 | 87.60 | 90.86 | 86.02 | 85.48 |
| | | Taxonomy | 90.67 | 93.10 | 97.28 | 97.31 | 97.31 | 96.24 |
| | 9 | pH | 72.72 | 75.85 | 83.36 | 82.26 | 83.33 | 81.18 |
| | | Taxonomy | 88.19 | 89.74 | 95.09 | 95.16 | 95.16 | 93.01 |
| **100 kbp** | 6 | pH | **93.07** | 91.46 | 93.01 | 90.86 | 83.87 | 90.32 |
| | | Taxonomy | 96.75 | 97.87 | 97.34 | **98.39** | 95.70 | 97.85 |
| | 7 | pH | 83.63 | 89.27 | 92.89 | 90.32 | 84.95 | 91.40 |
| | | Taxonomy | 96.81 | 96.26 | 97.84 | 97.85 | 96.24 | 97.31 |
| | 8 | pH | 81.29 | 85.47 | 90.85 | 90.32 | 86.56 | 89.25 |
| | | Taxonomy | 91.84 | 95.67 | **98.39** | 97.85 | 96.77 | 95.70 |
| | 9 | pH | 70.03 | 83.77 | 90.35 | 86.56 | 83.87 | 82.26 |
| | | Taxonomy | 77.63 | 93.01 | 96.75 | 96.24 | 96.77 | 94.09 |
| **250 kbp** | 6 | pH | 91.81 | 90.38 | **93.01** | 91.40 | 87.10 | 89.78 |
| | | Taxonomy | 97.84 | 95.67 | 98.39 | **98.92** | 96.77 | 96.77 |
| | 7 | pH | 91.49 | 89.74 | 92.92 | 91.94 | 87.10 | 90.32 |
| | | Taxonomy | 96.73 | 96.78 | 98.39 | 97.85 | 96.24 | 97.31 |
| | 8 | pH | 79.09 | 86.61 | 92.08 | 90.32 | 88.17 | 89.78 |
| | | Taxonomy | 90.91 | 96.26 | 97.34 | 97.85 | 97.31 | 96.77 |
| | 9 | pH | 81.17 | 86.43 | 92.49 | 90.86 | 88.71 | 86.56 |
| | | Taxonomy | 90.15 | 96.29 | 97.31 | 97.85 | 97.31 | 96.24 |
| **500 kbp** | 6 | pH | 91.43 | 89.36 | 92.05 | 90.32 | 87.10 | 90.32 |
| | | Taxonomy | 97.89 | 96.84 | 97.84 | 98.39 | 97.31 | 97.85 |
| | 7 | pH | **93.57** | 86.08 | 90.82 | 90.32 | 87.63 | 90.86 |
| | | Taxonomy | **98.92** | 97.34 | 98.36 | 98.39 | 97.31 | 97.85 |
| | 8 | pH | 87.22 | 86.61 | 92.02 | 90.32 | 88.71 | 89.78 |
| | | Taxonomy | 95.70 | 96.70 | 98.33 | 97.85 | 97.31 | 97.85 |
| | 9 | pH | 74.04 | 86.05 | 92.57 | 91.40 | 90.32 | 89.25 |
| | | Taxonomy | 95.12 | 94.65 | 98.36 | 97.85 | 97.31 | 96.77 |
| **1,000 kbp** | 6 | pH | **93.51** | 89.30 | 89.06 | 88.17 | 86.56 | 91.94 |
| | | Taxonomy | 97.34 | 97.28 | 97.84 | 97.85 | 97.85 | 98.39 |
| | 7 | pH | 90.29 | 89.27 | 91.43 | 89.25 | 87.63 | 90.32 |
| | | Taxonomy | 97.84 | 98.89 | 98.39 | 97.85 | 97.31 | 97.31 |
| | 8 | pH | 90.96 | 89.36 | 90.38 | 92.47 | 87.63 | 90.32 |
| | | Taxonomy | 97.25 | 98.42 | 98.39 | 98.39 | 97.85 | 97.31 |
| | 9 | pH | 87.11 | 88.19 | 91.96 | 90.86 | 88.17 | 90.86 |
| | | Taxonomy | 89.91 | 96.73 | 98.39 | 97.85 | **98.92** | 97.31 |

*Table 11:* The accuracy of six supervised learning classifiers trained on the *pH Dataset*, under the restricted scenario, for two different label assignments, taxonomy and environment category, and *k* values ranging from 6 to 9 and fragment lengths of 10kbp, 50kbp, 100 kbp, 250 kbp, 500 kbp, and 1000 kbp. The classification accuracy was determined through stratified 10-fold cross-validation with non-overlapping genera. The numbers in bold show the highest accuracy for the respective genome proxy length.

| Genome proxy length | k-value | Labeling type | ANN | RF | SVM | MLDSP-1 | MLDSP-2 | MLDSP-3 |
|---|---|---|---|---|---|---|---|---|
| **10 kbp** | 6 | pH | **83.30** | 78.42 | 83.27 | 82.30 | 75.30 | 80.60 |
| | | Taxonomy | 97.87 | 93.01 | **97.89** | 94.10 | 94.10 | 90.90 |
| | 7 | pH | 74.01 | 74.68 | 81.64 | 79.00 | 75.80 | 76.90 |
| | | Taxonomy | 86.05 | 88.65 | 97.34 | 89.80 | 93.50 | 87.60 |
| | 8 | pH | 69.71 | 65.56 | 74.65 | 71.00 | 72.60 | 70.40 |
| | | Taxonomy | 71.58 | 78.45 | 91.93 | 81.20 | 90.90 | 79.60 |
| | 9 | pH | 53.80 | 63.95 | 65.09 | 66.10 | 67.20 | 66.10 |
| | | Taxonomy | 62.89 | 69.82 | 76.26 | 76.30 | 76.90 | 74.70 |
| **50 kbp** | 6 | pH | 84.88 | 82.75 | 82.72 | 83.30 | 74.20 | 84.40 |
| | | Taxonomy | 97.34 | 95.20 | 97.89 | 98.40 | 95.20 | 96.80 |
| | 7 | pH | 83.83 | 80.61 | 85.99 | 85.50 | 76.90 | 84.90 |
| | | Taxonomy | 97.89 | 94.68 | **98.42** | 97.80 | 94.60 | 95.20 |
| | 8 | pH | 70.03 | 76.78 | 84.44 | **86.60** | 78.00 | 81.20 |
| | | Taxonomy | 92.40 | 91.96 | 97.34 | 96.20 | 96.20 | 96.20 |
| | 9 | pH | 68.25 | 70.82 | 77.34 | 76.30 | 78.00 | 74.20 |
| | | Taxonomy | 83.74 | 88.80 | 94.62 | 95.70 | 94.10 | 91.40 |
| **100 kbp** | 6 | pH | 84.39 | 82.78 | 83.80 | 85.50 | 75.80 | 84.90 |
| | | Taxonomy | **98.42** | 97.34 | **98.42** | 97.80 | 95.20 | 95.70 |
| | 7 | pH | 82.19 | 81.17 | 83.83 | 85.50 | 76.90 | 84.40 |
| | | Taxonomy | 97.87 | 96.23 | **98.42** | 97.80 | 95.70 | 96.20 |
| | 8 | pH | 75.12 | 77.84 | **86.55** | 86.00 | 80.60 | 83.90 |
| | | Taxonomy | 91.90 | 95.70 | 97.89 | 97.30 | 95.20 | 95.70 |
| | 9 | pH | 63.27 | 72.51 | 83.92 | 81.20 | 80.60 | 77.40 |
| | | Taxonomy | 74.97 | 93.60 | 97.34 | 95.20 | 95.20 | 93.50 |
| **250 kbp** | 6 | pH | 85.96 | 81.14 | 83.77 | 81.70 | 79.00 | 84.90 |
| | | Taxonomy | 97.87 | 96.81 | 98.42 | **98.90** | 94.10 | 97.30 |
| | 7 | pH | 84.94 | 83.77 | 84.36 | 85.50 | 78.50 | 86.60 |
| | | Taxonomy | 96.81 | 97.37 | 98.42 | 97.80 | 95.20 | 97.80 |
| | 8 | pH | 74.09 | 82.81 | 85.99 | **87.10** | 81.20 | 86.00 |
| | | Taxonomy | 89.27 | 95.18 | 98.42 | 97.80 | 95.20 | 95.70 |
| | 9 | pH | 70.47 | 77.98 | 87.08 | **87.10** | 83.30 | 82.30 |
| | | Taxonomy | 82.84 | 95.73 | 97.34 | 97.30 | 96.20 | 95.70 |
| **500 kbp** | 6 | pH | 85.44 | 81.20 | 82.75 | 82.30 | 79.60 | 86.00 |
| | | Taxonomy | 97.87 | 96.78 | 98.42 | 98.90 | 95.20 | 97.30 |
| | 7 | pH | 85.96 | 82.72 | 83.27 | 83.90 | 80.60 | 86.00 |
| | | Taxonomy | 98.39 | 94.65 | 98.42 | 98.40 | 95.70 | 97.80 |
| | 8 | pH | 78.95 | 81.70 | 84.39 | 86.00 | 80.10 | 87.10 |
| | | Taxonomy | 92.60 | 95.15 | 98.42 | 97.80 | 95.70 | 97.80 |
| | 9 | pH | 80.06 | 80.61 | 85.99 | **88.20** | 81.70 | 83.30 |
| | | Taxonomy | 91.32 | 93.04 | **98.95** | 97.80 | 96.20 | 96.20 |
| **1,000 kbp** | 6 | pH | 84.39 | 81.64 | 82.19 | 82.80 | 79.60 | 86.00 |
| | | Taxonomy | 97.87 | 96.26 | 97.87 | 97.80 | 96.80 | 97.80 |
| | 7 | pH | 85.38 | 79.50 | 83.25 | 84.90 | 79.60 | 86.60 |
| | | Taxonomy | 97.87 | 95.70 | 98.42 | 98.40 | 96.80 | 97.80 |
| | 8 | pH | 81.70 | 82.75 | 84.88 | 86.00 | 79.60 | 86.00 |
| | | Taxonomy | 96.29 | 95.15 | 98.42 | 97.80 | 97.80 | 97.80 |
| | 9 | pH | 76.17 | 82.81 | 85.96 | **87.10** | 80.10 | **87.10** |
| | | Taxonomy | 91.35 | 95.73 | **98.95** | 97.80 | 97.80 | 97.30 |

### F Phenotypic traits of final pairs

In our study, we studied the phenotypic traits of the species of the confirmed pairs groups, highlighting key attributes such as pH range, temperature, salinity, and cell shape. The Tables Table 12, Table 13, Table 14, Table 15, and Table 16 show the details of this study for Groups 1 to 5, respectively. The tables also include geochemical details of the environments from which the microbes were isolated.

*Table 12:* Characterizing phenotypic traits of species in Group 1 and their isolating environment. The table describes various phenotypic traits attributed to each of the species and the geochemical information associated with the environments the microbes were initially isolated. The optimized growth range(s), if known, are described in parentheses. Phenotypic trait categories lacking information for a given species are denoted with "—".

| **Species** | ***Thermoanaerobacterium thermosaccharolyticum*** [23, 24] | ***Caldisphaera lagunensis*** [18] |
|---|---|---|
| **Domain** | Bacteria | Archaea |
| **pH range** | 4.1-7.6 (5-5.25) | 2.3-5.4 (3.75) |
| **Temperature** | 60 °C | 45-80 °C (72.5°C) |
| **Salinity** | 0-2.5% NaCl | 0-1.5% NaCl |
| **Cell shape** | Long slender granulated bacilli | Mostly regular cocci |
| **Gram stain** | Positive | Negative |
| **Motility** | Peritrichous flagella | Non-motile |
| **S-Layer protein composition** | S-layer lattice | P3-Symmetry layer lattice |
| **Oxygen tolerance** | Anaerobic | Anaerobic |
| **Genome size** | 2.8 Mb | 1.5 Mb |
| **Intergenic sequence content** | Pseudogenes present | Pseudogenes present |
| **Geographic source** | Derived from Austrian beet sugar factory | Derived from Mud Spring Mt Maquiling Laguna Philippines |
| **Geochemical parameters** | Observed in geothermal hot springs | Observed in volcanic acidic hot springs |

*Table 13:* Characterizing phenotypic traits of species in Group 2 and their isolating environment. The table describes various phenotypic traits attributed to each of the species, and the geochemical information associated with the environments the microbes were initially isolated. The optimized growth range(s), if known, are described in parentheses. Phenotypic trait categories lacking information for a given species are denoted with “—".

| **Species** | ***Thermotoga petrophila***[42] | ***Geoglobus acetivorans***[39] |
|---|---|---|
| **Domain** | Bacteria | Archaea |
| **pH range** | 5.2-9.0 (7.0) | 5.0-7.5 (6.8) |
| **Temperature** | 47-88 °C (80 °C (81 °C) | |
| **Salinity** | 0.1-5.5% NaCl (1.0% NaCl) | 1.0-6.0% NaCl (2.5% NaCl) |
| **Cell shape** | Rods (bacilli) | Regular to irregular cocci |
| **Gram stain** | Negative | Negative |
| **Motility** | Subpolar and lateral flagella | Non-motile |
| **S-Layer protein composition** | S-layer lattice | S-layer lattice |
| **Oxygen tolerance** | Anaerobic | Anaerobic |
| **Genome Size** | 1.8 Mb | 1.9 Mb |
| **Intergenic sequence ccontent** | Pseudogenes present | Pseudogenes present |
| **Geographic source** | Isolated from production fluid in the Kubiki oil reservoir, Niigata prefecture, Japan | Isolated from Ashadze hydrothermal field, Mid-Atlantic Ridge |
| **Geochemical parameters** | Subterranean starved conditions | Black smoker field at depth of 4100 m |

*Table 14:* Characterizing phenotypic traits of species in Group 3 and their isolating environment. The table describes various phenotypic traits attributed to each of the species and the geochemical information associated with the environments the microbes were initially isolated. The optimized growth range(s), if known, are described in parentheses. Phenotypic trait categories lacking information for a given species are denoted with "—".

| **Species** | ***Thermocrinis ruber***[9, 16] | ***Pyrococcus furiosus***[6, 13] | ***Thermofilum adornatum***[12, 44] | ***Paleococcus pacificus***[46] | ***Pyrococcus chitonophagus***[17, 3] | ***Thermococcus litoralis***[28, 29] |
|---|---|---|---|---|---|---|
| **Domain** | Bacteria | Archaea | Archaea | Archaea | Archaea | Archaea |
| **pH range** | 7.0-8.5 | 5.0-9.0 | 5.3-8.5 | 5.0-8.0 | 3.5-9 | 6.0-8.5 |
| **Temperature** | 44-89 °C (80°C) | 70-100 °C (93 °C) | 50-95 °C (80 °C) | 50-90 °C (80 °C) | 60-93 °C (85°C) | 55-98 °C (88 °C) |
| **Salinity** | 0-0.4% NaCl | 0.5-5% NaCl | 0-2.5% NaCl | 1-4% NaCl (3% NaCl) | 1.8-6.5% NaCl (2.5% NaCl) | 1.8-6.5% NaCl (2.5% NaCl) |
| **Cell shape** | Rod-shaped cells | Slightly irregular cocci | Filamentous bacilli | Irregular cocci | Round to irregular cocci | Round to irregular cocci |
| **Gram stain** | Negative | Negative | Negative | Negative | Negative | Negative |
| **Motility** | Monopolar polytrichous flagella | Monopolar polytrichous flagella | Monopolar polytrichous flagella | Monopolar polytrichous flagella | Monopolar polytrichous flagella | Non-flagellated, non-motile |
| **S-Layer protein composition** | No evidence of a regularly arrayed SLP | Hexagonal lattice | — | — | Hexagonal lattice | Hexagonal lattice |
| **Oxygen tolerance** | Aerobic | Anaerobic | Anaerobic | Anaerobic | Anaerobic | Anaerobic |
| **Genome size** | 1.52 Mb | 1.89 Mb | 1.75 Mb | 1.9 Mb | 1.95 Mb | 1.82 Mb |
| **Intergenic sequence content** | Pseudogenes present | Pseudogenes and IS elements present | No pseudogenes | No pseudogenes | Pseudogenes and IS elements present | Pseudogenes and IS elements present |
| **Geographic source** | Octopus Spring Yellowstone National Park WY USA | Submarine solfataric field in the bay of Porto di Levante Vulcano Island Italy | Isolated from a Kamchatkan (Siberian) hot spring in 2009 | Isolated from a deep-sea hydrothermal vent field at a depth of 2737 m at the Niaochao site on the East Pacific Ocean Rise | Smoker Site Guaymas Basin Gulf of California Mexico | Shallow submarine solfataras near the beach of Lucrino Bay of Naples, Italy |
| **Geochemical parameters** | Evidence for hydrothermal petroleum-like substances in other hot springs atop the caldera | Evidence for gas discharges containing light hydrocarbons | Solfataric hot spring (sulphur-containing gas exhaust) | Geothermally heated marine sediments 2773 m | Evidence for gas discharges containing petroleum-like hydrocarbons | Evidence for gas discharges containing light hydrocarbons |

*Table 15:* Characterizing phenotypic traits of species in Group 4 and their isolating environment. The table describes various phenotypic traits attributed to each of the species, and the geochemical information associated with the environments the microbes were initially isolated. The optimized growth range(s), if known, are described in parentheses. Phenotypic trait categories lacking information for a given species are denoted with "—".

| **Species** | ***Pseudothermotoga_B elfii***[32] | ***Methanobacterium_C paludis***[5] | ***Methanosarcina vacuolata***[47] |
|---|---|---|---|
| **Domain** | Bacteria | Archaea | Archaea |
| **pH range** | 5.5–7.5 | 4.8–6.6 | 6.0–8.0 |
| **Temperature** | 66°C | 16–40°C | 18–42°C |
| **Salinity** | 0.0–2.8% NaCl | — | — |
| **Cell shape** | Regular bacilli | Regular bacilli | Irregular cocci |
| **Gram stain** | Negative | Negative | Positive |
| **Motility** | Peritrichous flagella | Non-motile | Non-motile |
| **S-Layer protein composition** | SLPs present | — | — |
| **Oxygen tolerance** | Anaerobic | Anaerobic | Anaerobic |
| **Genome size** | 2.2 Mb | 2.5 Mb | 4.6 Mb |
| **Intergenic sequence content** | Pseudogenes present | Pseudogenes present | Pseudogenes present |
| **Geographic source** | Isolated from an oil-producing well in Africa | Isolated from peat soil near Anchorage, Alaska | Isolated from an anaerobic digestor in the former USSR |
| **Geochemical parameters** | Deep subsurface environment | Described as "hydrogenotrophic, methanogen(ic)" | Initially found in sludge of methane tank; also found in wetlands and swamps |

*Table 16:* Characterizing phenotypic traits of species in Group 5 and their isolating environment. The table describes phenotypic traits attributed to each species and the geochemical information associated with the environments from which the microbes were initially isolated. Optimized growth ranges, if known, are shown in the pH, temperature, and salinity rows. Phenotypic trait categories lacking information for a given species are denoted with "—".

| **Species** | ***Rubrobacter indicoceani***[8] | ***Methanoculleus marisnigri***[1, 2] | ***Methanoculleus bourgensis***[25, 11] | ***Methanolinea mesophila***[36] | ***Methanoculleus horonobensis***[38] | ***Methanoculleus taiwanensis***[43] | ***Methanoculleus thermophilus***[27, 22] |
|---|---|---|---|---|---|---|---|
| **Domain** | Bacteria | Archaea | Archaea | Archaea | Archaea | Archaea | Archaea |
| **pH range** | 7.0–8.0 | 6.0–7.5 | 6.8–7.0 | 6.5–7.4 | 5.8–8.2 | 8.1 | — |
| **Temperature** | 20–37°C | 20–25°C | 37°C | 20–40°C | 25–45°C | 37°C | 55°C |
| **Salinity** | 1.0–5.0% NaCl | 0.0–0.7% NaCl | — | 0.0–0.025% NaCl | 0.0–1.3% NaCl | 0.0–0.1% NaCl | — |
| **Cell shape** | Short bacilli | Irregular cocci | — | Filamentous cocci | Irregular cocci | Irregular cocci | Irregular cocci |
| **Gram stain** | Positive | Negative | Negative | Negative | Negative | Negative | Negative |
| **Motility** | Non-motile | Peritrichous flagella | — | Non-motile | Non-motile | Non-motile | — |
| **S-Layer protein composition** | — | Hexagonal SLPs | — | — | — | SLPs present | — |
| **Oxygen tolerance** | Aerobic | Anaerobic | Anaerobic | Anaerobic | Anaerobic | Anaerobic | Anaerobic |
| **Genome size** | 3.07 Mb | 2.48 Mb | 2.79 Mb | 2.7 Mb | 2.4 Mb | 2.8 Mb | 2.2 Mb |
| **Intergenic sequence content** | Pseudogenes present | Pseudogenes present | Pseudogenes present | Pseudogenes present | Pseudogenes present | Pseudogenes present | Pseudogenes present |
| **Geographic source** | Deep-sea sediment, Indian Ocean | Sediment and anaerobic digestors | Sewage, sludge digestor | Rice field soil, Taiwan | Groundwater sampled from a diatomaceous shale formation in Horonobe, Japan | Deep-sea sediment sourced off the coast of Taiwan | Sediment under nuclear power plant |
| **Geochemical parameters** | 4602m depth (0.006154° N, 81.031163° E) | Methanogenic, anoxic settlements from the Black Sea | Methanogenic, found in high ammonia and high salt biogas-synthesizing digestors | Hydrogenotrophic, methanogenic | Hydrogenotrophic, methanogenic | Hydrogenotrophic, methanogenic | Hydrogenotrophic, methanogenic |

### G Final pairs FCGRs

Our study employed a three-layer method to identify microbial pairs that share similar genomic signatures despite originating from maximally divergent domains. The final set includes 15 bacterium-archaeaon pairs. The FCGRs of these pairs are shown in Figures Figure 9 and Figure 10. Figure Figure 9 illustrates Groups 1 to 3, all of which inhabit extreme environments, while Figure Figure 10 shows Groups 4 and 5, which mainly consist of mesophiles.

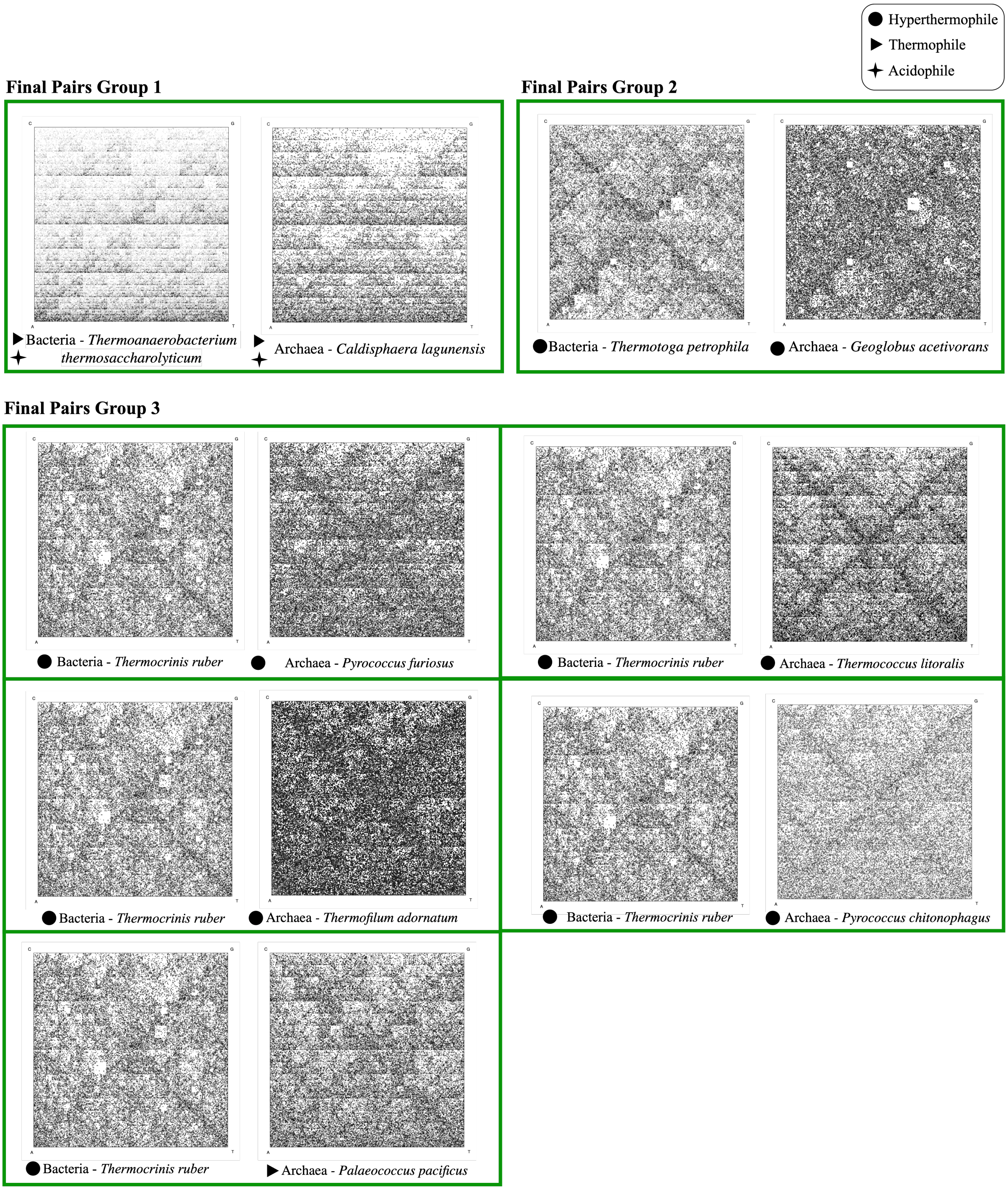


*Figure 9:* The FCGRs of all pairs in Groups 1, 2, and 3. We used $k = 8$ to generate the FCGRs.

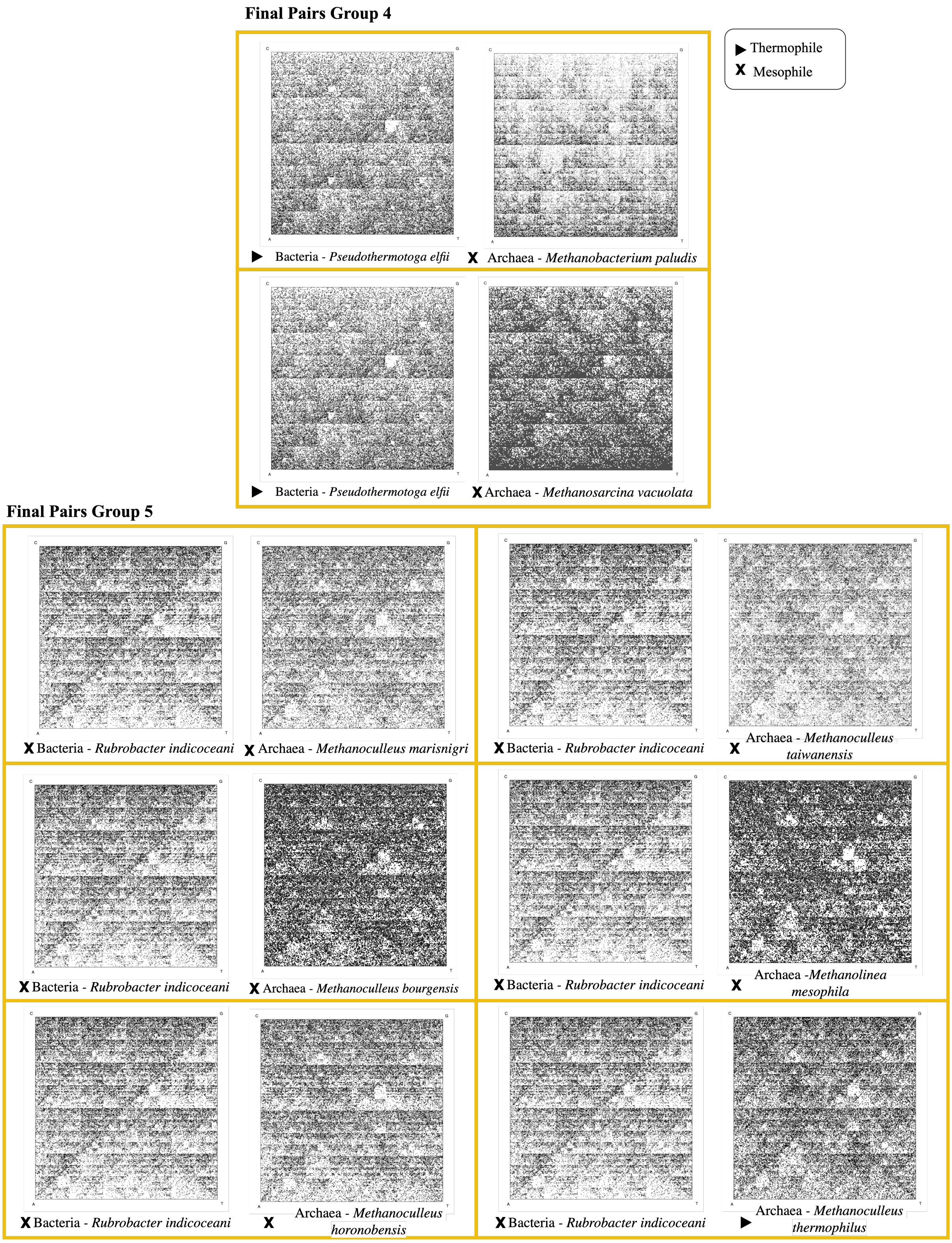


*Figure 10:* The FCGRs of all pairs in final pairs group 4 and 5. We used $k = 8$ to generate the FCGRs.

### H Analysis of 3-mer profiles of confirmed bacterium-archaeon pairs

To investigate potential biases in 3-mer usage related to environmental adaptation, we performed a detailed analysis of the 3-mer profiles for the genome proxies of the organisms in the “confirmed bacterium–archaeon pairs" groups. The focus was on $k = 3$, due to its biological significance, as codons form a subset of the set of 3-mers. Following the four-step analysis, this section examines how the 3-mer frequencies reflect environmental adaptations across taxonomically divergent microbes. The results of this comprehensive analysis are discussed separately below, for each of the five "confirmed bacterium–archaeon pairs" groups.

**Confirmed Pairs Group 1 (Compelling).** This group consists of two thermophilic acidophiles: *T. thermosaccharolyticum*, (bacterium), and *C. lagunensis* (archaeon). As detailed below, all four steps of the 3-mer analysis concurred in showing remarkable genomic signature similarities between the two organisms.

3-mer deviation analysis: As illustrated in Figure 11, both organisms exhibited similar patterns of 3-mer over- and under-representation, relative to the dataset average.

Environment-relevant 3-mers identification: Figure 11 shows that the two species had 10 environment-relevant 3-mers in common with similar over- and under-representation, out of a total of 15 environment-relevant 3-mers identified for each organism.

Correlation assessment: As shown in Figure 12, the similarity in 3-mer profiles was quantitatively confirmed by a Spearman's rank correlation coefficient of 0.96 with $p$-value $< 10^{-5}$ between the 3-mer counts of the two organisms. This indicates a statistically significant correlation in the 3-mer composition of the two genomic signatures.

Comparison with literature: As shown in Table 17, our findings in the 3-mer bias patterns in *C. lagunensis* showed nearly complete agreement with previously reported findings in the biology literature, with 12 out of 15 3-mers showing agreement with the literature. Similarly, our findings for *T. thermosaccharolyticum* showed strong agreement with published codon usage and amino acid abundance patterns, with 12 of 15 environment-relevant 3-mers agreeing with established findings. The comparison between the two species also revealed 9 3-mers with similar frequencies in both genomes, which also aligned with published findings in the literature.

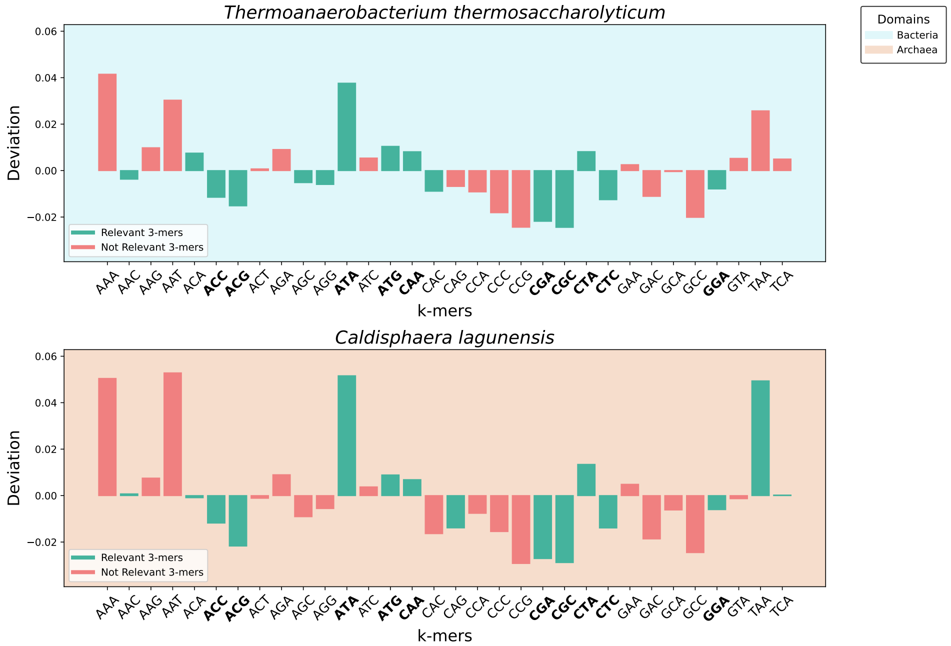


*Figure 11:* 3-mer usage bias analysis of the Confirmed Pairs Group 1, which includes two species: one bacterium, *Thermoanaerobacterium thermosaccharolyticum*, and one archaeon, *Caldisphaera lagunensis*. The top panel displays the deviation of each 3-mer in *T. thermosaccharolyticum* relative to its average frequency within both *Temperature Dataset* and *pH Dataset*, while the bottom panel shows the same analysis for *C. lagunensis*. Green bars indicate 3-mers identified as relevant to the environment-type classification for that species, whereas red bars represent 3-mers that did not influence that classification. Only canonical 3-mers are considered in this analysis. Shared environment-relevant 3-mers with similar over- and under-representation are indicated in boldface.

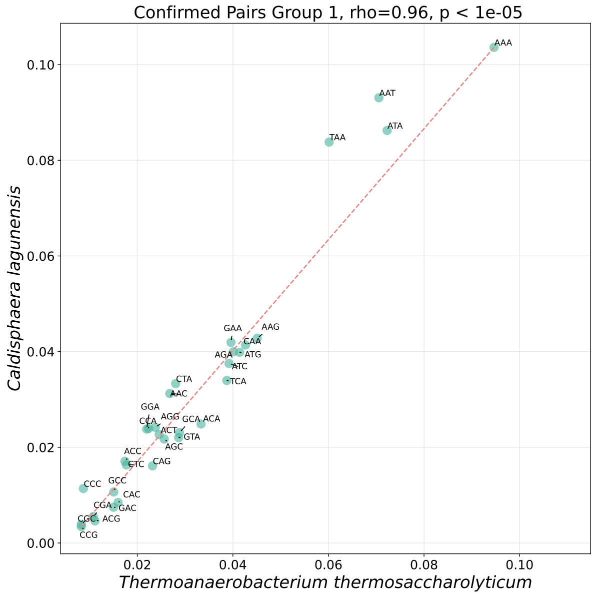


*Figure 12:* Correlation analysis of 3-mer usage between the bacterium *Thermoanaerobacterium thermosaccharolyticum* and the archaeon *Caldisphaera lagunensis*, pair-members from the Confirmed Pairs Group 1. Each point represents a specific 3-mer, with its normalized frequency in *T. thermosaccharolyticum* plotted on the $x$-axis and its frequency in *C. lagunensis* on the $y$-axis. The analysis reveals a strong positive Spearman's rank correlation coefficient, $\rho = 0.96$ with $p$-value $< 10^{-5}$, between the 3-mer usage patterns of the two organisms.

*Table 17:* Over- and under-representation of the relevant 3-mers, found by our method to be collectively associated with genomic signatures of extremophiles found in Group 1. The symbol ↑ (↓) indicates over-representation (under-representation) of a 3-mer/codon. Matched arrows, e.g., (↑, ↑) indicate that both our method and the literature agree in their finding. Mismatched arrows indicate disagreement. In cases where the letter "X" is found in place of a secondary arrow indicates a lack of supporting literature. The 3-mers with complete matching patterns between the bacteria and the archaea in this group are highlighted in bold.

| **Domain** | Bacteria | Archaea |
|---|---|---|
| **Species** | *Thermoanaerobacterium thermosaccharolyticum* | *Caldisphaera lagunensis* |
| **Temperature Label** | Thermophiles | Thermophiles |
| **pH Label** | Acidophiles | Acidophiles |
| **Corresponding Amino Acid** | **Representation of 3-mers** | |
| Arg | AGG (↓, ↓)<br>**CGC (↓, ↓)**<br>**CGA (↓, ↓)** | **CGA (↓, ↓)**<br>**CGC (↓, ↓)** |
| Asn | AAC (↓, ↑) | AAC (↑, ↑) |
| Gln | **CAA (↑, ↑)** | **CAA (↑, ↑)**<br>CAG (↓, ↓) |
| Gly | GGA (↓, ↑) | GGA (↓, ↓) |
| His | CAC (↓, ↓) | - |
| Ile | **ATA (↑, ↑)** | **ATA (↑, ↑)** |
| Leu | **CTA (↑, ↑)**<br>**CTC (↓, ↓)** | **CTA (↑, ↑)**<br>**CTC (↓, ↓)** |
| Met | ATG (↑, X) | ATG (↑, X) |
| Ser | AGC (↓, ↓) | TCA (↑, ↓) |
| STOP | - | TAA (↑, X) |
| Thr | **ACA (↑, ↑)**<br>**ACC (↓, ↓)**<br>**ACG (↓, ↓)** | **ACA (↑, ↑)**<br>**ACC (↓, ↓)**<br>**ACG (↓, ↓)** |

**Confirmed Pairs Group 2 (strong).** This group consists of one pair of hyperthermophiles, *T. petrophila* (bacterium), and *G. acetivorans* (archaeon). For this group, three out of the four steps of the analysis showed remarkable genomic signature similarities between the two organisms, with the fourth indicating a moderate level of agreement with biological literature.

3-mer deviation analysis: As illustrated in Figure 13, similar to Group 1, both organisms displayed remarkable convergence in their patterns of over- or under-representation relative to the dataset average.

Environment-relevant 3-mers identification: Feature importance analysis revealed that the two organisms had in common eight environment-relevant 3-mers with similar over- and under-representation patterns.

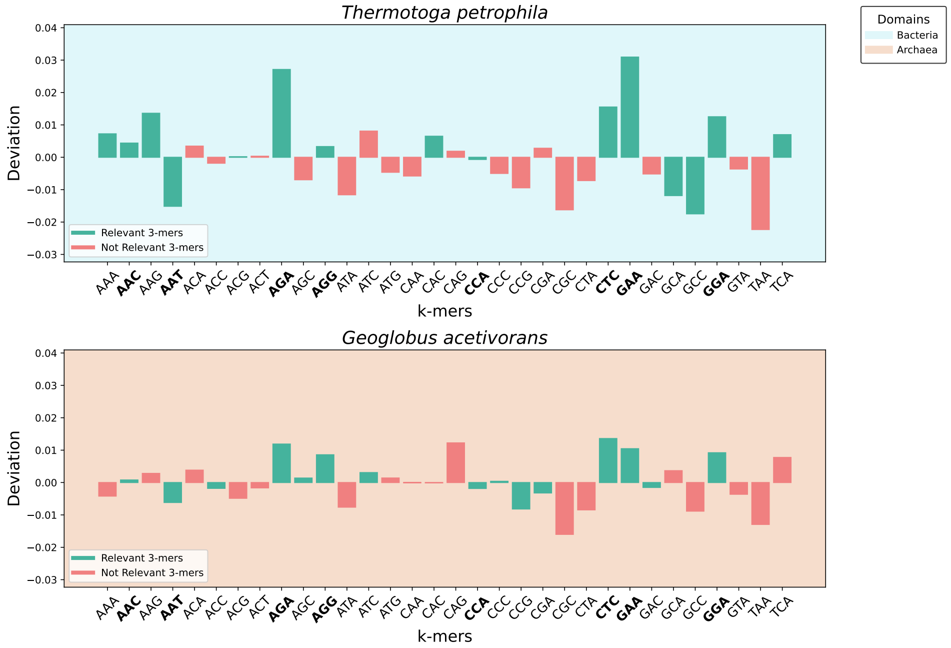


*Figure 13:* 3-mer usage bias analysis of the Confirmed Pairs Group 2, which includes two species: one bacterium, *T. petrophila*, and one archaeon, *G. acetivorans*. The top panel displays the deviation of each 3-mer in *T. petrophila* relative to its average frequency within *Temperature Dataset*, while the bottom panel shows the same analysis for *G. acetivorans*. Green bars indicate 3-mers identified as relevant to the environment-type classification for that species, whereas red bars represent 3-mers that did not influence that classification. Only canonical 3-mers are considered in this analysis. Shared environment-relevant 3-mers with similar over- and under-representation patterns are indicated in boldface.

Correlation assessment: As shown in Figure 14, the similarity in 3-mer frequency profiles was quantitatively confirmed by a Spearman's rank correlation coefficient of 0.81 with $p$-value $< 10^{-5}$ between the 3-mer counts of the two organisms.

Comparison with literature: As shown in Table 18, our findings revealed that for *T. petrophila*, seven out of 15 3-mers were either over- or under-represented in a pattern consistent with previously reported patterns in the literature. Similarly, for *G. acetivorans*, eight out of 15 3-mers showed agreement with known bias trends in the literature. Overall, there was a moderate level of agreement between the 3-mer bias patterns identified by our method and the codon usage and amino acid abundance biases reported in other studies. Additionally, the comparison between the two species revealed four 3-mers with similar frequencies across both genomes, further supporting observations previously documented in the literature.

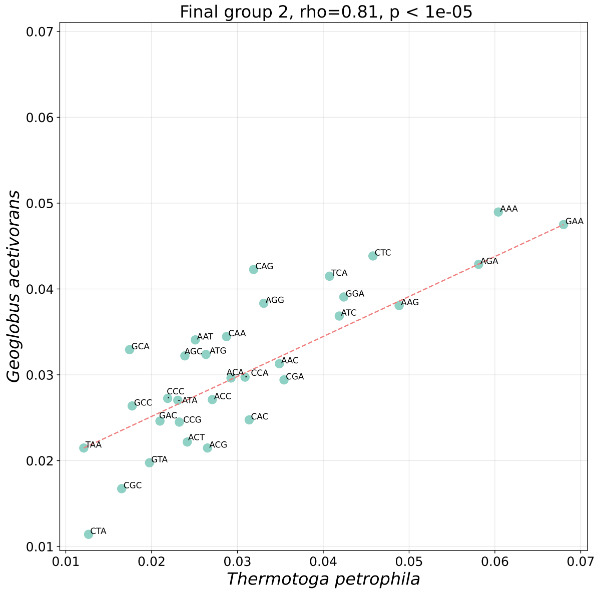


*Figure 14:* Correlation analysis of 3-mer usage between the bacterium *T. petrophila* and the archaeon *G. acetivorans*, pair, members from the Confirmed Pairs Group 2. Each point represents a specific 3-mer, with its normalized frequency in *T. petrophila* plotted on the $x$-axis and its frequency in *G. acetivorans* on the $y$-axis. The analysis reveals a strong positive Spearman's rank correlation coefficient, $\rho = 0.81$ with $p$-value $< 10^{-5}$, between the 3-mer usage patterns of the two organisms.

*Table 18:* Over- and under-representation of the relevant 3-mers, found by our method to be collectively associated with genomic signatures of extremophiles found in the Confirmed Pairs Group 2. The symbol ↑ (↓) indicates over-representation (under-representation) of a 3-mer/codon. Matched arrows, e.g., (↑, ↑) indicate that both our method and literature agree in their finding. Mismatched arrows indicate disagreement. In cases where the letter "X" is found in place of a secondary arrow indicates a lack of supporting literature. The 3-mers with complete matching patterns between the bacteria and the archaea in this group are highlighted in bold.

| **Domain** | Bacteria | Archaea |
|---|---|---|
| **Species** | *Thermotoga petrophila* | *Geoglobus acetivorans* |
| **Temperature Label** | Hyperthermophiles | Hyperthermophiles |
| **Corresponding Amino Acid** | **Representation of 3-mers** | |
| Ala | GCA (↓, X)<br>GCC (↓, X) | - |
| Arg | **AGA (↑, ↑)**<br>**AGG (↑, ↑)** | **AGA (↑, ↑)**<br>**AGG (↑, ↑)**<br>CGA (↓, ↑) |
| Asn | AAC (↑, ↓)<br>**AAT (↓, ↓)** | AAC (↑, ↓)<br>**AAT (↓, ↓)** |
| Asp | - | GAC (↓, X) |
| Glu | **GAA (↑, ↑)** | **GAA (↑, ↑)** |
| Gly | GGA (↑, X) | GGA (↑, X) |
| His | CAC (↑, ↓) | - |
| Ile | - | ATC (↑, ↑) |
| Leu | CTC (↑, ↑) | CTC (↑, X) |
| Lys | AAA (↑, ↑)<br>AAG (↑, ↑) | - |
| Pro | CCA (↑, ↓) | CCA (↓, ↓)<br>CCG (↓, ↓)<br>CCC (↑, ↓) |
| Ser | TCA (↑, ↓) | AGC (↑, ↓) |
| Thr | ACG (↑, ↓) | ACC (↓, ↓) |

**Confirmed Pairs Group 3 (very strong).** This group includes six hyperthermophiles: one bacterium, *T. ruber*, and five archaea: *P. furiosus*, *T. litoralis*, *T. adornatum*, *P. pacificus*, and *P. chitonophagus*. All four steps of the analysis consistently showed remarkable genomic signature similarities between the bacterium and at least one of the archaea.

3-mer deviation analysis: As shown in Figure 15, the 3-mer frequency profiles for all six organisms in this group exhibited similar patterns of over- and under-representation from the dataset average, with notable similarities between the bacterium and the five archaea.

Environment-relevant 3-mers identification: Feature importance analysis revealed a significant overlap in environment-relevant 3-mers across these taxonomically diverse organisms, which is shown in Figure 15.

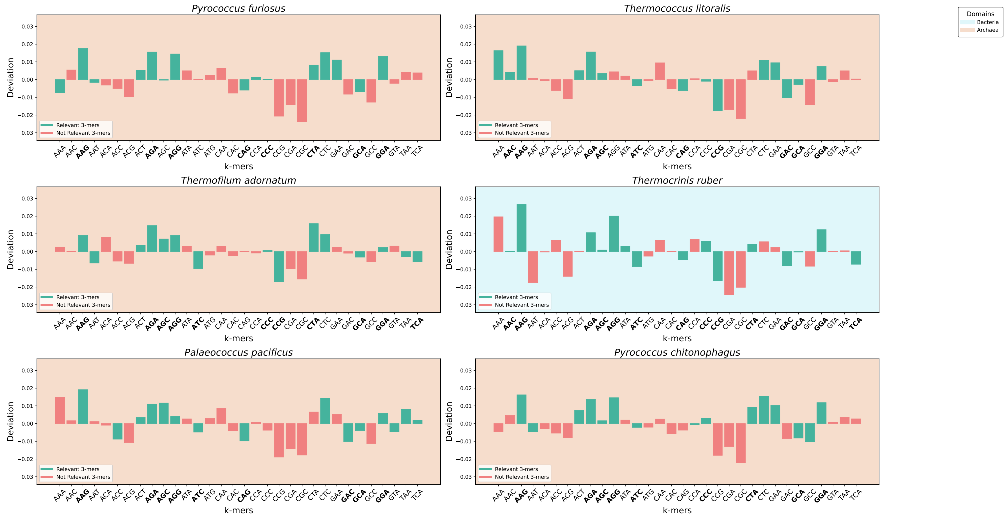


*Figure 15:* 3-mer usage bias analysis of the Confirmed Pairs Group 3. The blue panel shows the bacterium, and the orange panels show the archaea. The bars show the deviation of each 3-mer relative to the average of that 3-mer in the *Temperature dataset*. Green bars indicate 3-mers identified as relevant to the environment-type classification for that species, whereas red bars represent 3-mers that did not influence that classification. Only canonical 3-mers are considered in this analysis. Shared environment-relevant 3-mers with similar over- and under-representation are indicated in boldface.

Correlation assessment: As shown in Figure 16, the similarity in 3-mer profiles was confirmed by Spearman's rank correlation coefficients (mean correlation 0.79, range $0.76 - 0.81$, all with $p$-value $< 10^{-5}$) between the 3-mer counts of the bacterium and each of the five archaea.

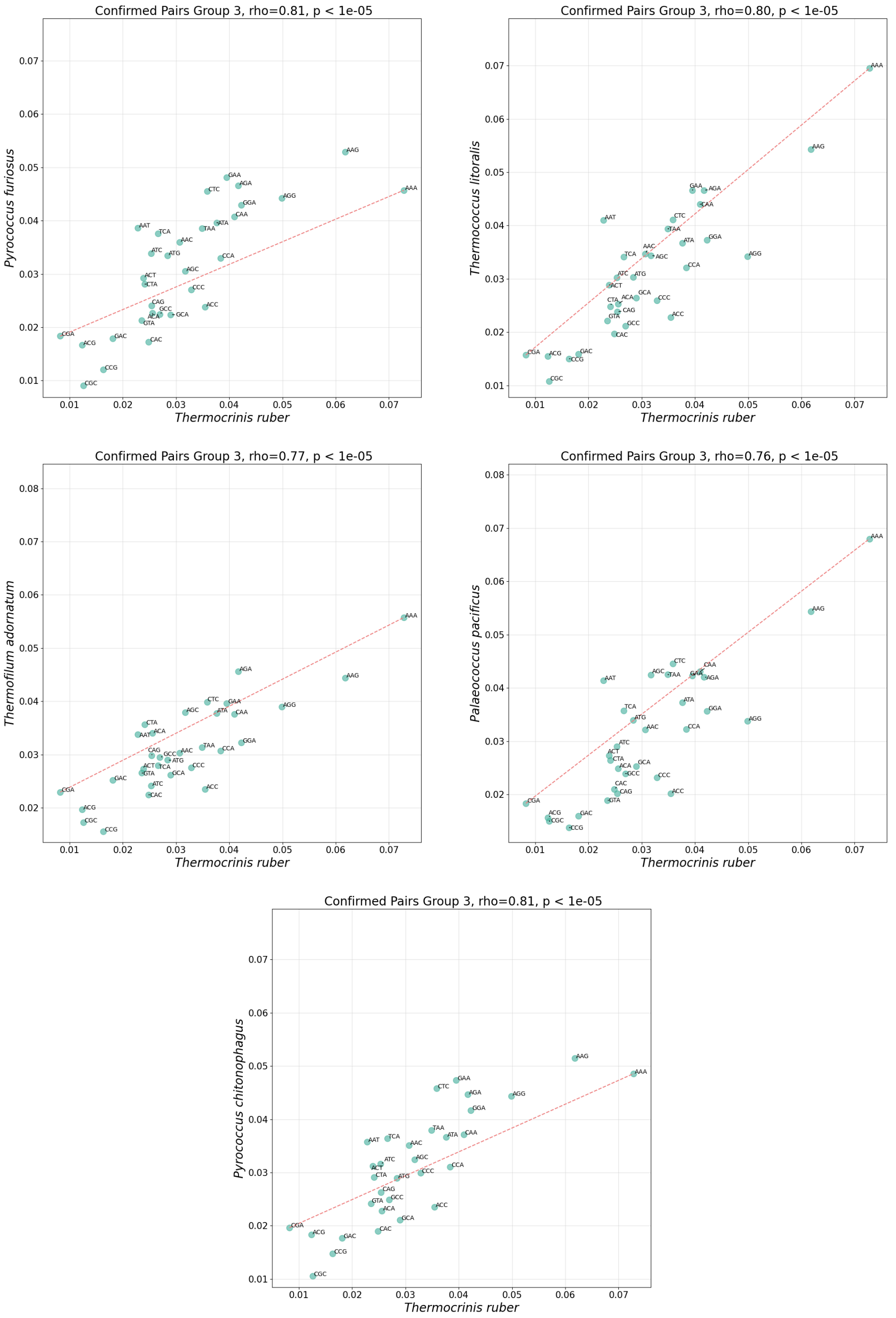


*Figure 16:* Correlation analysis of 3-mer usage between the pairs in Confirmed Pairs Group 3. Each point represents a specific 3-mer, with its normalized frequency in *T. ruber* plotted on the $x$-axis and its frequency in the five archaea on the $y$-axis. The analysis reveals a strong positive Spearman's rank correlation coefficient, with $p$-value $< 10^{-5}$ for all pairs.

Comparison with literature: As shown in Table 19, there was a moderate agreement between our findings and previously reported over- and under-representation patterns in the biology literature.

*Table 19:* Over- and under-representation of the relevant 3-mers, found by our method to be collectively associated with genomic signatures of extremophiles found in the Confirmed Pairs Group 3. The symbol ↑ (↓) indicates over-representation (under-representation) of a 3-mer/codon. Matched arrows, e.g., (↑, ↑ ref) indicate that both our method and literature agree in their finding. Mismatched arrows indicate disagreement. In cases where the letter "X" is found in place of a secondary arrow indicates a lack of supporting literature. The 3-mers with complete matching patterns are highlighted in bold.

| **Domain** | Bacteria | Archaea | | | | |
|---|---|---|---|---|---|---|
| **Species** | *Thermocrinis ruber* | *Pyrococcus furiosus* | *Thermofilum adornatum* | *Paleococcus pacificus* | *Pyrococcus chitonophagus* | *Thermococcus litoralis* |
| **Temperature Label** | Hyper-thermophiles | Hyper-thermophiles | Hyper-thermophiles | Hyper-thermophiles | Hyper-thermophiles | Hyper-thermophiles |
| **Corresponding Amino Acid** | **Representation of 3-mers** | | | | | |
| Ala | GCA (↓, *X*) | GCA (↓, *X*) | GCA (↑, *X*) | | GCA (↓, *X*)<br>GCC (↓, *X*) | GCA (↓, *X*) |
| Arg | **AGA (↑, ↑)**<br>**AGG (↑, ↑)** | **AGA (↑, ↑)**<br>**AGG (↑, ↑)** | **AGA (↑, ↑)**<br>**AGG (↑, ↑)** | **AGA (↑, ↑)**<br>**AGG (↑, ↑)** | **AGA (↑, ↑)**<br>**AGG (↑, ↑)** | **AGA (↑, ↑)** |
| Asn | AAC (↑, ↓) | AAT (↓, ↓) | AAT (↓, ↓) | | AAT (↓, ↓) | AAC (↑, ↓) |
| Asp | GAC (↓, *X*) | | | GAC (↓, *X*) | | GAC (↓, *X*) |
| Gln | **CAG (↓, ↓)** | **CAG (↓, ↓)** | | **CAG (↓, ↓)** | | **CAG (↓, ↓)** |
| Glu | | GAA (↑, ↑) | | | GAA (↑, ↑) | GAA (↑, ↑) |
| Gly | GGA (↑, *X*) | GGA (↑, *X*) | GGA (↓, *X*) | GGA (↑, *X*)<br>GCA (↓, *X*) | GGA (↑, *X*) | GGA (↑, *X*) |
| His | | | | | | |
| Ile | ATA (↑, ↑)<br>ATC (↓, ↑) | | ATC (↓, ↑) | ATC (↓, ↑) | ATC (↓, ↑) | ATC (↓, ↑) |
| Leu | **CTA (↑, ↑)** | **CTA (↑, ↑)**<br>CTC (↑, ↑) | **CTA (↑, ↑)**<br>CTC (↑, ↑) | CTC (↑, ↑) | **CTA (↑, ↑)**<br>CTC (↑, ↑) | CTC (↑, ↑) |
| Lys | **AAG (↑, ↑)** | **AAG (↑, ↑)**<br>AAA (↓, ↑) | **AAG (↑, ↑)** | **AAG (↑, ↑)** | **AAG (↑, ↑)** | **AAG (↑, ↑)**<br>AAA (↑, ↑) |
| Pro | CCC (↑, ↓)<br>**CCG (↓, ↓)** | CCA (↑, ↓) | CCC (↑, ↓)<br>**CCG (↓, ↓)** | | CCC (↑, ↓)<br>CCA (↓, ↓) | CCC (↓, ↓)<br>**CCG (↓, ↓)** |
| Ser | AGC (↑, ↓)<br>TCA (↓, ↓) | AGC (↓, ↓) | AGC (↑, ↓)<br>TCA (↑, ↓) | AGC (↑, ↓)<br>TCA (↑, ↓) | AGC (↑, ↓) | |
| STOP | | TAA (↑, *X*) | TAA (↑, *X*) | | | |
| Thr | | ACT (↑, ↓) | ACT (↑, ↓) | ACT (↑, ↓) | ACT (↑, ↓)<br>ACC (↓, ↓) | ACT (↑, ↓) |
| Val | | | | GTA (↓, ↑) | | |

**Confirmed Pairs Group 4 (strong).** This group includes one thermophilic bacterium, *P. elfii* and two mesophilic archaea, *M. paludis* and *M. vacuolata*. Three of four steps of the analysis consistently showed genomic signature similarities between the bacteria and at least one of the archaea, with the fourth indicating a moderate level of agreement with existing biological literature.

3-mer deviation analysis: As illustrated Figure 17 in the 3-mer frequency profiles for all three organisms in this group exhibited similar patterns of deviation from the dataset average.

Environment-relevant 3-mers identification: Feature importance analysis revealed an overlap in environment-relevant 3-mers across these organisms, which is shown in Figure 17.

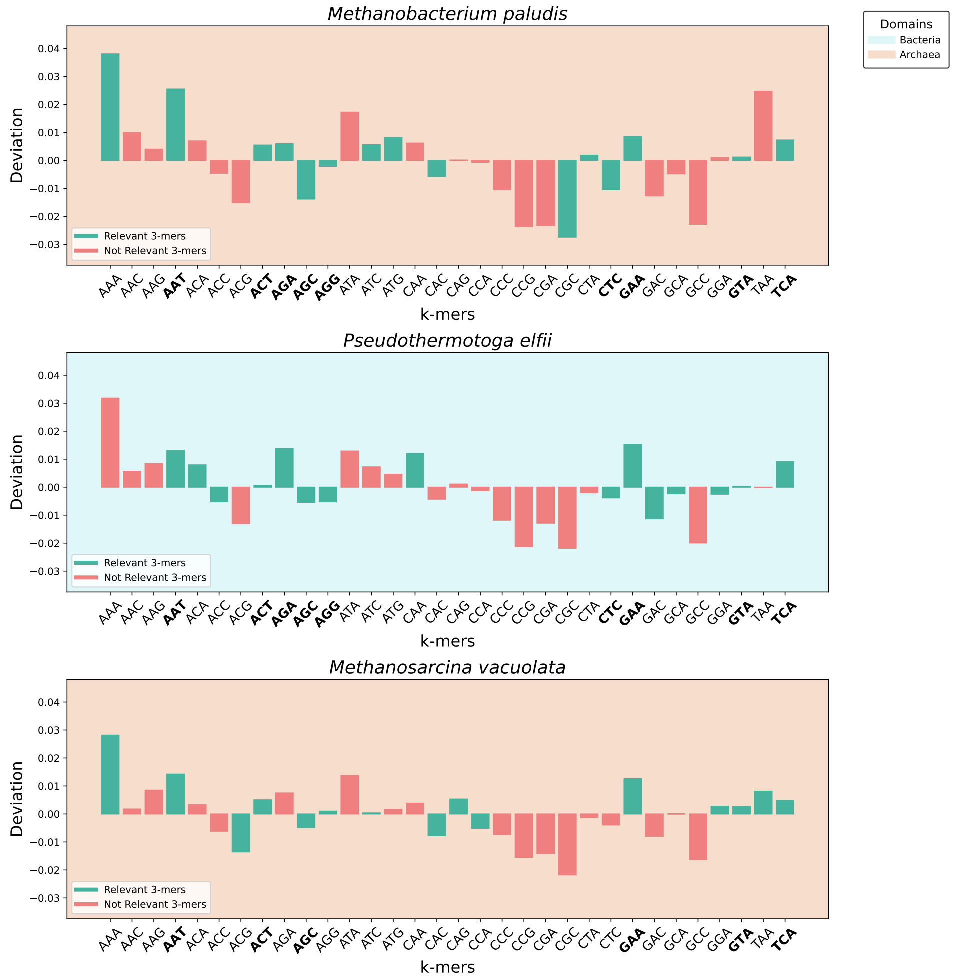


*Figure 17:* 3-mer usage bias analysis of the Confirmed Pairs Group 3. The blue panel shows the bacterium, and the orange panels show the archaea. The bars show the deviation of each 3-mer relative to the average of that 3-mer in the *Temperature dataset*. Green bars indicate 3-mers identified as relevant to the environment-type classification for that species, whereas red bars represent 3-mers that did not influence that classification. Only canonical 3-mers are considered in this analysis. Shared environment-relevant 3-mers with similar over- and under-representation are indicated in boldface.

Correlation assessment: As shown in Figure 18, the similarity in 3-mer profiles was quantitatively confirmed by high Spearman's rank correlation coefficients (mean correlation = 0.945, all with $p$-value $< 10^{-5}$) between the 3-mer counts of the bacterium and each of the two archaea.

Comparison with literature: As shown in Table 20, compared with Groups 1, 2 and 3, less agreement was found between the 3-mer biases identified by our method and biology literature findings.

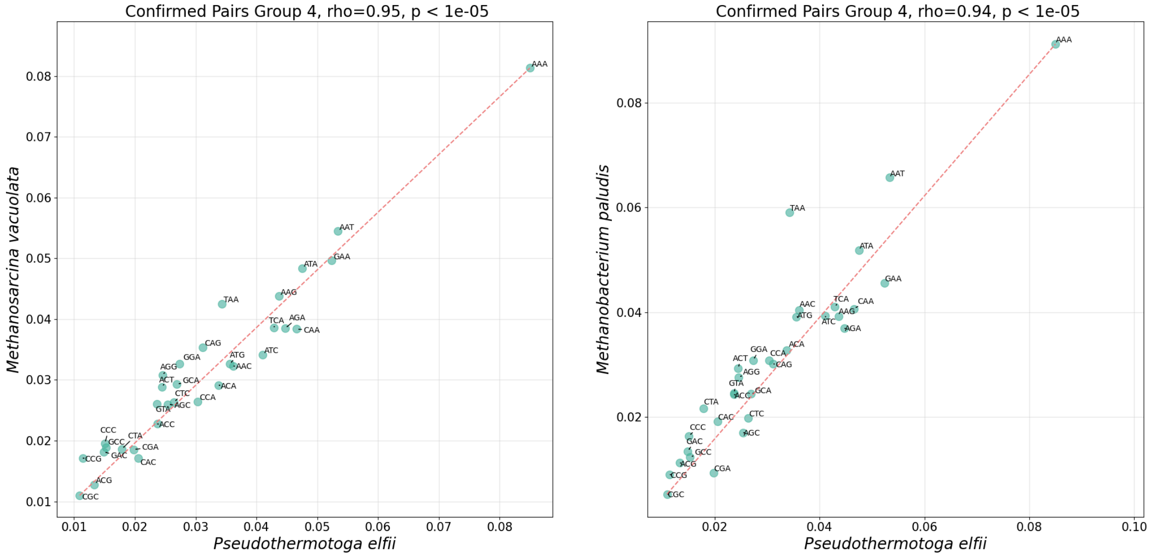


*Figure 18:* Correlation analysis of 3-mer usage between the pairs from the Confirmed Pairs Group 4. Each point represents a specific 3-mer, with its normalized frequency in *P. elfi* plotted on the $x$-axis and its frequency in the archaea on the $y$-axis. The analysis reveals a strong positive Spearman's rank correlation coefficient, with $p$-value $< 10^{-5}$ for all pairs.

*Table 20:* Over- and under-representation of the relevant 3-mers, found by our method to be collectively associated with genomic signatures of extremophiles found in Confirmed Pairs Group 4. The symbol ↑ (↓) indicates over-representation (under-representation) of a 3-mer/codon. Matched arrows, e.g., (↑, ↑) indicate that both our method and literature agree in their finding. Mismatched arrows indicate disagreement. In cases where the letter “X" is found in place of a secondary arrow indicates a lack of supporting literature. The 3-mers with complete matching patterns across species are highlighted in bold.

| **Domain** | Bacteria | Archaea | |
|---|---|---|---|
| **Species** | *Pseudothermotoga elfii* | *Methanobacterium paludis* | *Methanosarcina vacuolata* |
| **Temperature Label** | Thermophiles | Mesophiles | Mesophiles |
| **Corresponding Amino Acid** | **Representation of 3-mers** | | |
| Ala | GCA (↓, X) | - | - |
| Arg | AGG (↓,↑) | AGA (↑,↓)<br>AGG (↓,↓)<br>CGC (↓,↑) | AGG (↑,↓) |
| Asn | AAT (↑, X) | AAT (↑,↑) | AAT (↑,↑) |
| Asp | GAC (↓, X) | - | - |
| Gln | CAA (↑,↓) | - | CAG (↑,↑) |
| Glu | GAA (↑,↑) | GAA (↑, X) | GAA (↑, X) |
| Gly | GGA (↓,↑) | - | GGA (↑,↓) |
| His | - | CAC (↓,↑) | CAC (↓,↑) |
| Ile | - | ATC (↑,↓) | ATC (↑,↓) |
| Leu | CTC (↓,↑) | CTC (↓, X) | - |
| Lys | - | AAA (↑, X)<br>CTA (↑,↓) | AAA (↑, X) |
| Met | - | ATG (↑,↑) | - |
| Pro | - | - | CCA (↓,↓) |
| Ser | **AGC (↓,↓)**<br>TCA (↑,↓) | **AGC (↓,↓)**<br>TCA (↑,↑) | **AGC (↓,↓)**<br>TCA (↑,↑) |
| STOP | - | - | TAA (↑,↑) |
| Thr | ACA (↑, -)<br>ACC (↓,↓)<br>ACT (↓,↓) | ACT (↑,↓) | ACT (↑, X)<br>ACG (↓,↓) |
| Val | GTA (↑,↑) | GTA (↑, X) | GTA (↑, X) |

**Confirmed Pairs Group 5 (strong).** This group includes one mesophilic bacterium, *R. indicoceani*, five mesophilic archaea, *M. marisnigri*, *M. bourgensis*, *M. mesophila*, *M. horonobensis*, and *M. taiwanensis*, and one thermophilic archaeon *M. thermophilus*. Three out of four steps of the analysis revealed remarkable genomic signature similarities between the bacteria and at least one of the archaea, but less agreement was found with biological literature.

3-mer deviation analysis: As shown in Figure 19, the 3-mer frequency profiles for all three organisms in this group showed similar patterns of deviation from the dataset average.

Environment-relevant 3-mer identification: Feature importance analysis revealed an overlap in environment-relevant 3-mers across these organisms, which is shown in Figure 19.

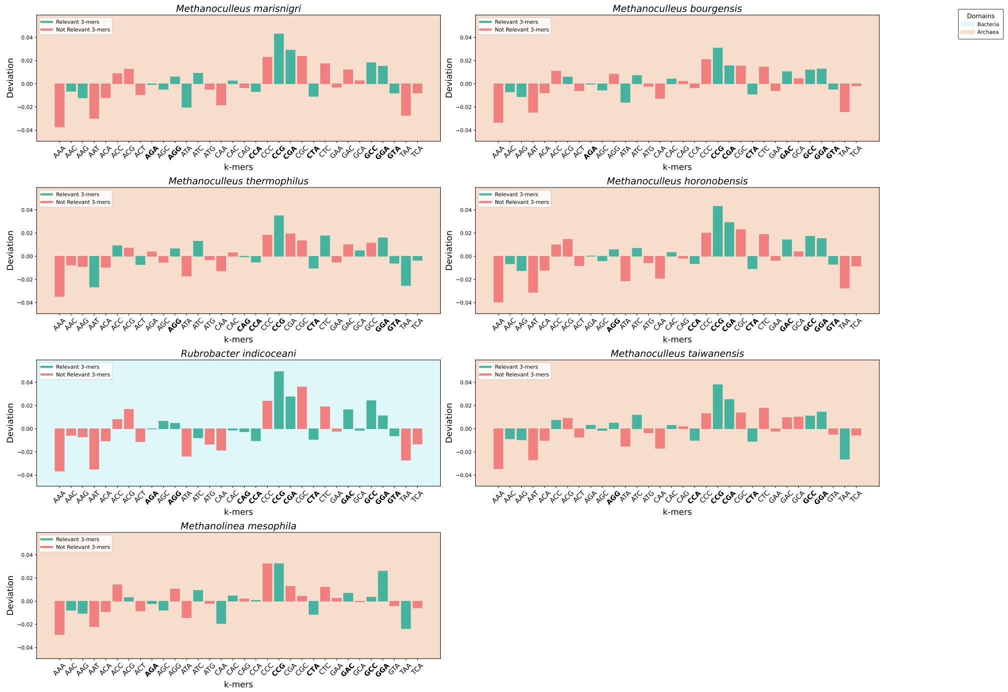


*Figure 19:* 3-mer usage bias analysis of the Confirmed Pairs Group 35. The blue panel shows the bacterium, and the orange panels show the archaea. The bars show the deviation of each 3-mer relative to the average of that 3-mer in the *Temperature dataset*. Green bars indicate 3-mers identified as relevant to the environment-type classification for that species, whereas red bars represent 3-mers that did not influence that classification. Only canonical 3-mers are considered in this analysis. Shared environment-relevant 3-mers with similar over- and under-representation are indicated in boldface.

Correlation assessment: As shown in Figure 20, the similarity in 3-mer profiles was quantitatively confirmed by high Spearman's rank correlation coefficients (all with $p$-value $< 10^{-5}$) between the 3-mer counts of the bacterium and each of the archaea.

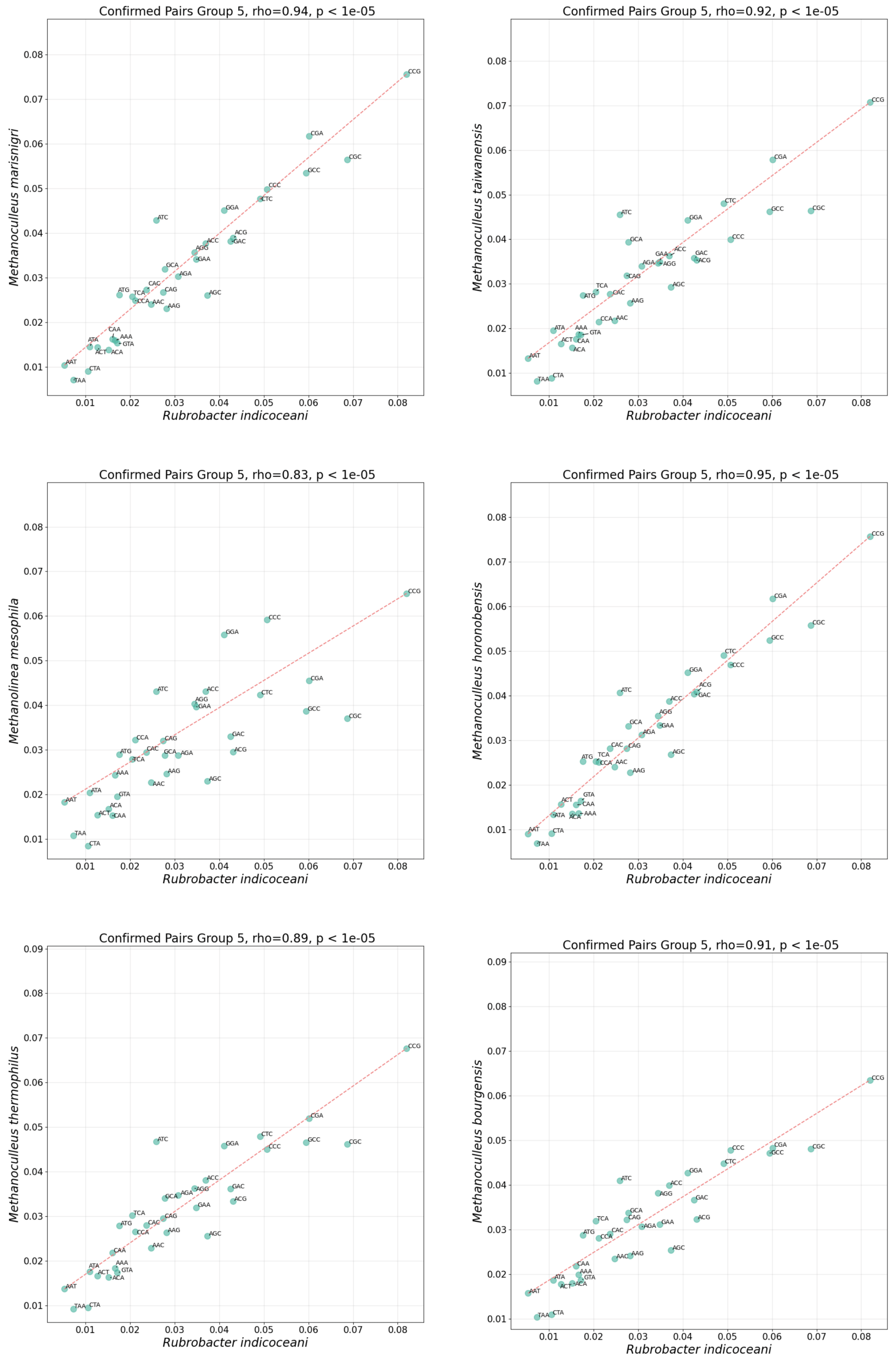


*Figure 20:* Correlation analysis of 3-mer usage between the pairs in Confirmed Pairs Group 5. Each point represents a specific 3-mer, with its normalized frequency in *R. indicoceani* plotted on the $x$-axis and its frequency in the six archaea on the $y$-axis. The analysis reveals a strong positive Spearman's rank correlation coefficient, with $p$-value $< 10^{-5}$ for all pairs.

Comparison with literature: As shown in Table 21, similar to Group 4, less agreement was found between 3-mer biases identified by our method and biology literature findings.

*Table 21:* Over- and under-representation of the relevant 3-mers, found by our method to be collectively associated with genomic signatures found in Confirmed Pairs Group 5. The symbol ↑ (↓) indicates over-representation (under-representation) of a 3-mer/codon. Matched arrows, e.g., (↑, ↑) indicate that both our method and literature agree in their finding. Mismatched arrows indicate disagreement. In cases where the letter "X" is found in place of a secondary arrow indicates a lack of supporting literature. The 3-mers with complete matching patterns across species are highlighted in bold.

| **Domain** | Bacteria | Archaea | | | | | |
|---|---|---|---|---|---|---|---|
| **Species** | *Rubrobacter indicoceani* | *Methanoculleus marisnigri* | *M. bourgensis* | *M. thermophilus* | *M. horonobensis* | *M. taiwanensis* | *Methanolinea mesophila* |
| **Temperature Label** | Mesophiles | Mesophiles | Mesophiles | Thermophiles | Mesophiles | Mesophiles | Mesophiles |
| **Corresponding Amino Acid** | **Representation of 3-mers** | | | | | | |
| Ala | AGA (↑, ↓)<br>GCA (↓,↓)<br>GCC (↑, X) | GCC (↑, X) | GCC (↑, X) | GCA (↑, X) | GCC (↑, X) | GCC (↑, X) | GCC (↑, X) |
| Arg | AGG (↑,↓)<br>**CGA (↑, ↑)** | AGA (↓,↓)<br>AGG (↑, ↓) | AGA (↓,↓)<br>**CGA (↑, ↑)** | AGG (↑, ↑) | AGG (↑,↓)<br>**CGA (↑, ↑)** | AGA (↓,↓)<br>AGG (↑, ↓)<br>**CGA (↑, ↑)** | AGA (↓,↓) |
| Asn | - | AAC (↓, ↑) | AAC (↓, ↑) | AAT (↓, X) | AAC (↓, ↑) | AAC (↓, ↑) | AAC (↓, ↑) |
| Asp | GAC (↑, X) | - | GAC (↑, X) | - | GAC (↑, X) | - | GAC (↑, X) |
| Gln | CAG (↓, ↑) | - | - | CAG (↓, ↑) | - | - | CAA (↓, X) |
| Glu | - | - | - | - | - | - | - |
| Gly | GGA (↑,↓) | GGA (↑,↓) | GGA (↑,↓) | GGA (↑, ↑) | GGA (↑,↓) | GGA (↑,↓) | GGA (↑,↓) |
| His | CAC (↓, ↑) | CAC (↑, ↑) | CAC (↑, ↑) | - | CAC (↑, ↑) | CAC (↑, ↑) | CAC (↑, ↑) |
| Ile | ATC (↓,↓) | ATA (↓, ↓)<br>ATC (↑, ↓) | ATA (↓, ↓)<br>ATC (↑, ↓) | ATC (↑, ↑) | ATC (↑,↓) | ATC (↑,↓) | ATC (↑,↓) |
| Leu | **CTA (↓,↓)** | **CTA (↓,↓)** | **CTA (↓,↓)** | CTA (↓, ↑)<br>CTC (↑, ↑) | **CTA (↓,↓)** | **CTA (↓,↓)** | **CTA (↓,↓)** |
| Lys | - | AAG (↓, X) | AAG (↓, X) | - | AAG (↓, X) | AAG (↓, X) | AAG (↓, X) |
| Met | - | - | - | - | - | - | - |
| Pro | **CCA (↓,↓)**<br>CCG (↑,↓) | **CCA (↓,↓)**<br>CCG (↑,↓) | CCG (↑,↓) | CCA (↓, ↑)<br>CCG (↑, X) | **CCA (↓,↓)**<br>CCG (↑,↓) | **CCA (↓,↓)**<br>CCG (↑,↓) | CCA (↑,↓)<br>CCG (↑,↓) |
| Ser | AGC (↑,↓) | AGC (↓,↓) | AGC (↓,↓) | TCA (↓,↓) | AGC (↓,↓) | AGC (↓,↓) | AGC (↓,↓) |
| STOP | - | - | - | TAA (↓, X) | - | TAA (↓, ↑) | TAA (↓, ↑) |
| Thr | - | - | ACG (↑,↓) | ACC (↑,↓)<br>ACT (↓,↓) | - | ACC (↑, X)<br>ACG (↑,↓) | - |
| Val | GTA (↓, X) | GTA (↓, X) | - | GTA (↓, ↑) | GTA (↓, X) | - | - |

Table 22 shows the references to the literature used for this study.

*Table 22:* Codon usage biases across extremophile groups as characterized in the literature. The table summarizes the various codon usage biases, considering related amino acid abundance patterns associated with prokaryotes across pH and temperature-adapted extremophilic categories. Each observation is followed by [ref], linking to the paper providing the rationale for this pattern.

| Codon Usage Biases | Acidophiles | Alkaliphiles | Psychrophiles | Mesophiles | Thermophiles | Hyperthermophiles |
|---|---|---|---|---|---|---|
| **Increased** | CAA [20] | AAC [20] | GCA [14] | ATG [30] | AGG [45] | AGG [45] |
| | GGA[20] | AAT [20] | CAA [4] | CAA [20] | AGA [45] | AGA [45] |
| | CCA [20] | GAC [20] | GAA [37] | AAT [20] | GGA [10] | CTC [45] |
| | CGC [20] | ATG [14] | CAG [20] | CTC [45] | CTA [45] | |
| | AGC [33] | CGC [7] | CCA [15] | GTA [10] | | |
| | TCA [33] | TAT [7] | CCC [15] | | | |
| **Decreased** | AGG [20] | CAA [20] | AGG [35] | ATC [41] | ACT [15] | CCT [15] |
| | CTC [20] | GAG [20] | CAC [35] | ACG [7] | ACG [15] | CCC [15] |
| | TCA [20] | | CTC [37] | TCC [7] | | CCA [15] |
| | AAA[14] | ACA [7] | | | CCG [15] | |
| | CCA [7] | | GGC [7] | | | |
| | AGT [7] | | | TCG [7] | | |

### I Co-occurrences details

This section provides details of the geographical co-occurrences of Groups 1 to 4, in Table 23, Table 24, Table 25, Table 26, respectively. No co-occurrence was found for Group 5.

*Table 23:* Description of co-occurrence of species found in Group 1. The following table describes the location data for each of the co-occurrences between the two species in Group 1, including the species name for the mapped 16s rRNA reads, sample IDs for the relevant Microbe Atlas Project entry, coordinate information (Latitude, Longitude), and notable environmental observations.

| **Attribute** | **Description** |
|---|---|
| Mapped Read 1 ID | *Thermoanaerobacterium thermosaccharolyticum* |
| Mapped Read 2 ID | *Caldisphaera lagunensis* |
| Sample ID | SRS5070100 |
| Environment Descriptor | Washburn Hot Springs [26] |
| Coordinates | 44.376° N, 110.69° W |
| Notable Environmental Observations | Described by McKay et al. [26] as an "ideal ancient habitat," displaying high levels of sulphate, sulphide, methane, hydrogen, carbon dioxide, and ammonium sulphate. |

*Table 24:* Description of co-occurrence of species in Group 2. The following table describes the location data for each of the co-occurrences between the species in Group 2, including the species name for the mapped 16s rRNA reads, sample IDs for the relevant Microbe Atlas Project entry, coordinate information (Latitude, Longitude), and notable environmental observations.

| **Attribute** | **Description** | |
|---|---|---|
| Mapped Read 1 ID | *Thermotoga petrophila* | |
| Mapped Read 2 ID | *Geoglobus acetivorans* | |
| Sample IDs | SRS5544491 | SRS730398 |
| Environment Descriptor | Brothers Volcano [34] | Juan de Fuca Ridge [19] |
| Coordinates | 34.86208959° S, 179.0573881° E | 47.753500° N, 127.763883° W |
| Notable Environmental Observations | Hydrothermally active submarine volcano | Deep biofilm community demonstrating iron redox chemistry |

*Table 25:* Description of co-occurrence of species found in Group 3. The following table describes the location data for each of the co-occurrences between species found in Group 3, including the species name for the mapped 16s rRNA reads, sample IDs for the relevant Microbe Atlas Project entry, coordinate information (Latitude, Longitude), and notable environmental observations. Habitats where an occurrence was identified for a particular species are specified, displaying unique co-occurrences where location data was available (duplicates are removed).

| **Sample ID** | **Environment** | **Coordinates** | **Notable Observations** | **Co-occurrences by species** | | |
|---|---|---|---|---|---|---|
| | | | | *T. ruber* | *T. litoralis* | *T. adornatum* |
| SRS7008575 | Norris-Mammoth Corridor, YNP, USA | 44.754° N, 110.7257° W | Hot spring water | ✓ | | ✓ |
| SRS4347131 | Culex Basin, YNP, USA | 44.5775° N, 110.7896° W | Hot spring sediment | ✓ | | ✓ |
| SRS6018221 | Geyser Creek Basin, YNP, USA | 44.6904° N, 110.7291° W | Hot spring sediment | ✓ | | ✓ |
| SRS1971309 | Joseph's Coat, YNP, USA | 44.376° N, 110.69° W | Hot spring sediment | ✓ | | ✓ |
| SRS2354974 | Mono Lake, USA | 37.97° N, 119.07° W | High salinity salt lake | ✓ | | ✓ |
| SRS3206925 | Guaymas Basin, Gulf of California | 27.0114° N, 110.5931° W | Hydrothermal vent sediment | ✓ | ✓ | |
| ERS1370018 | Mid-Cayman Rise, Cayman Trough | 18°32'49.9"N, 81°43'05.2"W | Deep sea hydrothermal vent | ✓ | ✓ | |
| SRS5070100 | Washburn Springs, YNP, USA | 44.376° N, 110.69° W | Extremophilic microbial mat | ✓ | | ✓ |
| SRS6608865 | Yellowstone Lake, YNP, USA | 44.5106° N, 110.3566° W | Filamentous streamer microbial communities | ✓ | | ✓ |
| SRS5544488 | Brothers Volcano, Pacific Ocean | 34.8611° S, 179.0576° E | Hydrothermal vent sediment | ✓ | | ✓ |
| ERS1372490 | Manus Basin, Papua New Guinea | 3°47'59.7"S, 152°06'03.1"E | Marine basaltic hydrothermal vent biome | ✓ | ✓ | |

*Table 26:* Description of co-occurrence of species found in Group 4. The following table describes the location data for each of the co-occurrences between the species in Group 4, including the species name for the mapped 16s rRNA reads, sample IDs for the relevant Microbe Atlas Project entry, coordinate information (Latitude, Longitude), and notable environmental observations.

| **Attribute** | **Description** | | |
|---|---|---|---|
| Mapped Read 1 ID | *Pseudothermotoga elfii* | | |
| Mapped Read 2 ID | *Methanobacterium paludis* | | |
| Mapped Read 3 ID | *Methanosarcina vacuolata* | | |
| Sample ID | DRS114146 | SRS586146 | SRS836040 |
| Environment Descriptor | Bioreactor, Tokyo, Japan [21] | Shengli Oil Field, China [31] | Norman, Oklahoma, USA |
| Coordinates | 35.7127° N, 139.7620° E | 37.54° N, 118.33° E | 35.2200° N, 97.4400° W |
| Notable Environmental Observations | Microbiome from a “thermophilic methanogenic biocathode” implying high temperature and methane conditions [21] | Anaerobic, mesophilic culture; involved in “methanogenic degradation of hydrocarbons” [31] | Landfill leachate exposed to “$CH_4$ and $CO_2$ gases” and complex soluble chemical compounds [40] |
